# Evidence for a Large-Scale Brain System Supporting Allostasis and Interoception in Humans

**DOI:** 10.1101/098970

**Authors:** Ian R. Kleckner, Jiahe Zhang, Alexandra Touroutoglou, Lorena Chanes, Chenjie Xia, W. Kyle Simmons, Karen S. Quigley, Bradford C. Dickerson, Lisa Feldman Barrett

**Affiliations:** Department of Psychology, Northeastern University, Boston, MA.; Department of Neurology, Massachusetts General Hospital and Harvard Medical School; Athinoula A. Martinos Center for Biomedical Imaging; Psychiatric Neuroimaging Division, Department of Psychiatry, Massachusetts General Hospital and Harvard Medical School, Charlestown, MA; Frontotemporal Disorders Unit, Department of Neurology, Massachusetts General Hospital and Harvard Medical School, Charlestown, MA; Laureate Institute for Brain Research, Tulsa, OK; School of Community Medicine, The University of Tulsa, Tulsa, OK; Edith Nourse Rogers Memorial VA Hospital, Bedford, MA

## Abstract

Large-scale intrinsic brain systems have been identified for exteroceptive senses (e.g., sight, hearing, touch). We introduce an analogous system for representing sensations from within the body, called interoception, and demonstrate its relation to regulating peripheral systems in the body, called allostasis. Employing the recently introduced Embodied Predictive Interoception Coding (EPIC) model, we used tract-tracing studies of macaque monkeys, followed by two intrinsic functional magnetic resonance imaging samples (*N* = 280 and *N* = 270) to evaluate the existence of an intrinsic allostatic/interoceptive system in the human brain. Another sample (*N* = 41) allowed us to evaluate the convergent validity of the hypothesized allostatic/interoceptive system by showing that individuals with stronger connectivity between system hubs performed better on an implicit index of interoceptive ability related to autonomic fluctuations. Implications include novel insights for the brain’s functional architecture, dissolving the artificial boundary between mind and body, and unifying mental and physical illness.

---

The brain contains intrinsic systems for processing exteroceptive sensory inputs from the world, such as vision, audition, and proprioception/touch (e.g., ^1^). Accumulating evidence indicates that these systems work via the principles of predictive coding (e.g., ^2–7^), where sensations are anticipated and then corrected by sensory inputs from the world. The brain, as a generative system, models the world by *predicting*, rather than reacting to, sensory inputs. Predictions guide action and perception by continually constructing possible representations of the immediate future based on their prior probabilities relative to the present context^8,9^. We and others have recently began studying the hypothesis that ascending sensory inputs from the organs and systems within the body’s *internal milieu* are similarly anticipated and represented (i.e., autonomic visceral and vascular function, neuroendocrine fluctuations, and neuroimmune function)^10−15^. These sensations are referred to as *interoception*^16–18^. Engineering studies of neural design^19^, along with physiological evidence^20^, indicate that the brain continually anticipates the body’s energy needs in an efficient manner and prepares to meet those needs before they arise (e.g., movements to cool the body’s temperature before it gets too hot). This process is called *allostasis*^19−21^. Allostasis is not a condition or state of the body – it is the process by which the brain efficiently maintains energy regulation in the body. Allostasis is defined in terms of prediction, and recent theories propose that the prediction of interoceptive signals is necessary for successful allostasis (e.g., ^10,15,22–24^). Thus, in addition to the ascending pathways and brain regions important for interoception (e.g., ^16,17,25 26^), recent theoretical discussions (e.g.,^11^) have proposed the existence of a distributed intrinsic allostatic/interoceptive system in the brain (analogous to the exteroceptive systems). A full investigation of the predictive nature of an allostatic/interoceptive brain system requires multiple studies under various conditions. Here, we begin this line of research by identifying the anatomical and functional substrates for a unified allostatic/interoceptive system in the human brain and reporting an association between connectivity within this system and individual differences in interoceptive-related behavior during allostatically-relevant events.

In this paper, we first review tract-tracing studies of non-human animals that provide the anatomical substrate for our hypothesis that the brain contains a unified, intrinsic system for allostasis and interoception. Next, we present evidence of this hypothesized system in humans using functional connectivity analyses on three samples of task-independent (i.e., “resting state”) functional magnetic resonance imaging (fMRI) data (also called “intrinsic” connectivity). We then present brain-behavior evidence to validate the hypothesized allostatic/interoceptive system by using an implicit measure of interoception during an allostatically challenging task. Finally, we summarize empirical evidence to show that this allostatic/interoceptive system is a *domain-general* system that supports a wide range of psychological functions including interoception, emotion, memory, reward, cognitive control, etc.^27,28^. That is, whatever else this system might be doing – remembering, directing attention, etc., – they are also predictively regulating the body’s physiological systems in the service of allostasis to achieve those functions^22^.

The novelty of our work is the synthesis of anatomical and functional brain studies that together evidence a single brain system – comprised of the salience and default mode networks – that supports not just allostasis but a wide range of psychological functions (emotion, pain, memory, decision-making, etc.) that can all be explained by their reliance on allostasis. To our knowledge, this evidence and our simple yet powerful explanation has not been presented despite the fact that many functional imaging studies show that the salience and default mode networks support a wide range of psychological functions (i.e., they are domain general; e.g., ^29^; for review, see ^27,28^). Our paper provides the groundwork for a theoretical and empirical framework for making sense of these findings in an anatomically principled way. Our key hypotheses and results are summarized in Table 1.

**Table 1.** Summary of this study’s hypotheses, predictions or questions, and results.

| Embodied Predictive Interoception Coding (EPIC) Hypothesis | Experimental Prediction | Result in the Current Study |
| --- | --- | --- |
| Interoception and visceromotor control are part of a unified brain system that supports allostasis (Fig 1) | Primary interoceptive cortex (e.g., dorsal mid/posterior insula) is anatomically and functionally connected to agranular and dysgranular visceromotor hubs of the cortex (e.g., sgACC, pACC, aMCC) | • The interoceptive and visceromotor hubs are anatomically connected in monkeys (Table 2)<br>• The interoception and visceromotor hubs are functionally connected in humans (Fig. 2, Table S1)<br>• Coordinates for human hubs are show in Table 3 |
|  | The allostatic/interoceptive system also includes subcortical and brainstem visceromotor regions. | • Previously established subcortical and brainstem visceromotor regions (e.g., hypothalamus, periaqueductal gray) are part of the unified system for allostasis/interoception (Fig. 4, Fig. S6) |
|  | The allostatic/interoceptive brain system will contain limbic cortices. | • The allostatic/interoceptive system comprises two established large scale brain networks that contain the majority of limbic cortices: the salience network and the default mode networks (Fig. 3, Fig. S3) |
|  | Connectivity in the allostatic/interoceptive system is related to an implicit, performance measure of interoception in humans | • The correspondence between sympathetic arousal (electrodermal activity) and experienced arousal during an allostatically challenging task is related to functional connectivity within the allostatic/interoceptive system in humans (Fig. S8) |
| The allostatic/interoceptive system is domain-general. | The allostatic/interoceptive system sits that the core of the brain’s computational architecture. | • Many hubs of the allostatic/interoceptive system have been previously identified as members of the “rich club” that are the most densely connected within the brain and therefore help constitute the brain’s “neural backbone” for coordinating neural synchrony(Fig. 3, Table S4) |
|  | Brain activity and connectivity in the allostatic/interoceptive system is associated with a variety of psychological functions | Both the default mode network and the salience network support a variety of mental phenomena across major psychological domains (e.g., cognition, emotion, perception, and action). (Fig. 5) |
| Other hypotheses, such as the computational dynamics of the proposed allostatic/interoceptive network are beyond the scope of this study. ACC = anterior cingulate cortex; aMCC = anterior midcingulate cortex; dmIns = dorsal mid insula; dpIns = dorsal posterior insula; pACC = pregenual anterior cingulate cortex; sgACC = subgenual anterior cingulate cortex. |  |  |

## Anatomical evidence supporting the proposed allostatic/interoceptive system

Over three decades of tract-tracing studies of the macaque monkey brain clearly demonstrate an anatomical substrate for the proposed flow of the brain’s prediction and prediction error signals. Specifically, anatomical studies indicate a flow of information within the laminar gradients of these cortical regions according to the structural model of corticocortical connections developed by Barbas colleagues (^30^; for a review, see ^31^). In addition, the structural model of corticocortical connections have been seamlessly integrated with a predictive coding framewor^11,12^. Unlike other models of information flow that work in specific regions of cortex, the structural model successfully predicts information flow in frontal, temporal, parietal, and occipital cortices^32–36^. Accordingly, prediction signals flow from regions with less laminar development (e.g., agranular regions) to regions with greater laminar development (e.g., granular regions), whereas prediction error signals flow in the other direction. In our recently developed theory of interoception, called the Embodied Predictive Interoception Coding (EPIC) model^11^, we integrated Friston’s active inference approach to predictive coding^37–39^ with Barbas’s structural model to hypothesize that less-differentiated agranular and dysgranular visceromotor cortices in the cingulate cortex and anterior insula initiate visceromotor predictions through their cascading connections to the hypothalamus, the periaqueductal gray (PAG), and other brainstem nuclei known to control the body’s internal milieu^40–43^ (also see ^31^; red pathways in Fig 1); simultaneously, the cingulate cortex and anterior insula send the anticipated sensory consequences of those visceromotor actions (i.e., interoceptive predictions) to the more granular primary interoceptive cortex in the dorsal mid to posterior insula (dmIns/dpIns^17,44,45^; blue solid pathways; Fig 1). Using this logic, we identified a key set of cortical regions with visceromotor connections that should form the basis of our unified system for interoception and allostasis (we also included one subcortical region, the dorsal amygdala (dAmy), in this analysis due to the role of the central nucleus in visceromotor regulation; for details, see endnote 1). This evidence is summarized in Table 2. As predicted by our EPIC model, most of the key visceromotor regions in the proposed interoceptive system do, in fact, have monosynaptic, bidirectional connections to primary interoceptive cortex, reinforcing the hypothesis that they directly exchange interoceptive prediction and prediction error signals. We also confirmed that these visceromotor cortical regions indeed monosynaptically project to the subcortical and brainstem regions that control the internal milieu (i.e., the autonomic nervous system, immune system, and neuroendocrine system), such as the hypothalamus, PAG, parabrachial nucleus (PBN), ventral striatum, and nucleus of the solitary tract (NTS) (Table 2, right column).

**Fig 1.**
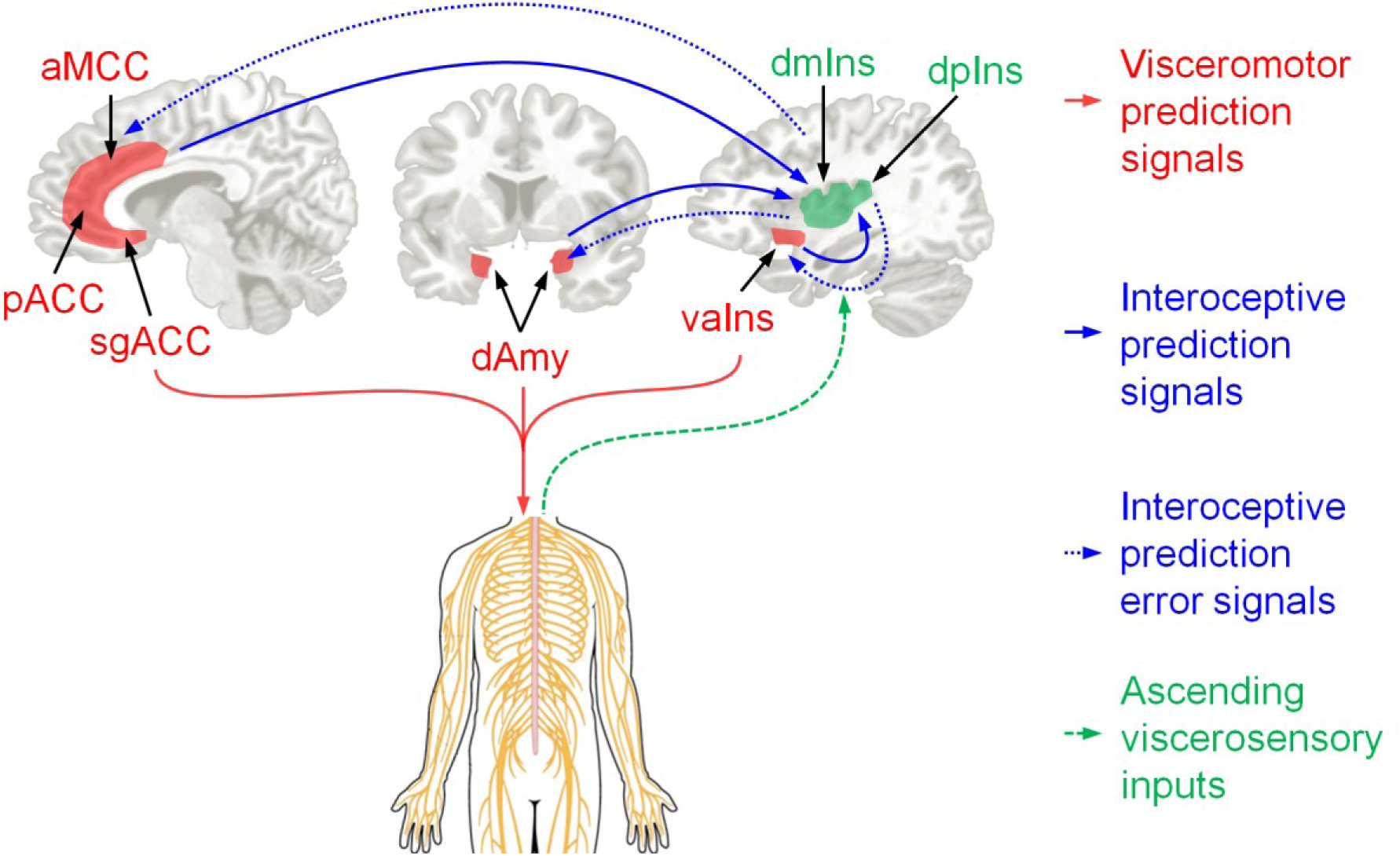
We identified key visceromotor cortical regions (in red) that provide cortical control the body’s internal milieu, including the anterior mid cingulate cortex (aMCC; also called dorsal anterior cingulate cortex (dACC), e.g., ^40,41^), pregenual anterior cingulate cortex (pACC), subgenual anterior cingulate cortex (sgACC; for a review of the cingulate, see ^162^), and the ventral anterior insula (vaIns; also called agranular insula^42,169^ or posterior orbitofrontal cortex^180^); these regions have a less-developed laminar structure (i.e., they are agranular or dysgranular^31,162^). We also included the dorsal amygdala because it contains the central nucleus which is also involved in visceromotor control (for a review, see ^139^). Primary interoceptive cortex spans the dorsal mid insula (dmIns) to the dorsal posterior insula (dpIns)^16^ along a dysgranular to granular^181^ gradient (green regions). Barrett & Simmons (2015) summarized preliminary tract-tracing evidence, supporting the EPIC model^11^, that allostasis and interoception are maintained within an integrated system involving limbic cortices (in red) that initiate visceromotor directions to the hypothalamus and brainstem nuclei (e.g., periaqueductal gray, parabrachial nucleus, nucleus of the solitary tract; citations in Table 2) to regulate the autonomic, neuroendocrine, and immune systems (red paths). These visceromotor control regions (less developed laminar organization) also send anticipated sensory consequences of visceromotor changes (as interoceptive prediction signals) to primary interoceptive cortex (more-developed laminar organization; solid blue paths). The incoming sensory inputs from the internal milieu of the body are carried along the vagus nerve and small diameter C and Aδ fibers (dashed green paths) to primary interoceptive cortex in the dorsal sector of the mid to posterior insula (for a review, see ^16^); comparisons between prediction signals and ascending sensory input results in interoceptive prediction error. Current interoceptive predictions can be updated by passing prediction error signals to visceromotor regions (dashed blue paths); prediction errors are learning signals and also adjust subsequent predictions. (For simplicity, ascending feedback to visceromotor regions is not shown). aMCC = anterior midcingulate cortex; dAmy = dorsal amygdala; dmIns = dorsal mid insula; dpIns = dorsal posterior insula; pACC = pregenual anterior cingulate cortex; sgACC = subgenual anterior cingulate cortex; vaIns = ventral anterior insula.

**Table 2.** Summary of tract-tracing study results in non-human animals, demonstrating anatomical connections between cortical visceromotor and primary interoceptive sensory regions, as well as between cortical and non-cortical visceromotor regions.

|  | Primary Interoceptive Cortex | Visceromotor Regions |  |  |  | Subcortical and Brainstem Visceromotor Structures |  |
| --- | --- | --- | --- | --- | --- | --- | --- |
|  | To dpIns/dmIns | To vaIns | To sgACC (BA 25) | To pACC (BA 24, 32) | To aMCC (BA 24) | To Amygdala | To other subcortical and brainstem regions<sup>a</sup> |
| **From dpIns/dmIns** | - | Case A, Fig 1<sup>141</sup> | Not evident<sup>b</sup> | Case 1, Fig 5<sup>143</sup> | Case B, Fig 3<sup>144</sup> | Case 2, Fig 3<sup>140</sup><br>Case BB-B, Fig 1<sup>59</sup> | Hypothalamus (rat)<sup>145</sup><br>PAG: not observed<sup>146</sup><br>PBN (rat)<sup>147,148</sup><br>V. Striatum<sup>149</sup><br>NTS (rat)<sup>148</sup> |
| **From vaIns<sup>c</sup>** | Case C, Fig 4<sup>141</sup><br>Case A, Fig 1<sup>144</sup> | - | Case OM20, Fig 8<sup>150</sup> | Case 1, Fig 5<sup>143</sup> | Case 2, Fig 6<sup>143</sup><br>Case A, Fig 1<sup>144</sup> | Case A, Fig 1<sup>144</sup><br>Case 103, Fig 3<sup>151</sup><br>Fig 2, Table 2<sup>152</sup> | Hypothalamus<sup>42</sup><br>PAG<sup>146</sup><br>PBN (rat)<sup>147</sup><br>V. striatum<sup>153</sup><br>NTS (rat)<sup>148</sup> |
| **From sgACC (BA 25)** | Not evident<sup>d</sup> | Case M707<sup>154</sup> | - | Case 1, Fig 5<sup>143</sup><br>Fig 2A<sup>155</sup> | Case 3, Fig 7<sup>143</sup><br>Fig 3A<sup>155</sup> | Case 103, Fig 3<sup>151</sup><br>Fig 5<sup>142</sup> | Hypothalamus<sup>140,156,157</sup><br>PAG<sup>146,157</sup><br>PBN<sup>157</sup><br>Striatum<sup>157</sup><br>NTS (rat)<sup>158,159</sup> |
| **From pACC (BA 24, 32)** | Not evident<sup>d</sup> | Case M776<sup>154</sup> | Fig 1<sup>155</sup> | - | Case 3, Fig 7<sup>143</sup><br>Fig 3A<sup>155</sup> | Case 103, Fig 3<sup>151</sup><br>Fig 5<sup>142</sup> | Hypothalamus<sup>42</sup>,<br>PAG<sup>146</sup><br>PBN (cat)<sup>160</sup><br>V. striatum (cat)<sup>160</sup><br>NTS (rat)<sup>159</sup> |
| **From aMCC (BA 24)** | Case C, Fig 4<sup>141</sup> | Case A, Fig 1<sup>141</sup> | Case 3, Fig 4<sup>161</sup> | Case 1, Fig 5<sup>143</sup><br>Fig 2A<sup>155</sup> | - | Case 103, Fig 3<sup>151</sup><br>Fig 5<sup>142</sup> | Hypothalamus<sup>42</sup><br>PAG<sup>146</sup><br>PBN: not present<sup>162</sup><br>V. striatum<sup>163</sup><br>NTS (rat)<sup>158</sup> |
| **From Amygdala** | Case C, Fig 4<sup>141</sup><br>Lateral basal nucleus; Case 5, Fig 6<sup>140</sup> | Case A, Fig 1<sup>141</sup><br>Case 4, Fig 5<sup>140</sup> | Fig 6<sup>142</sup> | Fig 13<sup>155</sup> | Fig 6<sup>142</sup> | - | Hypothalamus<sup>42</sup>,<br>PAG<sup>146</sup><br>PBN<sup>164</sup><br>V. striatum<sup>165</sup><br>NTS<sup>164</sup> |

Note. Connectivity evidence is in monkeys unless otherwise indicated (e.g., rats, cats). Some connections from dpIns/dmIns to the NTS are unclear due to ambiguity in how Saper (1982)<sup>148</sup> reported subregions of the insula.
<sup>a</sup>We did not assess for projections from subcortical and brainstem regions to cortical regions because we only wanted to determine if the cortical regions support visceromotor control.
<sup>b</sup>Connection from dpIns/dmIns to sgACC not evident in several monkey studies that have the potential to show them (e.g., <sup>144,155,166-168</sup>).
<sup>c</sup>The medial portion of the vaIns exhibits connectivity with subcortical and brainstem regions, but not the lateral portion of the vaIns<sup>42,169</sup>.
<sup>d</sup>Connection from sgACC to dpIns/dmIns and from pACC to dpIns/dmIns not evident in several monkey studies that have the potential to show them (e.g., <sup>141,154,166,167</sup>), although weak, direct connectivity is evident in a recent tractography study in humans (Ghaziri, et al., 2015<sup>170</sup>, Figure 5). Moreover, connections between sgACC, pACC, and dpIns have been observed in intrinsic functional connectivity analyses in humans (e.g., Fig. 6 of <sup>171</sup>). The discrepancy between human findings and the tract tracing studies in monkeys failing to show connectivity might reflect an expansion of Brodmann area (BA) 24 anterior and ventral to the corpus callosum in humans relative to monkeys and/or the presence of connections between BAs 25/32 and the posterior insula in humans that do not exist in monkeys (Evrard, H. personal communication, December 27, 2015).
BA = Brodmann area; aMCC = anterior midcingulate cortex; dmIns = dorsal mid insula; dpIns = dorsal posterior insula; NTS = nucleus of the solitary tract; PAG = periaqueductal gray; PBN = parabrachial nucleus; pACC = pregenual anterior cingulate cortex; sgACC = subgenual anterior cingulate cortex; V. striatum = ventral striatum.

Next, we tested for evidence of these connections in functional data from human brains. Axonal connections between neurons, both direct (monosynaptic) and indirect (e.g., disynaptic) connections, are closely reflected in intrinsic brain systems (for a review, see ^46,47^). As such, we tested for evidence of these connections in functional connectivity analyses on two samples of low-frequency Blood Oxygenation Level Dependent (BOLD) signals during task-independent (i.e., “resting state”) fMRI scans collected on human participants (discovery sample, *N* = 280, 174 female, mean age = 19.3 years, SD = 1.4 years; replication sample, *N* = 270,142 female, mean age = 22.3 years, SD = 2.1 years). We then examined the validity of these connections in a third independent sample of participants (*N* = 41, 19 female, mean age = 33.5 years, SD = 14.1 years), following which we situated these findings in the larger literature on network function.

## Results

### Cortical and amygdalar intrinsic connectivity supporting a unified allostatic/interoceptive system in humans

Our seed-based approach estimated the functional connectivity between a set of voxels of interest (i.e., the seed) and the voxels in the rest of the brain as the correlation between the low-frequency portion of their BOLD signals over time, producing a discovery map for each seed region. Starting with the anatomical regions of interest specified by the EPIC model, and verified in the anatomical literature, we selected seed regions guided by previously published functional studies. We selected two groupings of voxels in primary interoceptive cortex (dpIns and dmIns) that consistently showed increased activity during task-dependent fMRI studies of interoception (Table 3, first and second rows). We selected seed regions for cortical visceromotor regions and the dAmy using related studies (Table 3, remaining rows). As predicted, the voxels in primary interoceptive cortex and visceromotor cortices showed statistically significant intrinsic connectivity (Fig. 2; replication sample Fig. S1). The dpIns was intrinsically connected to all visceromotor areas of interest (seven two-tailed, one-sample t-tests were each significant at *p* < 10^-7^; Table S1), and dmIns was intrinsically connected to most of them (Table S1). The discovery and replication samples demonstrated high reliability for connectivity profiles of all seeds (*η^2^* mean = 0.99, SD = 0.004).

**Table 3.** Seeds used for intrinsic connectivity analyses.

| **Seed** | **Type of region predicted by EPIC model** | **Cortical Lamination** | **MNI Coordinates** |
| --- | --- | --- | --- |
| **dpIns** | Primary interoceptive cortex | Granular | 36, -32, 16<sup>172</sup> |
| **dmIns** | Primary interoceptive cortex | Dysgranular | 41, 2, 3<sup>173</sup> |
| **sgACC** | Visceromotor control | Agranular | 2, 14, -6<sup>174</sup> |
| **pACC** | Visceromotor control | Agranular | 13, 44, 0<sup>172</sup> |
| **aMCC** | Visceromotor control | Agranular | 9, 22, 33<sup>175</sup> |
| **mvaIns** | Visceromotor control | Agranular | 30, 16, -14<sup>176</sup> |
| **lvaIns** | Sensory integration | Agranular | 44, 6, -15<sup>175</sup> |
| **dAmy** | Visceromotor control | N/A | 27, 3, -12<sup>177</sup> |
| *Note:* All seeds are in the right hemisphere. Evidence for cortical lamination comes from Vogt (2005)<sup>41,178,179</sup>.<br>Each anatomical region of interest was represented by one 4-mm-radius seed except for the ventral anterior insula (vaIns), which required a medial and a lateral seed (mvaIns and lvaIns, respectively) to capture the previously-established functional distinction between the medial visceromotor network (containing mvaIns) and the orbital sensory integration network (containing lvaIns) in the orbitofrontal cortex<sup>169</sup>.<br>aMCC = anterior midcingulate cortex; dAmy = dorsal amygdala; dmIns = dorsal mid insula; dpIns = dorsal posterior insula; lvaIns = lateral ventral anterior insula; mvaIns = medial ventral anterior insula; pACC = pregenual anterior cingulate cortex; sgACC = subgenual anterior cingulate cortex. |  |  |  |

**Fig. 2.**
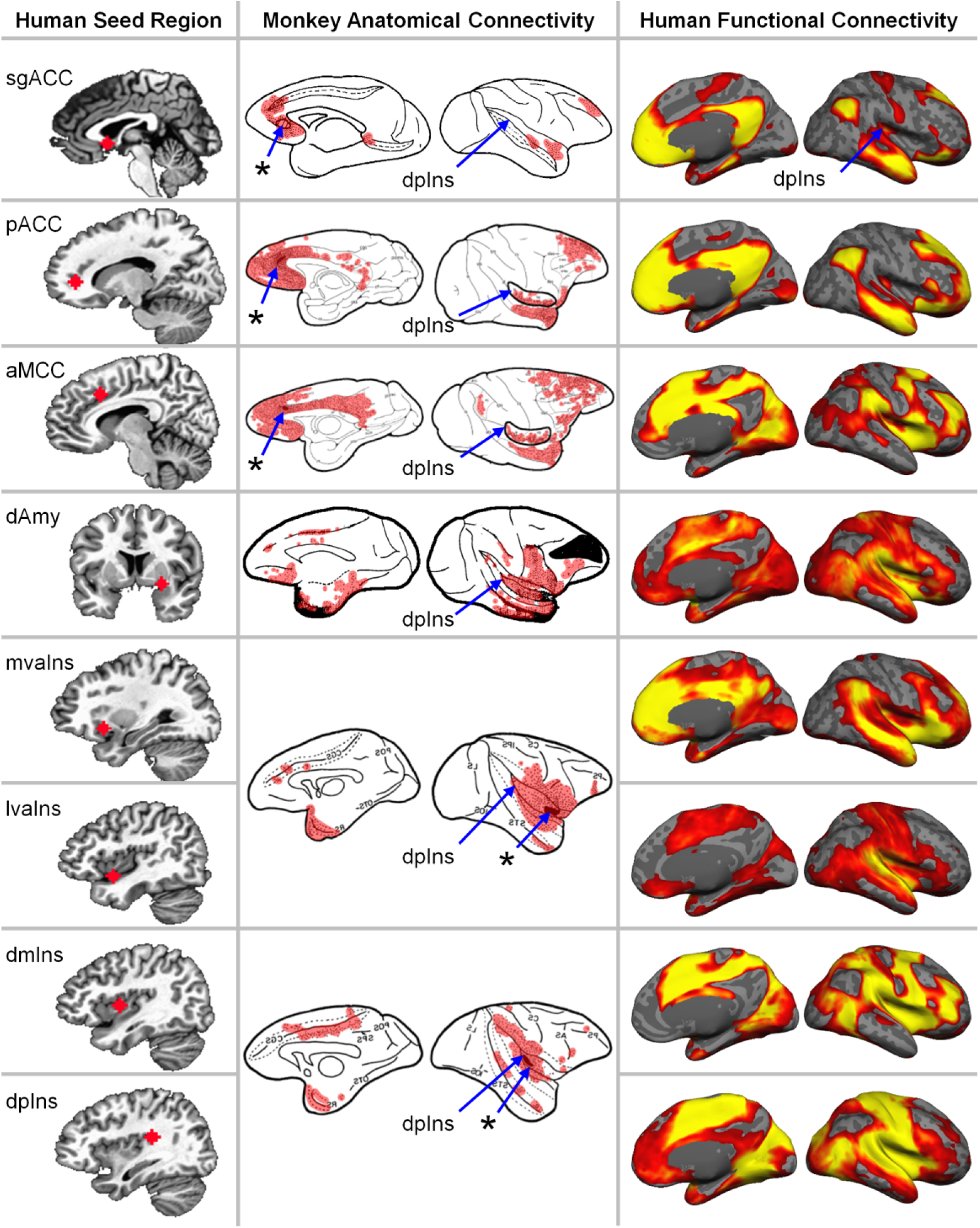
Eight regions (“seeds”) used to estimate the unified allostasis/interoceptive system connecting the cortical and amygdalar visceromotor regions and primary interoceptive regions. The left column shows the “seed” region for each discovery map on a human brain template. The middle column summarizes the anatomical connectivity derived from anterograde and/or retrograde tracers injected in macaque brains at a location homologous to the human seed (asterisks with blue arrows). The right column shows the human intrinsic connectivity discovery maps depicting all voxels whose time course is correlated with the seed’s (ranging from*p*< 10^-5^ in red to *p* < 10^-40^ in yellow, uncorrected, *N* = 280). To avoid Type I and Type II errors, which are enhanced with the use of stringent statistical thresholds^182^, we opted to separate signal from random noise using replication, according to the mathematics of classical measurement theory^183^. These results replicated in a second sample, *N* = 270 participants, indicating that they are reliable and cannot be attributed to random error (Fig. S1). Functional connectivity to the entire amygdala and other subcortical regions are shown in Fig. 4. Tract tracing figures were adapted with permission as follows: subgenual anterior cingulate cortex (sgACC) via retrograde tracers in Fig 1 of Vogt & Pandya (1987)^155^, pregenual ACC (pACC) via retrograde tracers in Fig 5 of Morecraft, et al. (2012)^143^, anterior midcingulate cortex (aMCC) via retrograde tracers in Fig 7 of Morecraft, et al. (2012)^143^, dorsal amygdala (dAmy) via retrograde tracers in Fig 3 of Aggleton, et al. (1980)^151^, medial ventral anterior insula (mvaIns) and lateral ventral anterior insula (lvaIns) via anterograde tracers in Fig 1 of Mesulam & Mufson (1982)^144^, dorsal mid insula (dmIns) and dorsal posterior insula (dpIns) via anterograde tracers in Fig 3 of Mesulam & Mufson (1982)^144^. The monkey anatomical connectivity figures were colored red to visualize results and some were mirrored to match the orientation of the human brain maps. The figures from Morecraft, et al. (2012)^143^ were adapted to show the insula in its lateral view.

Next, we computed *η*^2^ for all pairs of maps to determine their spatial similarity^48^ (mean = 0.56, SD = 0.17), and then performed K-means clustering of the *η*^2^ similarity matrix to determine the configuration of the system. Results indicated that the allostatic/interoceptive system is composed of two intrinsic networks connected in a set of overlapping regions (Fig. 3; replication sample, Fig. S2). The spatial topography of one network resembled an intrinsic network commonly known as the *default mode network* (Fig. S3 and Fig. S4; for a review, see ^49^). The second network resembled an intrinsic network commonly known as the *salience network* (Fig.S3 and Fig. S4; e.g., ^50,51^), the cingulo-opercular network^52^, or the ventral attention network^53^. Resemblance was confirmed quantitatively by comparing the percent overlap in our observed networks to reconstructions of the default mode and salience networks reported in Yeo, et al.^54^ (Table S2). Other cortical regions within the interoceptive system shown in Fig. 3 (e.g., dorsomedial prefrontal cortex, middle frontal gyrus), not listed in Table 2, support visceromotor control via direct anatomical projections to the hypothalamus and PAG (Table S3), supporting our hypothesis that this system plays a fundamental role in visceromotor control and allostasis.

**Fig. 3.**
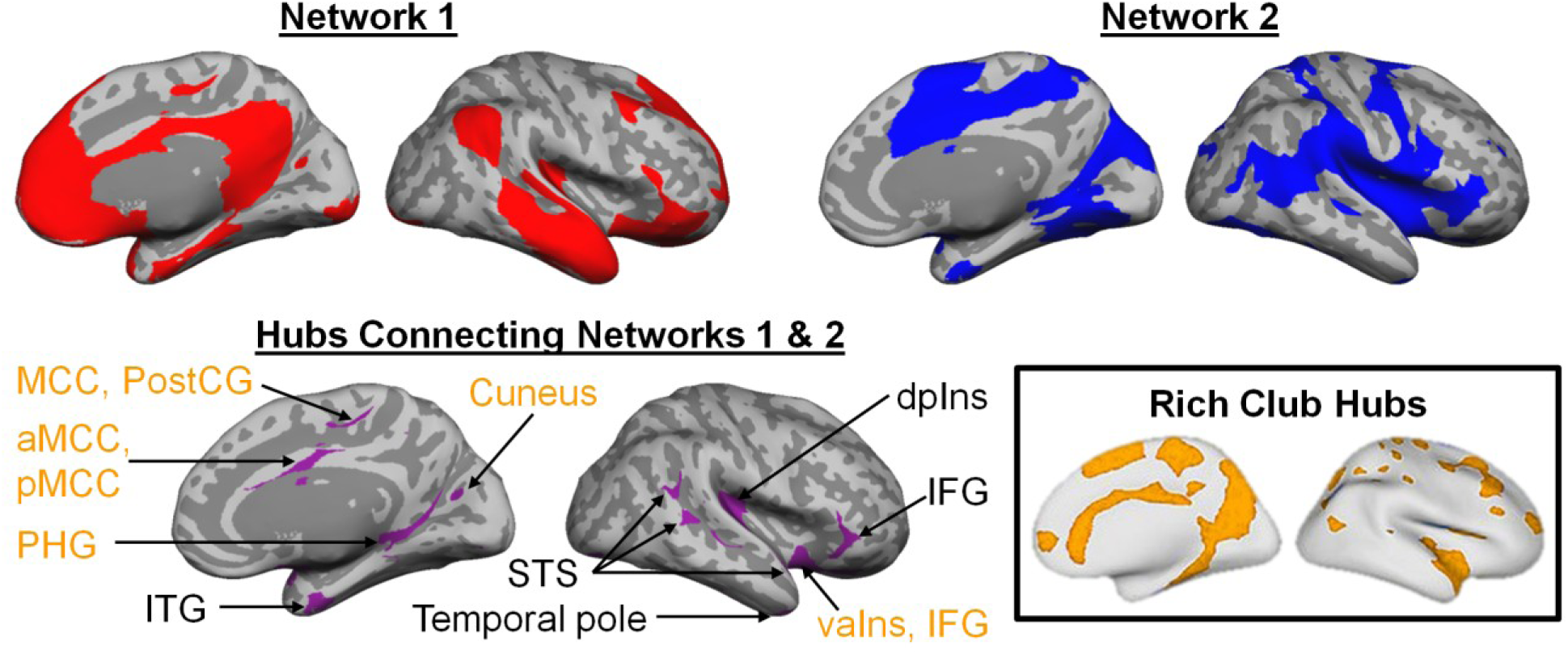
The unified allostatic/interoceptive system is composed of two large-scale intrinsic networks (shown in red and blue) that share several hubs (shown in purple; for coordinates, see Table S4). Hubs belonging to the “rich club” are shown in yellow. Rich club hubs figure adapted with permission from van den Heuval & Sporns (2013)^84^. All maps result from the sample of 280 participants binarized at p < 10^-5^ uncorrected from a one-sample two-tailed t-test. These results replicated in a second sample, *N* = 270 participants, indicating that they are reliable and cannot be attributed to random error (Fig. S2). aMCC = anterior midcingulate cortex; dAmy = dorsal amygdala; dpIns = dorsal posterior insula; dmIns = dorsal mid insula; IFG = inferior frontal gyrus; ITG = inferior temporal gyrus; lvaIns = lateral ventral anterior insula; MCC = midcingulate cortex; mvaIns = medial ventral anterior insula; pACC = pregenual anterior cingulate cortex; PHG = parahippocampal gyrus; pMCC = posterior midcingulate cortex; PostCG = postcentral gyrus; sgACC = subgenual anterior cingulate cortex; STS = superior temporal sulcus.

### Subcortical, hippocampal, brainstem, and cerebellar connectivity supporting a unified allostatic/interoceptive system in humans

Using a similar analysis strategy, we assessed the intrinsic connectivity between the cortical and dorsal amygdalar seeds of interest and the thalamus, hypothalamus, cerebellum, the entire amygdala, hippocampus, ventral striatum, PAG, PBN, and NTS. The observed functional connections with these cortical and amygdalar seeds, which regulate energy balance, strongly suggest that the proposed allostatic/interoceptive system itself also regulates energy balance (see Supplementary Results for details). All results replicated in our independent sample (*N* = 270; Fig. S5, *η*^2^ mean = 0.98, SD = 0.008). Fig. 4 illustrates the connectivity between default mode and salience networks and the non-cortical targets in the discovery sample. Fig. S6 shows connectivity between the individual cortical and amygdalar seed regions listed in Table 2. We also observed specificity in the proposed allostasis/interoception system: non-visceromotor brain regions that are unimportant to interoception and allostasis, such as the superior parietal lobule (Fig. S7), did not show functional connectivity to the subcortical regions of interest.

**Fig. 4.**
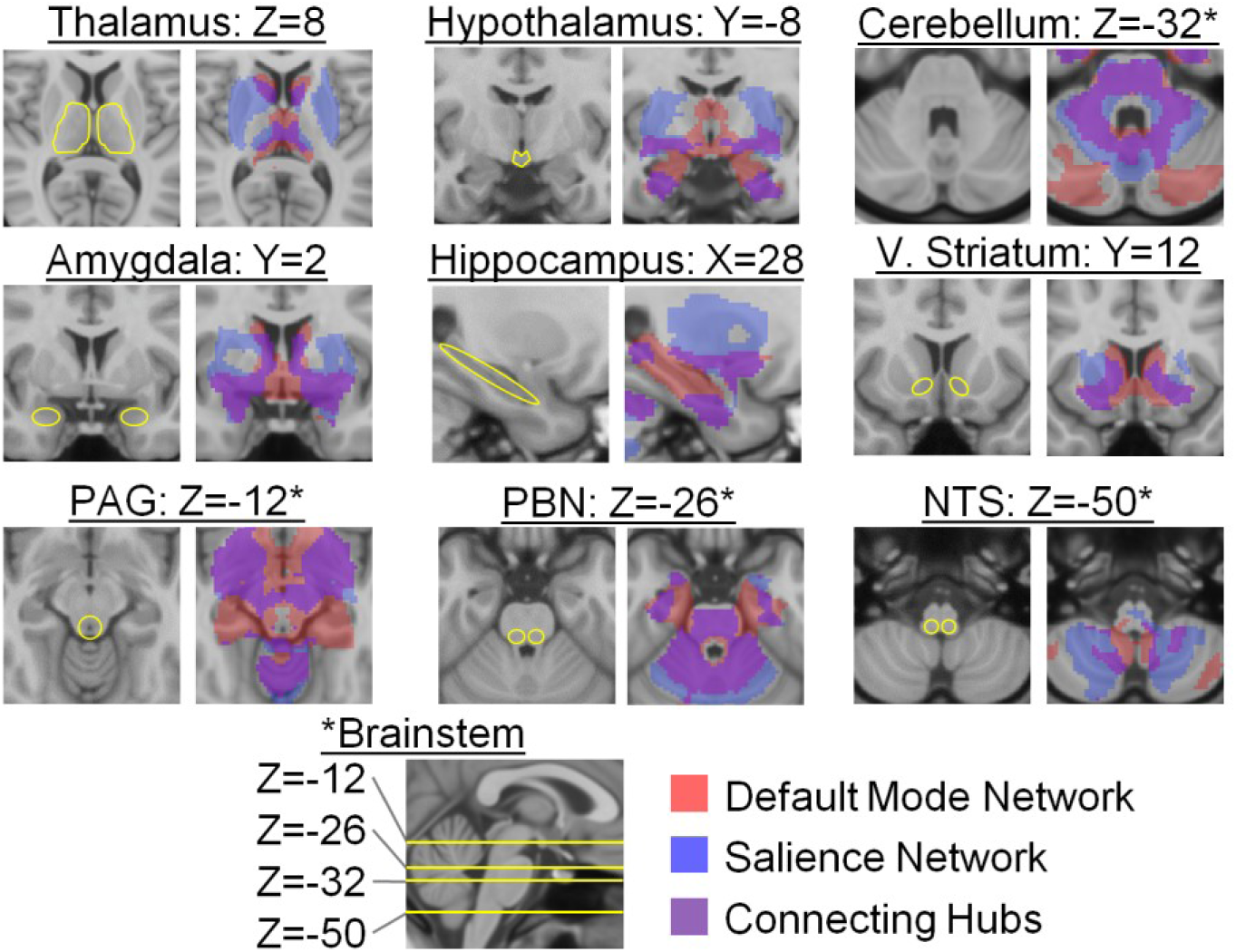
Subcortical connectivity of the two integrated intrinsic networks within the allostatic/interoceptive system (*N* = 280; *p* < 0.05 uncorrected). These results replicated in a second sample of *N* = 270 (Fig. S5). PAG = periaqueductal gray; PBN = parabrachial nucleus; V. Striatum = ventral striatum; NTS = nucleus of the solitary tract.

The cortical hubs of the allostatic/interoceptive system also overlapped in their connectivity to non-cortical regions involved in allostasis (purple in Fig. 4), including the dAmy,the hypothalamus, the PBN, and two thalamic nuclei – the VMpo and both the medial and lateral sectors of the mediodorsal nucleus (MD, which shares strong reciprocal connections with medial and orbital sectors of the frontal cortex, the lateral sector of the amygdala, and other parts of the basal forebrain; for a review, see ^55^). Additionally, the connector hubs also shared projections in the cerebellum and hippocampus (see Fig. 4).

Taken together, our intrinsic connectivity analyses failed to confirm only five monosynaptic connections (8%) that were predicted from non-human tract-tracing studies: hypothalamus-dAmy, hypothalamus-dpIns, PAG-dAmy, PAG-medial ventral anterior insula (mvaIns), and NTS-subgenual anterior cingulate cortex (sgACC). This is approximately what we would expect by chance; however, there are several factors that might account for why these predicted connections did not materialize in our discovery and replication samples. First, all discrepancies involved the sgACC, PAG, or hypothalamus, whose BOLD data exhibit poor signal to noise ratio due to their small size and their proximity to white matter or pulsating ventricles and arteries^56^. Second, individual differences in anatomical structure can make inter-subject alignment challenging, particularly in 3-T imaging of the brainstem where clear landmarks are not always available. Of the connections that did not replicate, one involved the anterior insula; there is some disagreement in the macaque anatomical literature as to the exact location of the anterior insula (e.g., ^44,57–59^), which might help explain any lack of correspondence between intrinsic and tract-tracing findings that we observed.

### Validating the functions of the allostatic/interoceptive system in humans

The allostatic/interoceptive system reported in Fig. 3 replicated in the validation sample (*η*^2^ mean = 0.84, SD = 0.05 compared with discovery sample cortical maps; *η^2^* mean = 0.76, SD = 0.07 compared with discovery sample subcortical maps). These *η*^2^ values are respectable and demonstrate adequate reliability of the system according to conventional psychometric theory, although the lower *η*^2^ values are likely due to the smaller sample size which magnifies the effects of poor signal-to-noise ratio in subcortical regions. Convergent validity for the proposed allostatic/interoceptive system was demonstrated in that individuals with stronger functional connectivity within the system also reported greater arousal while viewing images that evoked greater sympathetic nervous system activity. Participants viewed ninety evocative photos known to induce a range of autonomic nervous system changes and corresponding feelings of arousal^60^, as well as changes in BOLD activity within these regions^61,62^. We predicted, and found, that individuals showing stronger intrinsic connectivity within the allostatic/interoceptive system (specifically, connectivity between dpIns and anterior midcingulate cortex (aMCC)) also demonstrated a stronger concordance between objective and subjective measures of bodily arousal while viewing allostatically relevant images (*p* = 0.003; see Fig. S8; see Supplementary Results for details).

There were three reasons for demonstrating the convergent validity of the proposed allostatic/interoceptive system using this task. First, there is a decades-old body of research indicating that interoception enables the subjective experience of arousal (^63^; e.g., ^64,65^). Thus, the amount of joint information shared by an objective, psychophysiological measure of visceromotor change (skin conductance) and the subjective experience of arousal (self-report ratings) is an implicit, behavioral measure of interoceptive ability. Indeed, individuals with more accurate interoceptive ability exhibit a stronger correspondence between subjective arousal and physiological arousal in response to similar evocative photos^66^. Second, explicit reports of interoceptive performance on heartbeat detection tasks (e.g., ^67–69^) require synthesizing and comparing information from other systems, including the somatosensory system^70^, frontoparietal control systems, and, for heartbeat detection, the auditory system – adding an additional level of difficulty and complexity – in tasks that are sometimes too hard (yielding floor effects) or have questionable validity^69^.

At this juncture, it is tempting to ask if the unified allostatic/interoceptive system is specific to allostasis and interoception. From our perspective, this is the wrong question to be asking. The last two decades of neuroscience research have brought us to the brink of a paradigm shift in understanding the workings of the brain, setting the stage to revolutionize brain: mind mapping. Neuroscience research is increasingly acknowledging that brain networks have a one (network) to many (function) mappings ^27–29,71–73^. Our findings contribute to this discussion: a brain system that is fundamental to allostasis and interoception is not unique to those functions, but instead is also important for a wide range of psychological phenomena that span cognitive, emotional, and perceptual domains (Fig. 5.). This finding is not a failure of reverse inference. It suggests a functional feature of how the brain works.

**Fig. 5.**
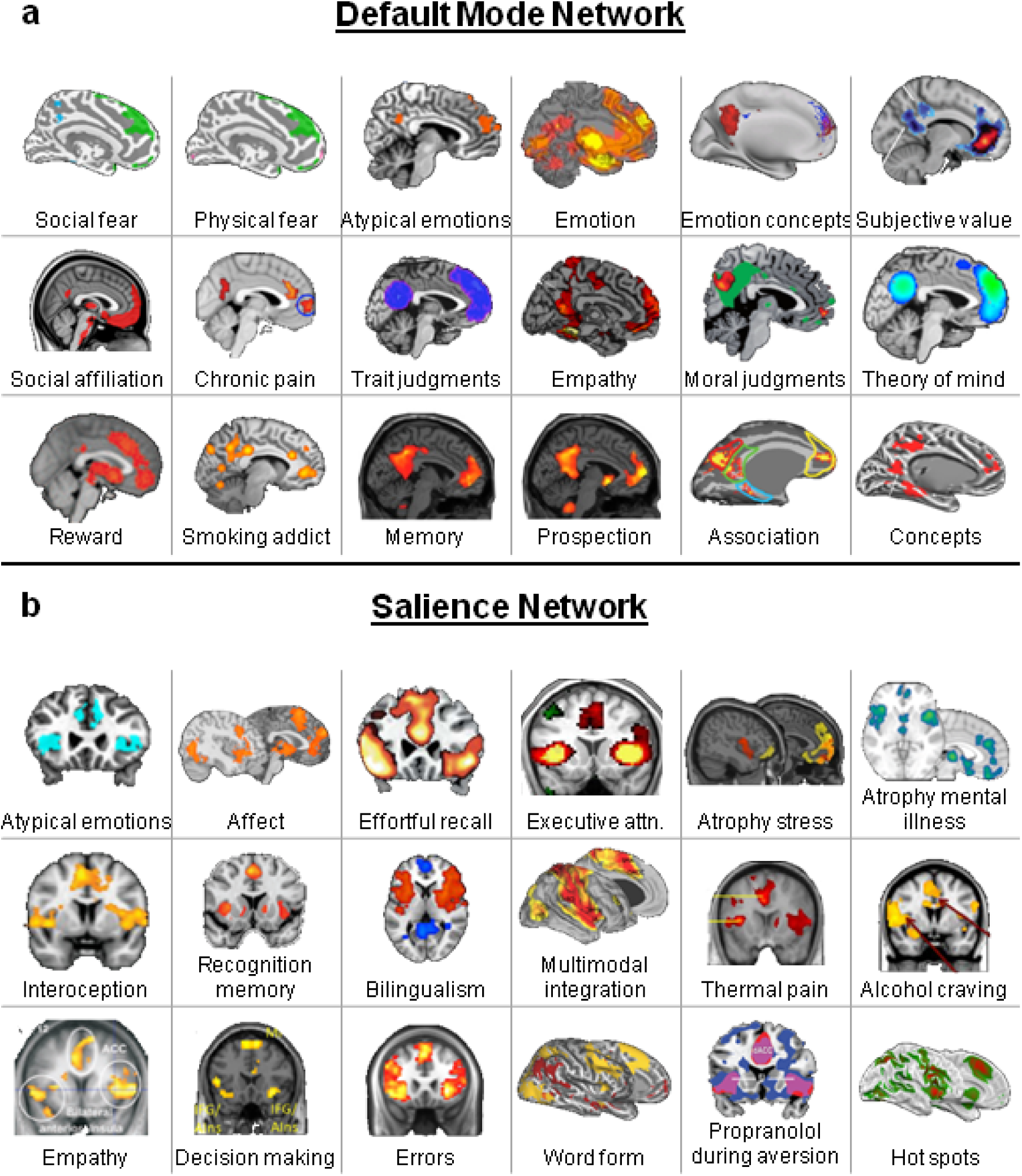
The default mode and salience networks each support a wide array of psychological functions, as evidenced by a literature review of psychological or other states that are sensitive to functional or structural features of these networks. These results are consistent with the idea that the default mode and salience networks are domain-general networks that support interoception and allostasis, which we propose are key processes that contribute to all psychological functions. Each sub-figure shows a set of results from an independent study, with citations as follows. Default mode network: Social fear^184^, Physical fear^184^, Atypical emotions^185^, Emotion^186^, Emotion concepts^187^, Subjective value^188^, Social affiliation^189^, Chronic pain^190^, Trait judgments^191^, Empathy^192^, Moral judgments^193^, Theory of mind^191^, Reward^194^, Smoking addiction^195^, Memory^196^, Prospection^196^, Association^197^, and Concepts^198^. Salience network: Atypical emotion^185^, Affect^199^, Effortful recall^200^, Executive attention^201^, Atrophy and stress (chronic yellow, current red)^202^, Atrophy and mental illness^121^, Interoception ^203^, Recognition memory, Bilingualism^205^, Multimodal integration^1^, Thermal pain, Alcohol craving, Empathy^208^, Decision making^209^, Errors^210^, Word form (yellow)^211^, Propranolol during aversion^212^, and Hot spots^213^.

## Discussion

The integrated allostatic/interoceptive brain system is a complex cortical and subcortical system consisting of connected intrinsic networks. The novelty of our work is our demonstration of a single brain system that supports not just allostasis but also a wide range of psychological phenomena (emotions, memory, decision-making, pain) that can all be explained by their reliance on allostasis. Other studies have already shown that regions controlling physiology are also regions that control emotion. In fact, this was Papez’s original logic for assuming that the “limbic system” was functionally for emotion. This paper goes beyond this observation. Regions controlling and mapping of inner body physiology lie in networks that also social affiliation, pain, judgments, empathy, reward, addiction, memory, stress, craving, decision making, etc. (Fig. 5). More and more, functional imaging studies are finding that the salience and default mode networks are domain-general (e.g., ^29^; for review, see ^27,28^). Our paper provides the groundwork for a theoretical and empirical framework for making sense of these findings in an anatomically principled way.

Our investigation was strengthened by our theoretical framework (the EPIC model^11^), the converging evidence from structural studies of the brain (i.e., tract-tracing studies in monkeys plus the well-validated structural model of information flow), our use of multiple methods (intrinsic connectivity in humans, as well as brain-behavior relationships), and our ability to replicate the system in three separate samples totaling over 600 human participants. Our results are consistent with prior anatomical and functional studies that have investigated portions of this system at cortical and subcortical levels (e.g.,^16,17,25,26 74,-77^), including evidence that limbic cortical regions control the brainstem circuitry involved with allostatic functions such as cardiovascular control, respiratory control, and thermoregulatory control^78^, as well as prior investigations that focused on the intrinsic connectivity of individual regions such as the insula (e.g., ^79^), the cingulate cortex (e.g., ^80^), the amygdala (e.g.,^81^), and the ventromedial prefrontal cortex (e.g., ^82^); importantly, our results go beyond these prior studies in several ways. First, we observed an often-overlooked finding when interpreting the functional significance of certain brain regions: the dorsomedial prefrontal cortex, the ventrolateral prefrontal cortex, the hippocampus, and several other regions have both a structural and functional pattern of connectivity that indicates their role in visceromotor control. A second often-overlooked finding is that relatively weaker connectivity patterns (e.g., between the visceromotor sgACC and primary interoceptive cortex) are reliable, and future studies may find that they are of functional significance. Third, we demonstrated behavioral relevance of connectivity within this network, something that prior studies of large-scale autonomic control networks have yet to test (e.g., ^74–76^). Taken together, our results strongly support the EPIC model’s hypothesis that visceromotor control and interoceptive inputs are integrated within one unified system^11^, as opposed to the traditional view that the cerebral cortical regions sending visceromotor signals and those that receive interoceptive signals are organized as two segregated systems, similar to the corticospinal skeletomotor efferent system and the primary somatosensory afferent system.

Perhaps most importantly, the allostatic/interoceptive system has been shown to play a role in a wide range of psychological phenomena, suggesting that allostasis and interoception are fundamental features of the nervous system. Anatomical, physiological, and signal processing evidence suggests that a brain did not evolve for rationality, happiness, or accurate perception; rather, all brains accomplish the same core task^19^: to efficiently ensure resources for physiological systems within an animal’s body (i.e., its internal milieu) so that an animal can grow, survive, thrive, and reproduce. That is, the brain evolved to regulate *allostasis*^20^. All psychological functions performed in the service of growing, surviving, thriving, and reproducing (such as remembering, emoting, paying attention, deciding, etc.) require the efficient regulation of metabolic and other biological resources.

Our findings add an important new dimension to the existing observations that the default mode and salience networks serve as a high-capacity backbone for integrating information across the entire brain^83^. Diffusion tensor imaging studies indicate, for example, that these two networks contain the highest proportion of hubs belonging to the brain’s “rich club,” defined as the most densely interconnected regions in the cortex^72,84^ (several of which are connector hubs within the allostatic/interoceptive system; see Fig. 3, Table S4). All other sensory and motor networks communicate with the default mode and salience networks, and potentially with one another, through these hub^1,84^. The agranular hubs within the two networks, which are also visceromotor control regions, are the most powerful predictors in the brai^11,31^. Indeed, hub regions in these networks display a pattern of connectivity that positions them to easily send prediction signals to every other sensory system in the brain^12,31^.

The fact that default mode and salience networks are concurrently regulating and representing the internal milieu, while they are routinely engaged during a wide range of tasks spanning cognitive, perceptual, and emotion domains, all of which involve value-based decision-making and action^85^ (e.g., ^86–88^; ^29^; for a review, see ^87^), suggest a provocative hypothesis for future research: whatever other psychological functions the default mode and salience networks are performing during any given brain state, they are simultaneously maintaining or attempting to restore allostasis and are integrating sensory representations of the internal milieu with the rest of the brain. Therefore, our results, when situated in the published literature, suggest that the default mode and salience networks create a highly connected functional ensemble for integrating information across the brain, with interoceptive and allostatic information at its core, even though it may not be apparent much of the time.

When understood in this framework, our current findings do more than just pile on more functions to the ever-growing list attributed to the default mode and salience networks (which currently spans cognition, attention, emotion, perception, stress, and action; see ^27,29^). Our results offer an anatomically plausible computational hypothesis for a set of brain networks that have long been observed but whose functions have not been fully understood. The observation that allostasis (regulating the internal milieu) and interoception (representing the internal milieu) are at the anatomical and functional core of the nervous system^17,19^ further offer a generative avenue for further behavioral hypotheses. For example, it has recently been observed that many of the visceromotor regions within the unified allostatic/interoceptive system contribute to the ability of *SuperAgers* to perform memory and executive function tasks like young^89^.

Furthermore, our findings also help to shed light on two psychological concepts that are constantly confused in the psychological and neuroscience literatures: affect and emotion. If, whatever else your brain is doing—thinking, feeling, perceiving, moving—it is also regulating your autonomic nervous system, your immune system, and your endocrine system, then it is also continually representing the interoceptive consequences of those physical changes. Interoceptive sensations are usually experienced as lower-dimensional feelings of affect^90,91^. As such, the properties of affect—valence and arousal^92,93^—can be thought of as basic features of consciousness^94–100^ that, importantly, are not unique to instances of emotion.

Perhaps the most valuable aspect of our findings is their value for moving beyond traditional domain-specific or “modular” views of brain structure/function relationships^101^, which assume a significant degree of specificity in the functions of various brain systems. A growing body of evidence requires that these traditional modular views be abandoned^27,102,103^in favor of models that acknowledge that neural populations are domain-general or multi-use. The idea of domain-generality even applies to primary sensory networks, as evidenced by the fact that multisensory processing occurs in brain regions that are traditionally considered unimodal (e.g., auditory cortex responding to visual stimulatio^104,105)^. The absence of specificity in brain structure/function relationships is not a measurement error or some biological dysfunction, but rather it is a useful feature that reflects core principles of biological degeneracy that are also evident in the genome, the immune system, and every other biological system shaped by natural selection^106^.

No study is without limitations. First, there are potential issues identifying homologous regions between monkey and human brains^46^; nonetheless, we still found evidence for the majority of the monosynaptic connections predicted by the EPIC model. Second, we used an indirect measure of brain connectivity in humans (functional connectivity analyses of low-frequency BOLD data acquired at rest) that reflects both direct and indirect connections and can, in principle, inflate the extent of an intrinsic network^46^. Moreover, low frequency BOLD correlations may reflect vascular rather than neural effects in brain^107^. Nonetheless, our results exhibit specificity: the integrated allostatic/interoceptive system conforms to well-established salience and default mode networks and is remarkably consistent with both cortical and subcortical connections repeatedly observed in tract-tracing studies of non-human animals. Third, although our fMRI procedures were not optimized to identify subcortical and brainstem structures and study their connectivity (e.g., ^56,74,75,108^), we nonetheless observed 92% of the predicted connectivity results. Finally, many studies find that activity in the default mode and salience networks have an inverse or negative relationship (sometimes referred to as “anticorrelated”), meaning that as one network increases its neural activity relative to baseline, the other decreases. Such findings and interpretations have recently been challenged on both statistical and theoretical grounds (e.g., ^109^; see Supplementary Results). In fact, when global signal is not removed in pre-processing, the two networks can show a pattern of positive connectivity (e.g., ^110^). Fourth, our demonstration of a brain/behavior relationship (using the evocative pictures) was merely a first look at how individual differences in the function of this system are related to individual differences in behavior. Additionally, our use of electrodermal activity as a measure of sympathetic nervous system activity is arguably too specific because different components of the sympathetic nervous system react differently^111^, and peripheral sensations associated with SCRs themselves might not be processed by the interoceptive brain circuitry that we are studying here, thus complicating the interpretation of our results. However, we did not intend to assess a particular path carrying information about SCRs specifically, and we believe that – despite their limitations – our results are still useful and hypothesis-generating. Future work will be needed to understand this and other brain/behavior relationships of this system more thoroughly.

This work is the first in a series of studies to precisely test the EPIC model, including its predictive coding features (not just the anatomical and functional correlates as shown here). Future research must focus on the ongoing dynamics by which the default mode and salience networks support allostasis and interoception, including the predictions they issue to other sensory and motor systems. It is possible, for example, that both networks use past experience in a generative way to issue prediction signals, but that the default mode network generates an internal model of the world via multisensory predictions (consistent with ^112–114^), whereas the salience network issues predictions, as precision signals, to tune this model with prediction error (consistent with the salience network’s role in attention regulation and executive control; e.g., ^50,115,116^). Unexpected sensory inputs that are anticipated to have allostatic implications (i.e., likely to impact survival, offering reward or threat) will be encoded as “signal” and learned to better support allostasis in the future, with all other prediction error is treated as “noise” and safely ignored (^117^; for discussion, see ^118^). These and other hypotheses regarding the flow of predictions and prediction errors in the brain (e.g., incorporating the cerebellum, ventral striatum, and thalamus^23^ can be tested using new methods such laminar MRI scanning at high (7 T) magnetic field strengths (e.g., ^119^).

Future research that provides a more mechanistic understanding of how the default mode and salience networks support interoception and allostasis will also reveal new insights into the mind-body connections at the root of mental and physical illness and their comorbidities. For example, in illness, the neural representations of the world that underlie action and experience may be directed more by predicted allostatic relevance of information than by the need for accuracy and completeness in representing the environment. Indeed, atrophy or dysfunction within parts of the interoceptive system are considered common neurobiological substrates for mental and physical illness^120–112^, including depression^123^, anxiety^124^, addiction^125^, chronic Pain1^26^, obesity ^127^, and chronic stress^128,129^. By contrast, increased cortical thickness in MCC is linked to the preserved memory of *SuperAgers* relative to their more typically performing elderly peers^130,131^, suggesting a potential mechanism for how exercise (via the sustained visceromotor regulation it requires) benefits cognitive function in aging^132^ and why certain activities, such as mindfulness or contemplative practice, can be beneficial (e.g., ^133,134^). Ultimately, a better understanding of how the mind is linked to the physical state of the body through allostasis and interoception may help to resolve some of the most critical health problems of our time, such as the comorbidities among mental and physical disorders related to metabolic syndrome (e.g., depression and heart disease^135^), or how chronic stress speeds cancer progression^136^, as well as offer key insights into how an opioid crisis^137^ and recorded numbers of suicides^138^ emerge.

## Author contributions

The study was designed by all the authors, analyzed by all the authors, and the manuscript was written by I.R.K. and L.F.B with comments and edits from other authors. The authors acknowledge Miguel Angel Garcia-Cabezas for comments and advice on neuroanatomy and Henry Evrard for helpful discussions on anatomical connectivity. This research was supported by the National Institutes on Aging (R01 AG030311) to L.F.B. and B.C.D., the US Army Research Institute for the Behavioral and Social Sciences Contracts (W5J9CQ-11-C-0046 and W5J9CQ-12-C-0049) to L.F.B., the National Institute of Mental Health Ruth L. Kirschstein National Research Service Award (F32MH096533) to I.R.K., as well as the National Institutes of Mental Health (K01MH096175-01) and Oklahoma Tobacco Research Center grants to W.K.S, and the Fonds de recherche sante Quebec fellowship award to C.X. The views, opinions, and/or findings contained in this paper are those of the authors and shall not be construed as an official Department of the Army position, policy, or decision, unless so designated by other documents.

## Online Methods

### Materials and Methods

#### Participants

##### Discovery and replication samples

We randomly selected 660 participants (365 female, 55%) with age range 18-30 years from a core dataset of 1,000 healthy young adults originally presented by Buckner and colleagues^54,214^. The 1,000 participants were clinically healthy without a history of neurological or psychiatric conditions, native English-speaking adults (ages 18-35) with normal or corrected-to-normal vision. Additional details about the sample are discussed in Yeo, et al.^54^ We removed 79 participants (11%) due to head motion and outlying voxel intensities using established criteria (Analysis of Functional Neuroimages, AFNI; http://afni.nimh.nih.gov); we removed an additional 31 participants (4.7%) due to lack of signal in the most superior and lateral parts of the brain (see Analysis section). Our final dataset consisted of 550 participants with high quality fMRI BOLD data ready for further analysis. These 550 participants were randomly divided into a discovery sample of *N* = 280 (174 female, 62%, mean age = 19.3 years, SD = 1.4 years) and a replication sample of *N* = 270 (142 female, 53%, mean age = 22.3 years, SD = 2.1 years).

In addition, from the entire participant pool (*N* = 1,000), we randomly selected 150 participants (75 female, 50%, mean age = 22.5, SD = 2.0 years) to generate maps of the established default mode and salience networks.

##### Validity sample

We selected all 66 young and middle aged participants (33 female) with age range 18-60 years (mean age = 34.8 years, SD = 13.8 years) from an existing dataset of 111 participants (56 female) with age range 18-81 years, mean age 46.6 years, SD 18.9 years) recruited from the Boston area during 2012-2014 for a study examining age-related changes in how affect supports memory^215^. Only 41 participants (14 female, 47%) with age range 20-60 years (mean age = 33.8 years, SD = 14.1 years) had both high quality fMRI BOLD data and sufficient electrodermal activity, according to previously established quality assessment procedures described in the Analysis section (12 participants were disqualified for excessive head motion and outlying voxel intensities; 16 participants were disqualified for lack of electrodermal responses). Participants were right-handed, native English speakers and had normal or corrected-to-normal vision. None reported any history of neurologic or psychiatric condition, learning disability or serious head trauma. Participants did not smoke and did not ingest substances that interfere with autonomic responsiveness (e.g., beta-blockers or anti-cholinergic medications).

**Sample size. No pre-specified effect size was known, so we used a large portion of a third-party dataset (N=660) and the maximum size of a second dataset collected in our lab with young and middle-aged adults (N=66).Procedure**

##### Discovery and replication samples

Participants were greeted and provided written informed consent in accordance with the guidelines set by the institutional review boards of Harvard University or Partners Healthcare. Participants completed MRI structural and resting state scans, and other tasks unrelated to the current analysis. MRI data were acquired at Harvard and the Massachusetts General Hospital across a series of matched 3T Tim Trio scanners (Siemens, Erlangen, Germany) using a 12-channel phased-array head coil. Structural data included a high-resolution multiecho T1-weighted magnetization-prepared gradient-echo image (multiecho MP-RAGE). Parameters for the structural scan were as follows: repetition time (TR) = 2,200 ms, inversion time (TI) = 1,100 ms, echo time (TE) = 1.54 ms for image 1 to 7.01 ms for image 4, flip angle (FA) = 7°, 1.2 × 1.2 × 1.2-mm voxels, and field of view (FOV) = 230 mm. The functional resting state scan lasted 6.2 min (124 time points). The echo planar imaging (EPI) parameters for functional connectivity analyses were as follows: TR = 3,000 ms, TE = 30 ms, FA = 85°, 3 × 3 × 3-mm voxels, FOV = 216 mm, and 47 axial slices collected with interleaved acquisition and no gap between slices.

##### Validity sample

Participants were greeted and consented in accordance with the institutional review board. Data were acquired on separate sessions across several few days. The first laboratory testing session consisted of a 6-min seated baseline assessment of peripheral physiology, the EXAMINER cognitive battery^216^, a second 6-min seated baseline, the evocative images task, and other tasks unrelated to the current study. Only the evocative images task is relevant for this study. Electrodes were placed on the chest, hands, and face to record electrocardiogram, skin conductance, and facial electromyography, respectively. A belt with a piezoelectric sensor was secured on the chest to record respiration. Only the skin conductance data are reported here. Skin conductance responses were recorded using disposable electrodermal electrodes (containing isotonic paste) affixed to the thenar and hypothenar eminences of the left hand. Data were collected using Mindware’s data acquisition software (BioLab Acquisition Software version 3.0.13; Mindware Technologies, Gahanna, OH, USA). During the task, participants sat upright in a comfortable chair in a dimly lit room. After a 6-min baseline of seated rest, ninety full-color photos were selected from the International Affective Picture System (IAPS) and used to induce affective experiences^60^. The pictures were selected based on normative ratings of pleasure/displeasure (valence) and arousal experienced when viewing them (i.e. unpleasant-high arousal, pleasant-high arousal, unpleasant-low arousal, pleasant-low arousal, neutral valence-low arousal). The 90 IAPS numbers are shown in Table S5. Participants viewed each of the ninety IAPS photos sequentially on a 120 × 75 cm high definition (Sharp, Aquos) screen two meters away. Photos were grouped into three blocks of thirty each, with the order of the photos within each block fully randomized. For each trial, participants viewed an IAPS photo for six seconds, and then rated their experience for valence and arousal using the Self Assessment Manikin (SAM^217^). Only the arousal ratings are relevant to this report and they ranged from 1 (“Very calm”) to 5 (“Very activated”). A variable inter-trial interval of 10-15 seconds followed the rating prior to presentation of the next picture. Before beginning the task, participants were familiarized with the SAM rating procedure and practiced by rating five pictures. The photos and rating scales were administered via E-Prime software (Psychology Software Tools, Pittsburgh, PA).

The second laboratory testing session involved MRI scanning, consisting of a structural scan, resting state scan, and other tasks unrelated to the present report (but presented here^215^) including an affect induction and memory task. MRI data were acquired using a 3T Tim Trio scanner (Siemens, Erlangen, Germany) using a 12-channel phased-array head coil. Structural data included a high-resolution T1-weighted MP-RAGE with TR = 2,530 ms, TE = 3.48 ms, FA = 7°, and 1 mm isotropic voxels. The functional resting state scan lasted 6.40 min (76 time points). The EPI parameters were as follows: TR = 5,000 ms, TE = 30 ms, FA = 90°, 2 mm isotropic voxels, and 55 axial slices collected with interleaved acquisition and no gap between slices. Participants were instructed to keep their eyes open without fixating and remain as still as possible.

**Analysis of task-independent (“resting state”) functional magnetic resonance imaging (fMRI) data**

##### Quality assessment

We applied established censoring protocols for head motion and outlying signal intensities using AFNI following Jo, et al. ^218^ and described in the following three steps: First, we disqualified an fMRI volume if AFNI’s *enorm motion* derivative parameter (derived from *afni_proc.py*) was greater than 0.3 mm. Second, we also disqualified an fMRI volume if the fraction of voxels with outlying signal intensity (AFNI’s *3dToutcount* command) was greater than 0.05. Third, if a volume surpassed either criterion, then we removed that volume, the prior volume, and the next two volumes. In a separate procedure, we disqualified discovery and replication participants who lost more than 10% of their 124 volumes due to either criterion (79 participants, 11%). Quality assessment for surface-based processing required removing 31 additional participants (4.7%) due to a lack of signal in the most superior and lateral parts of the brain, which would result in incomplete group connectivity maps; no participants were removed for this reason in the validity sample. In the validity sample, we removed participants who lost more than 40% of their 76 volumes, removing 12 participants (18%); we used a more lenient threshold due to the small sample size (*N* = 66). The fraction of volumes censored per participant using the aforementioned approach by Jo, et al.^141^ yielded nearly identical results to another established censoring approach described in Power, et al. ^219^ as implemented in AFNI’s *afni_restproc.py* script.

##### Preprocessing

We applied standard Freesurfer preprocessing steps to both samples of resting state data (http://surfer.nmr.mgh.harvard.edu). These included removal of the first four volumes, motion correction, slice timing correction, resampling to the MNI152 cortical surface (left and right hemispheres) and MNI305 subcortical volume (2 mm isotropic voxels), spatial smoothing (6 mm FWHM, surface and volume separately) and temporal filtering (0.01 Hz high-pass filter and 0.08 Hz low-pass filter). We did not use global signal regression to prevent spurious negative correlations (“anti-correlated networks”), which can interfere with interpreting the connectivity results^109^.

##### Functional connectivity analysis

We estimated cortical connectivity using surface-based analyses, affording more sensitive and reliable discovery maps and reducing artifacts around sulcal and opercular borders (which are more apparent in traditional volume-based connectivity methods)^220^. The surface-based intrinsic analyses also allowed us to incorporate the selected subcortical seed (dAmy), but did not allow us to analyze connectivity to subcortical structures more broadly (see next paragraph for description of more comprehensive cortical-subcortical connectivity analyses). The surface-based method registers a participant’s native space to MNI152 space via Freesurfer’s reconstruction of each participant’s cortical surfaces. Therefore, compared to volumetric processing in a standard space, this method facilitates more accurate signal extraction on an individual level. We first created a 4-mm sphere centered on the MNI coordinates identified in Table 3 and found the vertex on the MNI152 pial surface that is closest to the spherical seed. We then smoothed this single vertex by 4 mm on the surface and mapped the resulting cortical label to each individual subject’s cortex. The individual cortical label was projected back into the subject’s native volumetric space to calculate the averaged time series within the seed. For the subcortical seed (dAmy), we directly projected the spherical seed into each subject’s native volumetric space and extracted its time course. On the subject level, we ran a voxel-wise regression on left and right hemispheres of MNI152 and subcortical volume of MNI305 to compute the partial correlation coefficient and correlation effect size of the seed time series, taking into account several nuisance variables: cerebrospinal fluid signal, white matter signal, motion correction parameters, and a 5^th^ order polynomial. On the group level, we concatenated the contrast effect size maps from all subjects and ran a general linear model analysis to test if the group mean differed from zero. This yielded final group maps that showed regions whose fluctuations significantly correlated with the seed’s BOLD time series.

To estimate cortical-subcortical connectivity, we used a more liberal statistical threshold (as compared to the analyses of corticocortical connectivity). The smaller size of subcortical regions, as well as their anatomical placement, renders their signal noisier and less reliable^56^, yielding relatively smaller estimates of intrinsic connectivity (i.e., the strength of connection between two groups of voxels is estimated as the magnitude of correlation between their BOLD time courses, which is ultimately limited by the reliability of the two signals). Thus, guided by classical measurement theory^183^, we relied on replication to determine which connectivity values were meaningful (because only reliable signal replicates across samples; random noise does not).

##### K-means cluster analysis of discovery maps

First, we computed the 8×8 *η*^2^ similarity matrix for each pair of maps^48^. Based on visual inspection of the eight maps, we used K-means clustering with *k* = 2 and *k* = 3 using the *kmeans* function in MATLAB (Mathworks, Natick, MA). Our results confirmed that *k* = 2 captured the default mode versus salience distinction across these maps, whereas *k* = 3 further divided the ‘salience cluster’ into two sub-categories depending on whether or not somatosensory cortices are included. Because sub-categories within the salience network were not important to our study goals, we used the *k* = 2 cluster solution.

##### Identification of the interoceptive system networks

We confirmed that Network 1 is the established default mode network (for a review, see ^49^) and Network 2 is the established salience network^51,52^. The reference maps were constructed using coordinates obtained from Yeo, et al.^54^ as follows. Using a random sample of *N* = 150, we created a mask of the default mode network by conjoining functional connectivity maps from two hubs in the default mode network^54^: a 4-mm seed at the dorsomedial prefrontal cortex (MNI 0, 50, 24) and a 4-mm seed at the posterior cingulate cortex (MNI 0, -64, 40). We likewise created a mask of the salience network by conjoining functional connectivity maps from two bilateral hubs in the salience network (labeled as the ventral attention network in Yeo, et al.^54^): 4-mm seeds at the left and right supramarginal gyrus (MNI ±60, -30, 28) and 4-mm seeds at the left and right anterior insula (MNI ±40, 12, -4). We thresholded our maps to *p* < 10^-5^ uncorrected (as in all our analyses) and we thresholded the default mode and salience networks to *z*(*r*) > 0.05 where z is the Fisher’s r-to-z transformation. We then calculated the percent of each established network (default mode or salience) that covered each of our networks (Network 1 or 2), and the complementary measure: the percentage of each of our networks (Network 1 or 2) that covered each established network (default mode or salience). These calculations used only the right hemisphere.

##### Reliability analyses

We used *η*^2^ as an index of reliability because it shows similarity between maps while discounting scaling and offset effects^48^. An *η*^2^ value of 1 indicates spatially identical maps, while an *η*^2^ value of 0.5 indicates statistically independent maps. For each of our eight cortical and amygdalar seeds, we calculated *η*^2^ between the discovery and replication samples using the effect size (gamma) maps generated by the group-level general linear model analysis. Then we calculated the mean and SD of the eight *η*^2^ values across all seeds to index overall similarity between samples. This was done separately for the cortical and subcortical maps. We repeated the same procedure to compare the reliability between the discovery and validation samples.

### Analysis of the evocative images task

We analyzed skin conductance data using Mindware software for scoring electrodermal activity (Electrodermal Activity/Skin Conductance Analysis version 3.0.21). For each 6-second trial when the test photo was visible, we measured the number of event-related phasic skin conductance responses (SCRs) according to best practices^221^. We considered an SCR to be event-related if both the response onset and peak occurred between 1 and 6 seconds after stimulus onset, with an onset to peak amplitude of at least 0.01 μS. It is commonly observed that a substantial proportion of healthy adults produce relatively few if any SCRs (i.e., they are considered “stabile” electrodermal responders)^222^. We disqualified 16 of our 66 participants (24%) because they were stabile (they generated event-related SCRs during fewer than 5% of the evocative photo trials). We analyzed our data using the number of SCRs (as opposed to the amplitude of the SCRs) model because it was consistent with prior work from our lab (e.g., ^223^) and prior work from other labs (e.g., ^224^).

#### Multilevel linear modeling to assess correspondence between objective physiological and subjective arousal during an allostatically relevant task

Analyses were conducted using HLM version 7.01 (Student Version) and the reported parameters and statistics reflect HLM’s robust estimates. Level-1 of the multi-level regression model estimated the linear relationship (slope and intercept) between physiological arousal (number of event-related skin conductance responses (NSCRs) and subjective arousal (rated from 1 = “Very calm” to 5 = “Very activated”) in response to each of ninety photos. Thus, the model was adjusted for mean individual reactivity. Level-2 of the regression model estimated the extent to which intrinsic connectivity between viscerosensory and visceromotor regions (e.g., dpIns-aMCC connectivity) moderated the relationship between objective and subjective arousal (i.e., moderated the slope of the Level 1 model). All variables were unstandardized. Level-1 variables were group-mean centered (for each participant) and Level-2 variables were grand-mean centered (across participants).

## Supplementary Results

### Detailed description of subcortical, hippocampal, brainstem, and cerebellar connectivity within the interoceptive system

Here, we briefly justify the inclusion of each non-cortical region in our connectivity analyses, report its observed intrinsic connectivity patterns with the allostatic/interoceptive system seeds (Fig. 4 and Fig. S6), and compare our results with published tract-tracing studies showing monosynaptic anatomical connectivity among these regions (Table 2). Discrepancies between our results and tract-tracing are only indicated if our fMRI results failed to show connectivity between regions that are monosynaptically connected. fMRI intrinsic connectivity reflects both direct and indirect (multisynaptic) connections^46,47^ and thus our results sometimes show connectivity for regions that are disynaptically connected.

### Thalamus

We examined connectivity to two thalamic nuclei: the posterior part of the ventromedial nucleus (VMpo; for a review, see ^16^) for interoceptive input specifically and the larger ventral posterior (VP) complex for somatic input more broadly^55^. Our fMRI results revealed that all cortical and amygdalar seeds exhibited connectivity with the VMpo and VP. This is entirely consistent with tract-tracing studies showing direct projections of VMpo or VP to dpIns, dmIns, and dorsal ACC/aMCC (for a review, see ^16^). Our other cortical and amygdalar seeds have multisynaptic connectivity to VMpo and VP by way of aMCC.

### Hypothalamus

The hypothalamus is a critical region for allostatic regulation of the body; the paraventricular nucleus in the medial zone is particularly responsible for visceromotor control of autonomic, endocrine, and immune function^225^. Our review of tract tracing studies clearly indicated connectivity from each cortical and amygdalar seed except with dpIns, despite evidence from tract-tracing, and with lvaIns to the hypothalamus (which was not predicted) ^42,58,140,145,156,157,169^. The lack of functional connectivity between lvaIns and the hypothalamus is not surprising because the lateral portion of the ventral anterior insula is part of a sensory integration network in orbitofrontal cortex in monkeys and humans^42,58^ that has little connection to the hypothalamus, except for light connections to the posterior lateral hypothalamus at the level of the mammillary bodies^42^. We were unable to identify the expected connectivity between the dAmy and any part of the hypothalamus.

### Cerebellum

The cerebellum is a key structure in sensorimotor regulation because it sends efferent copies (i.e., predictions) to the cortex to help the brain distinguish between the anticipated sensory consequences of self-initiated by the body vs. those that are unexpected^226,227^. All cortical seeds and the dAmy seed exhibited connectivity with the cerebellum. More specifically, all seeds exhibited connectivity to lobules IV, V, VI and VIIIB, consistent with the cerebellar “somatosensory” network. The default mode portion of the interoceptive system is additionally connected to lobule IX and Crus I, whereas the salience portion of system is additionally connected to lobule VIIIA (for a specific parcellation of cerebellar intrinsic connectivity, see ^214^).

### Amygdala

The amygdala is a key subcortical region for both interoceptive input (via its lateral nucleus) and visceromotor control (via its central nucleus)^228^, and is part of both the salience and the default mode networks^49,51^. All of our cortical seeds exhibited connectivity to the amygdala seed except for the pACC seed, which had limited connectivity to the left amygdala (only the dorsal section, which contains the central nucleus). This is consistent with results from tract-tracing studies, which show that each of our cortical seeds projects monosynaptically to the dorsal amygdala^59,140,142,144,151,152^.

### Hippocampus

The hippocampus is a key subcortical hub in the default mode network^49^ that is strongly connected to the amygdala (for a review, see ^229^). Our fMRI results showed that all cortical seeds and the dAmy seed exhibited connectivity to the entire hippocampus except for the aMCC seed, which exhibited connectivity to only the posterior hippocampus, and the dmIns seed, which exhibited connectivity only to the anterior and posterior hippocampus. Our results are consistent with tract-tracing studies indicating direct projections from the amygdala to the hippocampus and indirect projections from many regions of the cortex to the hippocampus; specifically, vaIns, sgACC, pACC, and aMCC all project to the entorhinal cortex, which projects to the hippocampus (for a review, see ^229^). Other cortical regions such as the aMCC and dmIns can connect to the hippocampus in three steps: first to a cortical hub (e.g., vaIns), then to and the entorhinal cortex, then to the hippocampus.

### Ventral striatum

The striatum is a subcortical region in the basal ganglia comprising the caudate, putamen, and nucleus accumbens, and its ventral portion is important for controlling inhibitory signals to brainstem visceromotor targets^230,231^Specifically, cortical regions send excitatory (glutamatergic) signals to the striatum, which enhance inhibition of brainstem visceromotor targets; these connections also have the capacity to release visceromotor targets from tonic inhibition via striatal connections to the pallidum^231^. All of our cortical and amygdalar seeds exhibited connectivity to the ventral striatum, except for dmIns, which exhibited connectivity to a portion of the ventral striatum. This is consistent with tract-tracing studies showing monosynaptic connections from each of our seeds to the ventral striatum^149,153,157,160,163,164^.

### Periaqueductal gray (PAG)

The PAG is a midbrain nucleus important visceromotor control^232^. It is difficult to precisely localize using 3-T scanning procedures because it encircles the cerebral aqueduct (e.g., ^108^). Nonetheless, the results of our intrinsic connectivity analysis largely mirror those for the hypothalamus. The tract-tracing literature has identified monosynaptic connectivity to the PAG from all seeds^60,121^ except the dmIns and dpIns^146^, and the lvaIns^146^. All cortical visceromotor seeds demonstrated the expected connections with the PAG: aMCC, pACC, and sgACC. We did not observe the expected connectivity with the mvaIns nor with the dAmy. As expected, connectivity with the lvaIns and dpIns was not observed.

### Parabrachial nucleus (PBN)

The PBN is a nucleus in the pons that relays interoceptive input from the body to the brain^16^ and also serves visceromotor functions^233^. All cortical seeds and the dAmy seed exhibited connectivity with the PBN. This is in agreement with the tract-tracing literature showing monosynaptic connectivity to the PBN from each of our seeds^147,148,157,160,164^ except for the aMCC^162^, which must first project to another cortical hub (e.g., vaIns) before projecting to the PBN.

### Nucleus of the solitary tract (NTS)

The NTS is a key relay nucleus in the medulla that is on a cranial interoceptive pathway from the viscera to the brain^12,13,22,23^ and also contributes to visceromotor control^16,17,25,26,233^ We observed NTS connectivity with the dAmy seed and with all cortical seeds except for the sgACC. This is largely consistent with the tract-tracing literature showing monosynaptic projections to the NTS from each of our seeds^148,158,159,164^. Failure to observe the sgACC connection is perhaps due to the small size of the sgACC and increased noise due to partial-volume effects of the nearby white matter in the corpus callosum.

### Laboratory validation of the allostatic/interoceptive system in humans

The following text details our findings of the association between connectivity in the allostatic/interoceptive system in humans and an index of interoception: the concordance between objective and subjective measures of bodily arousal. We measured sympathetic nervous system arousal using SCRs^234^ while participants viewed each photo for six seconds. We selected SCRs as an index of sympathetic nervous system activity, and that although effects from SCRs specifically might not ascend to interoceptive brain systems, the simultaneous non-SCR effects of sympathetic nervous system activity likely are processed through interoceptive pathways. After each image, participants reported their subjective experience of arousal using a validated self-report scale^217^. Using multi-level regression procedure to account for the nested and repeated-measures design of this experiment (for a review, see ^235^), we found that the number of SCRs in response a picture (either 0, 1, 2, 3, or 4 SCRs to a given picture) predicted the intensity of arousal experiences in response to the same picture across all participants and pictures (*B* = 0.21, *p* < 0.001), consistent with prior research (e.g., ^236^). Furthermore, as predicted, individuals with stronger intrinsic connectivity between primary interoceptive cortex (dpIns) and the aMCC had a stronger correspondence between sympathetic arousal and subjective experience of arousal than did those with weaker connectivity (regression *B* = 0.56; *p* < 0.003; Fig. S8). We focused on the aMCC because it was an a priori visceromotor seed region (Table 1), a connector hub^72^, and consistently replicated tract-tracing connectivity to non-cortical allostatic nuclei. For completeness, our results focused on the number of SCRs in response to each image and we did not find analogous results using the amplitude of SCRs in response to each image.

### Reconciling prior studies that reported a negative correlation between default mode and salience network activity

Many studies have found that the default mode and salience networks have task-related activity that is negatively correlated (i.e., when the BOLD signal in one network goes up, the BOLD signal in the other network goes down). Such findings are often interpreted as evidence that the brain has either an internal focus on an external focus. This inverse relationship is often the consequence of removing the mean signal change from all voxels in each volume before proceeding with data analysis (called “global-signal regression”; e.g., ^109^). A more reasonable interpretation, however, may be that when one network increases its neural activity compared to some baseline, the other might show a relative decrease in activity from that baseline (which does not mean that network is irrelevant to the task at hand). Alternatively, one network might show a smaller increase than the other (which, when mean signal change is removed, would appear as a negative relationship between the two networks).

**Table S1.**
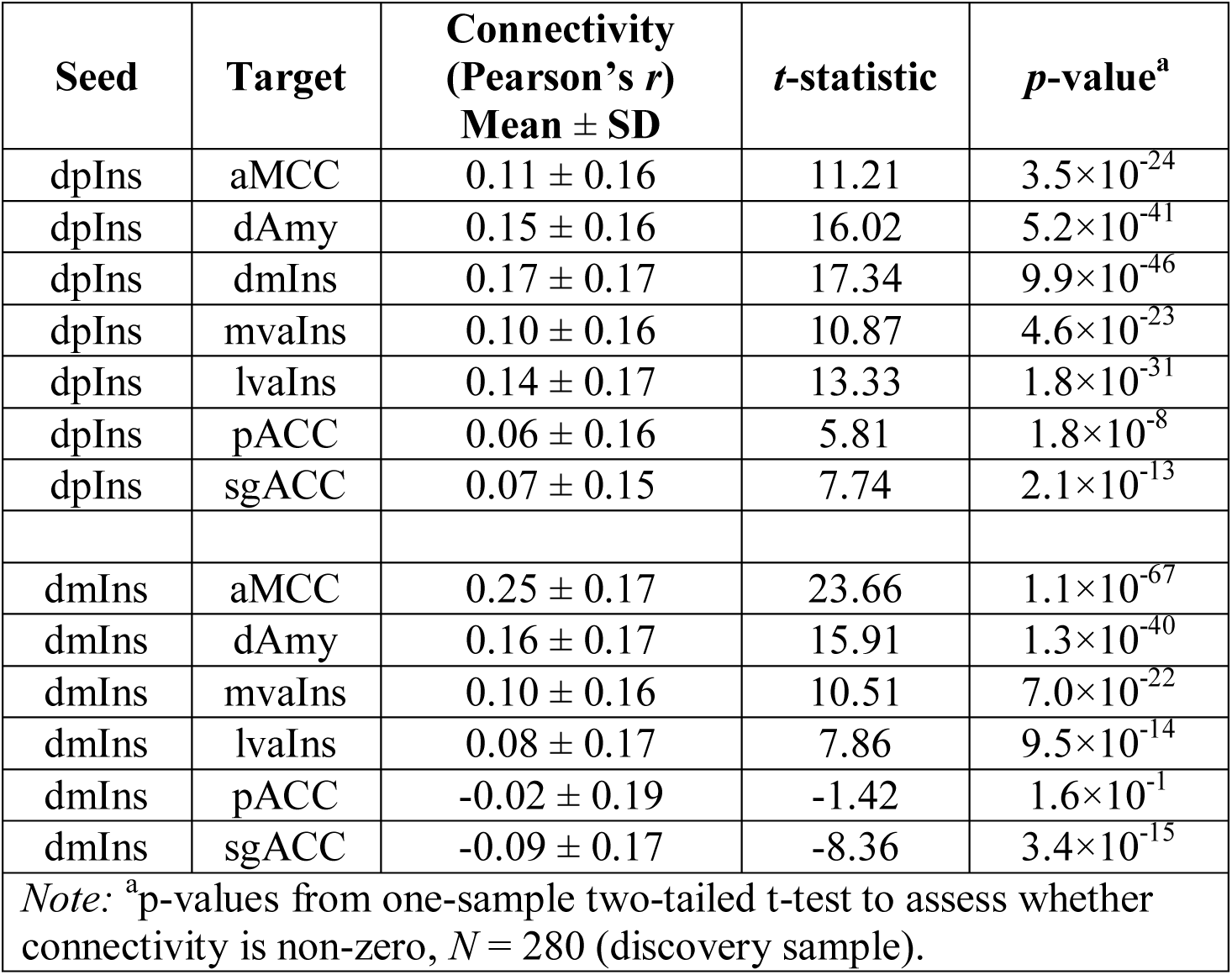
Functional connectivity between interoceptive and visceromotor regions in humans.

| Seed | Target | Connectivity<br>(Pearson's $r$)<br>Mean ± SD | $t$-statistic | $p$-value<sup>a</sup> |
| --- | --- | --- | --- | --- |
| dpIns | aMCC | 0.11 ± 0.16 | 11.21 | 3.5×10<sup>-24</sup> |
| dpIns | dAmy | 0.15 ± 0.16 | 16.02 | 5.2×10<sup>-41</sup> |
| dpIns | dmIns | 0.17 ± 0.17 | 17.34 | 9.9×10<sup>-46</sup> |
| dpIns | mvaIns | 0.10 ± 0.16 | 10.87 | 4.6×10<sup>-23</sup> |
| dpIns | lvaIns | 0.14 ± 0.17 | 13.33 | 1.8×10<sup>-31</sup> |
| dpIns | pACC | 0.06 ± 0.16 | 5.81 | 1.8×10<sup>-8</sup> |
| dpIns | sgACC | 0.07 ± 0.15 | 7.74 | 2.1×10<sup>-13</sup> |
| dmIns | aMCC | 0.25 ± 0.17 | 23.66 | 1.1×10<sup>-67</sup> |
| dmIns | dAmy | 0.16 ± 0.17 | 15.91 | 1.3×10<sup>-40</sup> |
| dmIns | mvaIns | 0.10 ± 0.16 | 10.51 | 7.0×10<sup>-22</sup> |
| dmIns | lvaIns | 0.08 ± 0.17 | 7.86 | 9.5×10<sup>-14</sup> |
| dmIns | pACC | -0.02 ± 0.19 | -1.42 | 1.6×10<sup>-1</sup> |
| dmIns | sgACC | -0.09 ± 0.17 | -8.36 | 3.4×10<sup>-15</sup> |
| *Note:* <sup>a</sup>$p$-values from one-sample two-tailed $t$-test to assess whether connectivity is non-zero, $N = 280$ (discovery sample). | | | | |

**Table S2.** Correspondence between networks in the interoceptive system and established resting state networks in humans.

| **Percent of networks 1 and 2 in each established resting state network** |  |  |  |
| --- | --- | --- | --- |
|  |  | Default mode network | Salience network |
| Network 1 | Discovery | **63%** | 15% |
|  | Replication | **75%** | 15% |
| Network 2 | Discovery | 7% | **60%** |
|  | Replication | 6% | **57%** |
| **Percent of each established resting state network in networks 1 and 2** |  |  |  |
|  |  | Default mode network | Salience network |
| Network 1 | Discovery | **77%** | 14% |
|  | Replication | **86%** | 14% |
| Network 2 | Discovery | 9% | **65%** |
|  | Replication | 9% | **64%** |
| *Note:* The percent in each cell indicates the fraction of the brain map in the row that overlaps with the brain map in the column. The top part of the table shows that the majority of Network 1 overlaps with the default mode network and the majority of Network 2 overlaps with the salience network. The bottom part of the table shows that the majority of the default mode network overlaps with Network 1 and the majority of the salience network overlaps with Network 2.<br>Calculations of networks 1 and 2 were performed at the threshold shown in Fig. 3 ($p < 10^{-5}$ uncorrected using $N = 280$). The default mode and the salience networks were defined based on default mode and ventral attention networks from Yeo, et al.<sup>54</sup> at a threshold of $z(r) > 0.05$ where $z$ is the Fisher's $z$ transformation. |  |  |  |

**Table S3.** Anatomical connections from the interoceptive system to visceromotor control regions (hypothalamus or PAG).

|  | **Cortical region** | **To hypothalamus** | **To PAG** |
| --- | --- | --- | --- |
| **Default Mode Network Only** | From ventromedial prefrontal cortex (BA 12) | Yes | Yes |
|  | From medial orbitofrontal cortex (BA 11, 13) | Yes | No |
|  | From dorsomedial prefrontal cortex (BA 9, 10) | Yes | Yes |
|  | From posterior cingulate cortex (BA 23, 31) | No | BA 23ab but not BA 31 |
|  | From ventral precuneus (BA 7) | No | No |
|  | From angular gyrus (BA 39) | No | No |
|  | From middle frontal gyrus (BA 46) | Yes | Yes |
| **Salience Network Only** | From dorsal precuneus (BA 5) | No | No |
|  | From medial frontal gyrus (BA 6) | Yes | No |
|  | From fusiform gyrus (BA 37) | No | No |
|  | From middle temporal gyrus (BA 21) | No | Yes |
|  | From precentral gyrus (BA 6) | Yes | Yes |
| **Hubs Connecting Networks Default Mode and Salience Networks** | From midcingulate cortex (BA 31) | Yes | Yes |
|  | From medial postcentral gyrus (BA 6) | No | No |
|  | From parahippocampal gyrus (BA 30) | No | No |
|  | From inferior temporal gyrus (BA 36) | No | No |
|  | From cuneus (BA 30) | No | No |
|  | From superior temporal sulcus (BA 13, 22) | No | Yes |
|  | From temporal pole (BA 38) | No | No |
|  | From inferior frontal gyrus (BA 47) | Yes | Yes |
| *Note:* Each cell shows the citation number for the paper indicating tract-tracing results in monkey (or sometimes in rat <sup>237</sup>). Hypothalamus projections obtained from references <sup>42,237</sup>; PAG projections obtained from references <sup>146,232,238,239</sup>. We only assessed projections from cortical regions to the hypothalamus or PAG (not from hypothalamus or PAG to cortical) because we wanted to assess if these cortical regions support visceromotor control.<br>PAG = periaqueductal gray. |  |  |  |

**Table S4.**
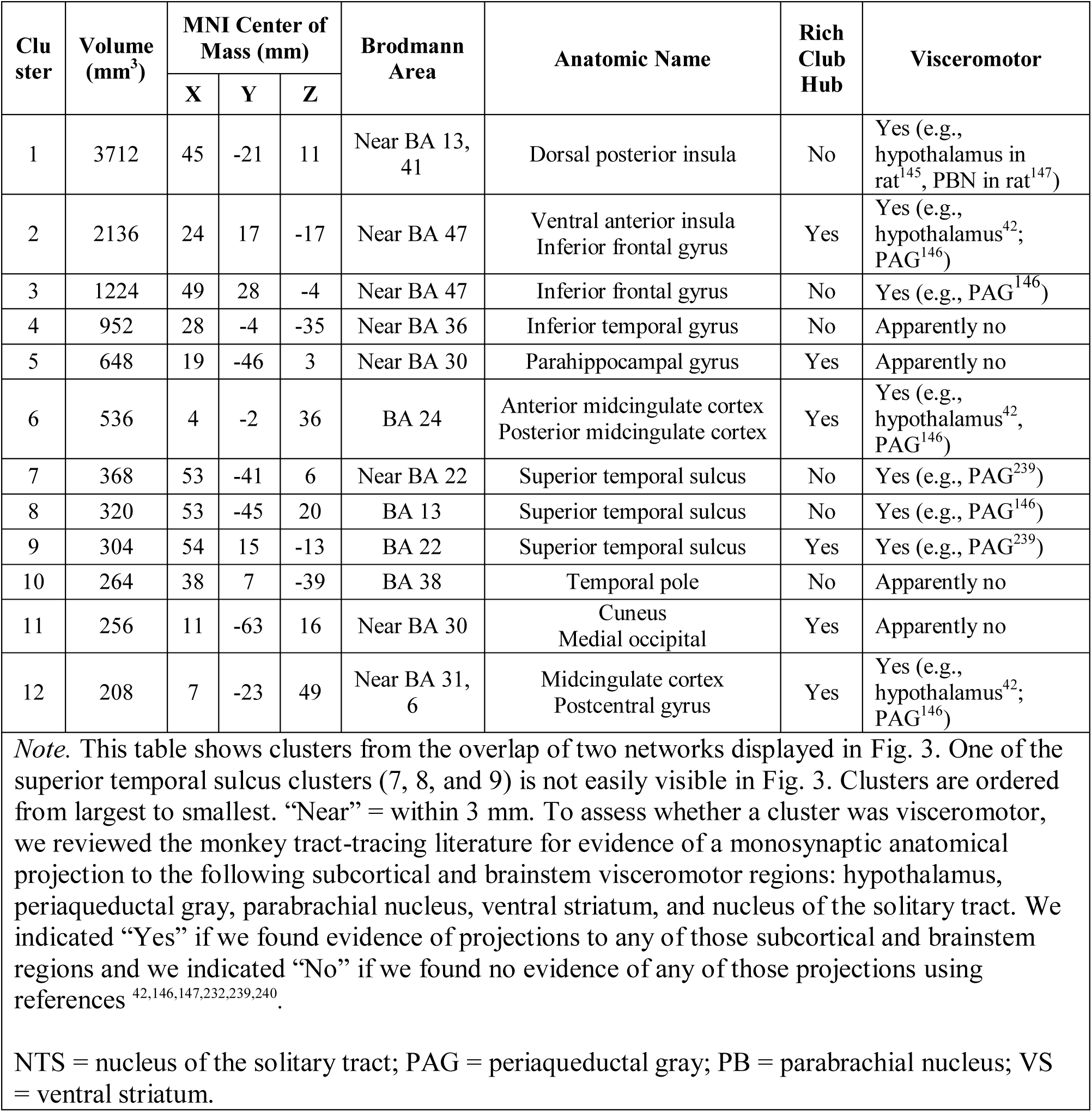
Connector hubs within the interoceptive system.

| Cluster | Volume (mm<sup>3</sup>) | MNI Center of Mass (mm) |  |  | Brodmann Area | Anatomic Name | Rich Club Hub | Visceromotor |
| --- | --- | --- | --- | --- | --- | --- | --- | --- |
|  |  | X | Y | Z |  |  |  |  |
| 1 | 3712 | 45 | -21 | 11 | Near BA 13, 41 | Dorsal posterior insula | No | Yes (e.g., hypothalamus in rat<sup>145</sup>, PBN in rat<sup>147</sup>) |
| 2 | 2136 | 24 | 17 | -17 | Near BA 47 | Ventral anterior insula<br>Inferior frontal gyrus | Yes | Yes (e.g., hypothalamus<sup>42</sup>, PAG<sup>146</sup>) |
| 3 | 1224 | 49 | 28 | -4 | Near BA 47 | Inferior frontal gyrus | No | Yes (e.g., PAG<sup>146</sup>) |
| 4 | 952 | 28 | -4 | -35 | Near BA 36 | Inferior temporal gyrus | No | Apparently no |
| 5 | 648 | 19 | -46 | 3 | Near BA 30 | Parahippocampal gyrus | Yes | Apparently no |
| 6 | 536 | 4 | -2 | 36 | BA 24 | Anterior midcingulate cortex<br>Posterior midcingulate cortex | Yes | Yes (e.g., hypothalamus<sup>42</sup>, PAG<sup>146</sup>) |
| 7 | 368 | 53 | -41 | 6 | Near BA 22 | Superior temporal sulcus | No | Yes (e.g., PAG<sup>239</sup>) |
| 8 | 320 | 53 | -45 | 20 | BA 13 | Superior temporal sulcus | No | Yes (e.g., PAG<sup>146</sup>) |
| 9 | 304 | 54 | 15 | -13 | BA 22 | Superior temporal sulcus | Yes | Yes (e.g., PAG<sup>239</sup>) |
| 10 | 264 | 38 | 7 | -39 | BA 38 | Temporal pole | No | Apparently no |
| 11 | 256 | 11 | -63 | 16 | Near BA 30 | Cuneus<br>Medial occipital | Yes | Apparently no |
| 12 | 208 | 7 | -23 | 49 | Near BA 31, 6 | Midcingulate cortex<br>Postcentral gyrus | Yes | Yes (e.g., hypothalamus<sup>42</sup>, PAG<sup>146</sup>) |

*Note.* This table shows clusters from the overlap of two networks displayed in Fig. 3. One of the superior temporal sulcus clusters (7, 8, and 9) is not easily visible in Fig. 3. Clusters are ordered from largest to smallest. “Near” = within 3 mm. To assess whether a cluster was visceromotor, we reviewed the monkey tract-tracing literature for evidence of a monosynaptic anatomical projection to the following subcortical and brainstem visceromotor regions: hypothalamus, periaqueductal gray, parabrachial nucleus, ventral striatum, and nucleus of the solitary tract. We indicated “Yes” if we found evidence of projections to any of those subcortical and brainstem regions and we indicated “No” if we found no evidence of any of those projections using references <sup>42,146,147,232,239,240</sup>.
NTS = nucleus of the solitary tract; PAG = periaqueductal gray; PB = parabrachial nucleus; VS = ventral striatum.

**Table S5.**
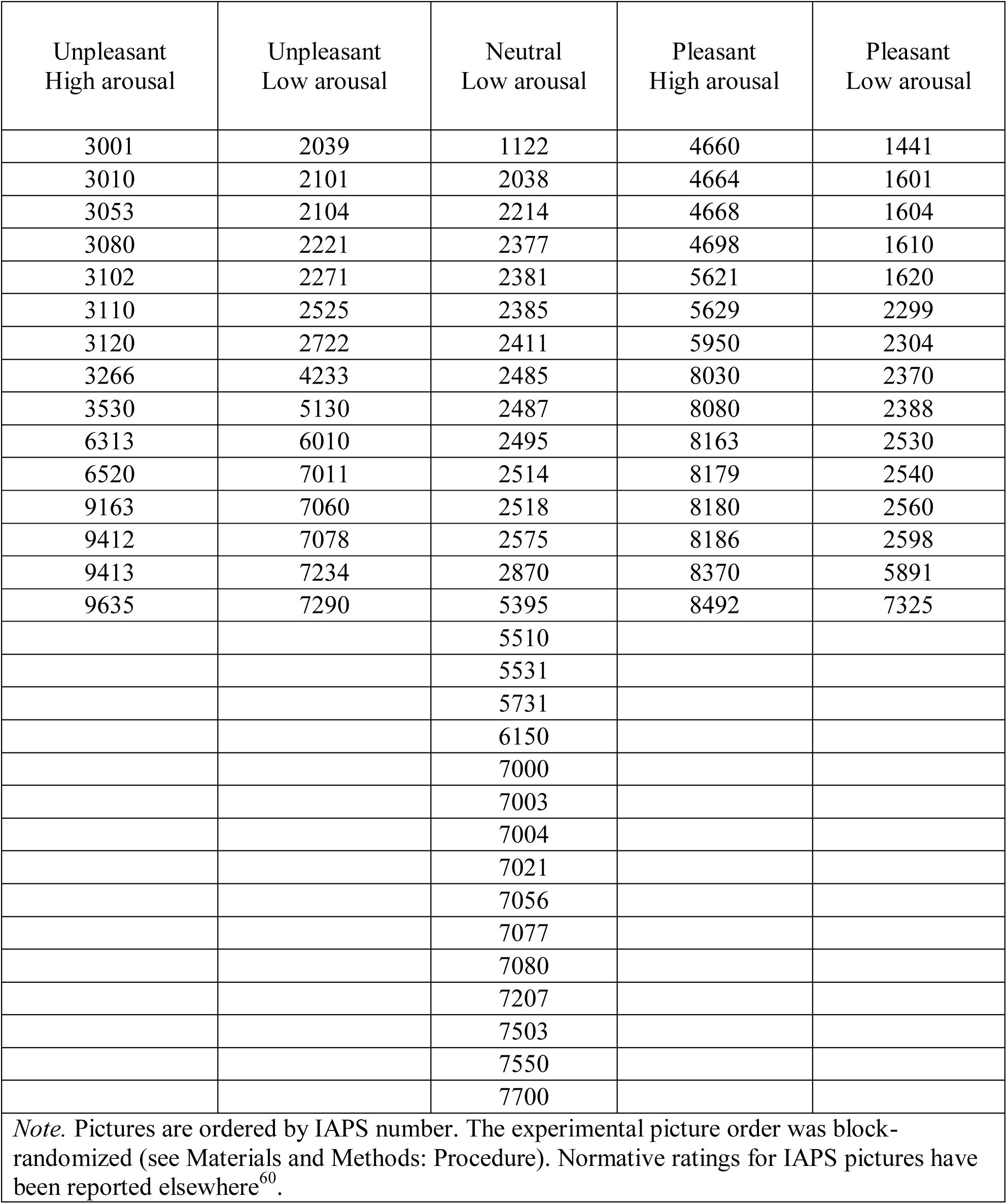
International affective picture system (IAPS) images used in the evocative pictures task.

| Unpleasant<br>High arousal | Unpleasant<br>Low arousal | Neutral<br>Low arousal | Pleasant<br>High arousal | Pleasant<br>Low arousal |
| --- | --- | --- | --- | --- |
| 3001 | 2039 | 1122 | 4660 | 1441 |
| 3010 | 2101 | 2038 | 4664 | 1601 |
| 3053 | 2104 | 2214 | 4668 | 1604 |
| 3080 | 2221 | 2377 | 4698 | 1610 |
| 3102 | 2271 | 2381 | 5621 | 1620 |
| 3110 | 2525 | 2385 | 5629 | 2299 |
| 3120 | 2722 | 2411 | 5950 | 2304 |
| 3266 | 4233 | 2485 | 8030 | 2370 |
| 3530 | 5130 | 2487 | 8080 | 2388 |
| 6313 | 6010 | 2495 | 8163 | 2530 |
| 6520 | 7011 | 2514 | 8179 | 2540 |
| 9163 | 7060 | 2518 | 8180 | 2560 |
| 9412 | 7078 | 2575 | 8186 | 2598 |
| 9413 | 7234 | 2870 | 8370 | 5891 |
| 9635 | 7290 | 5395 | 8492 | 7325 |
|  |  | 5510 |  |  |
|  |  | 5531 |  |  |
|  |  | 5731 |  |  |
|  |  | 6150 |  |  |
|  |  | 7000 |  |  |
|  |  | 7003 |  |  |
|  |  | 7004 |  |  |
|  |  | 7021 |  |  |
|  |  | 7056 |  |  |
|  |  | 7077 |  |  |
|  |  | 7080 |  |  |
|  |  | 7207 |  |  |
|  |  | 7503 |  |  |
|  |  | 7550 |  |  |
|  |  | 7700 |  |  |
| *Note.* Pictures are ordered by IAPS number. The experimental picture order was block-randomized (see Materials and Methods: Procedure). Normative ratings for IAPS pictures have been reported elsewhere<sup>60</sup>. |  |  |  |  |

**Fig. S1.**
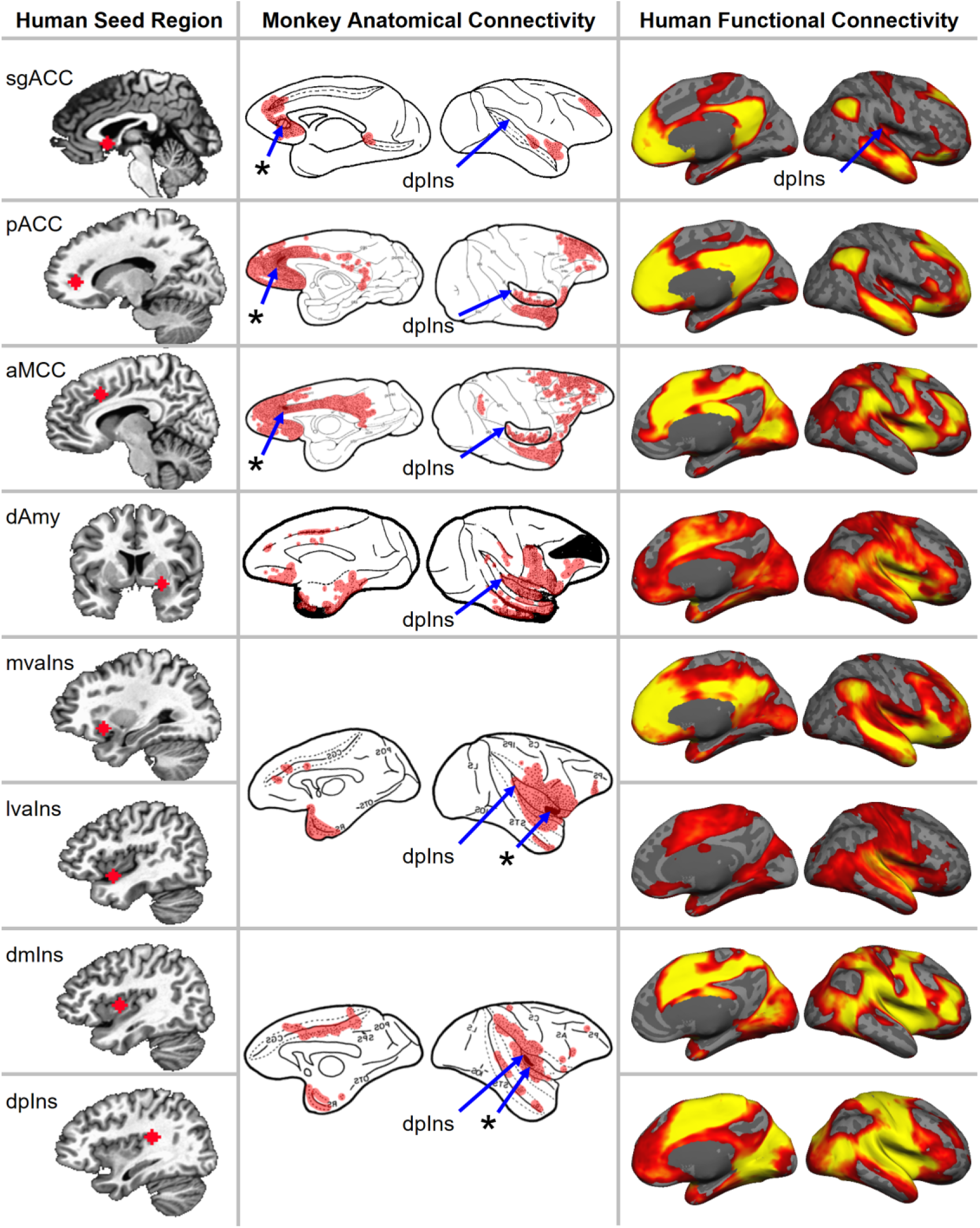
Replication of the interoceptive system connecting the cortical and amygdalar visceromotor regions and primary interoceptive regions shown in Fig. 2 using the replication sample (*N* = 270). The color map ranges from *p* = 10^-5^ in red to *p* = 10^-40^ in yellow, uncorrected and on a log scale.

**Fig. S2.**
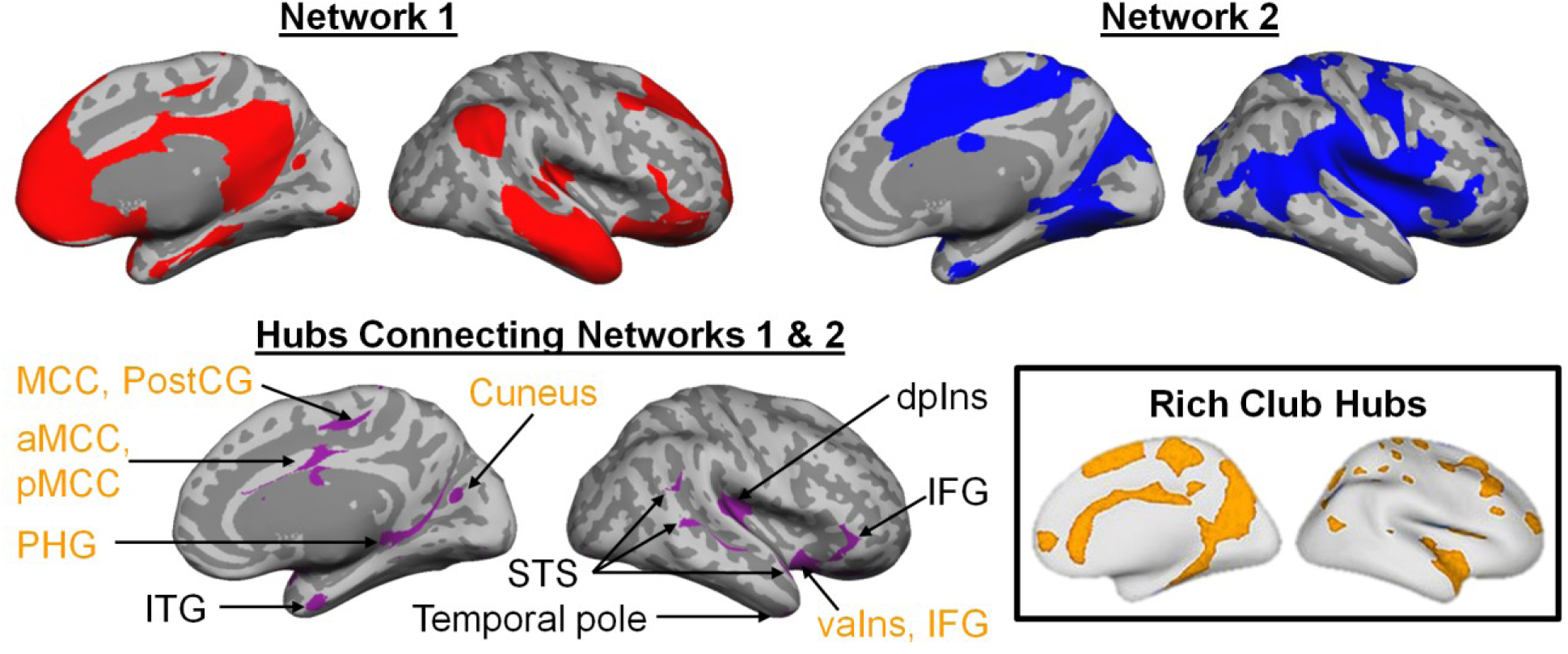
Replication of the interoceptive system shown in Fig. 3 using the replication sample (N = 270). Intrinsic connectivity maps binarized at *p* < 10^-5^ uncorrected. Rich club hubs figure adapted with permission from van den Heuval & Sporns (2013)^84^.

**Fig. S3.**
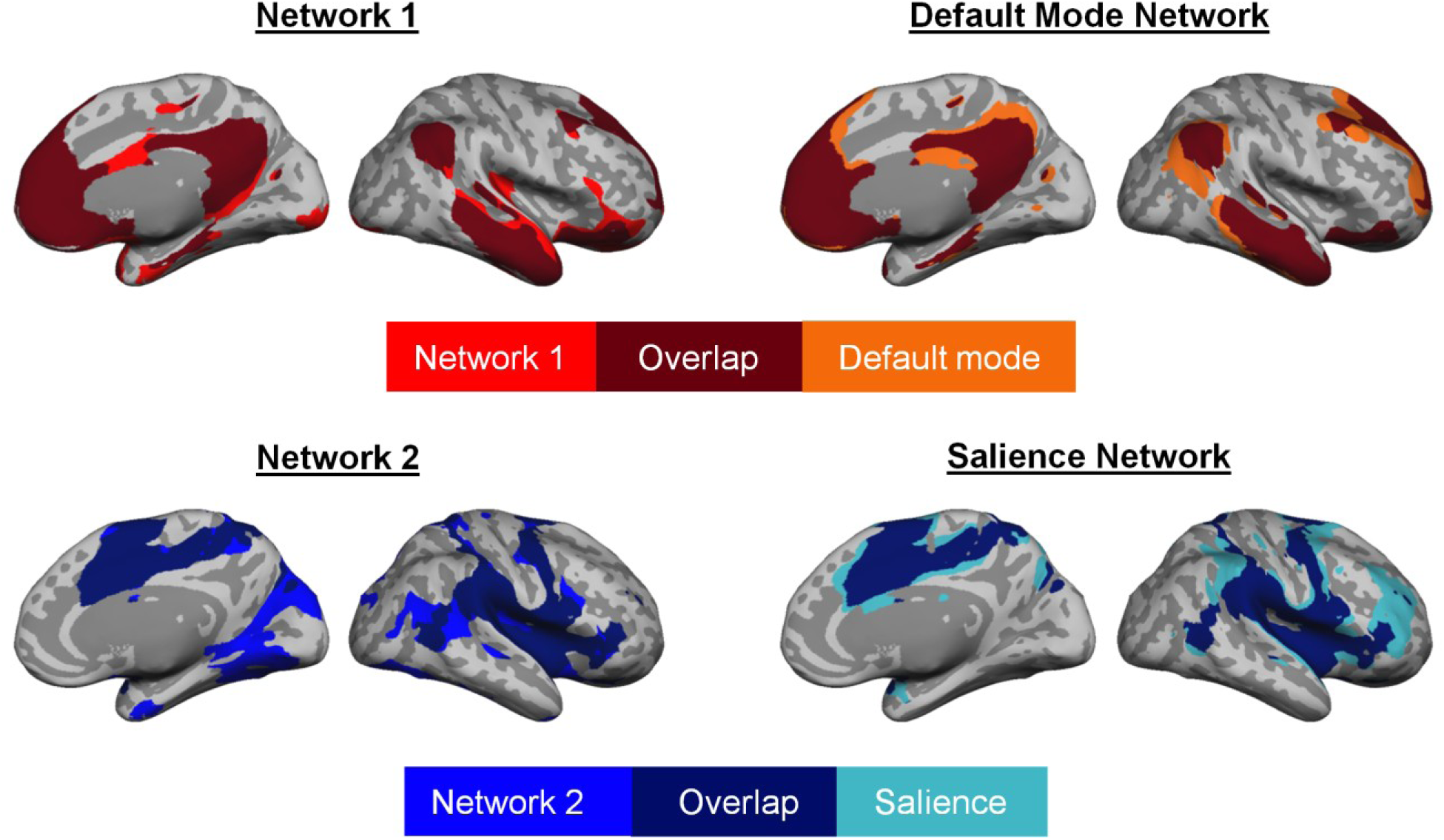
Correspondence between allostatic/interoceptive system networks (binarized at *p <* 10^-5^ uncorrected) and established default mode and salience networks in the discovery sample (*N* = 280). The masks of the established networks were computed based on Yeo et al. (2011)^54^ using a traditional volume-based approac^215,241^on a sample of 150 subjects (see Methods for description of subjects). The correspondence between the interoceptive system networks and established networks was replicated in a second sample of *N* = 270 (Fig. S4).

**Fig. S4.**
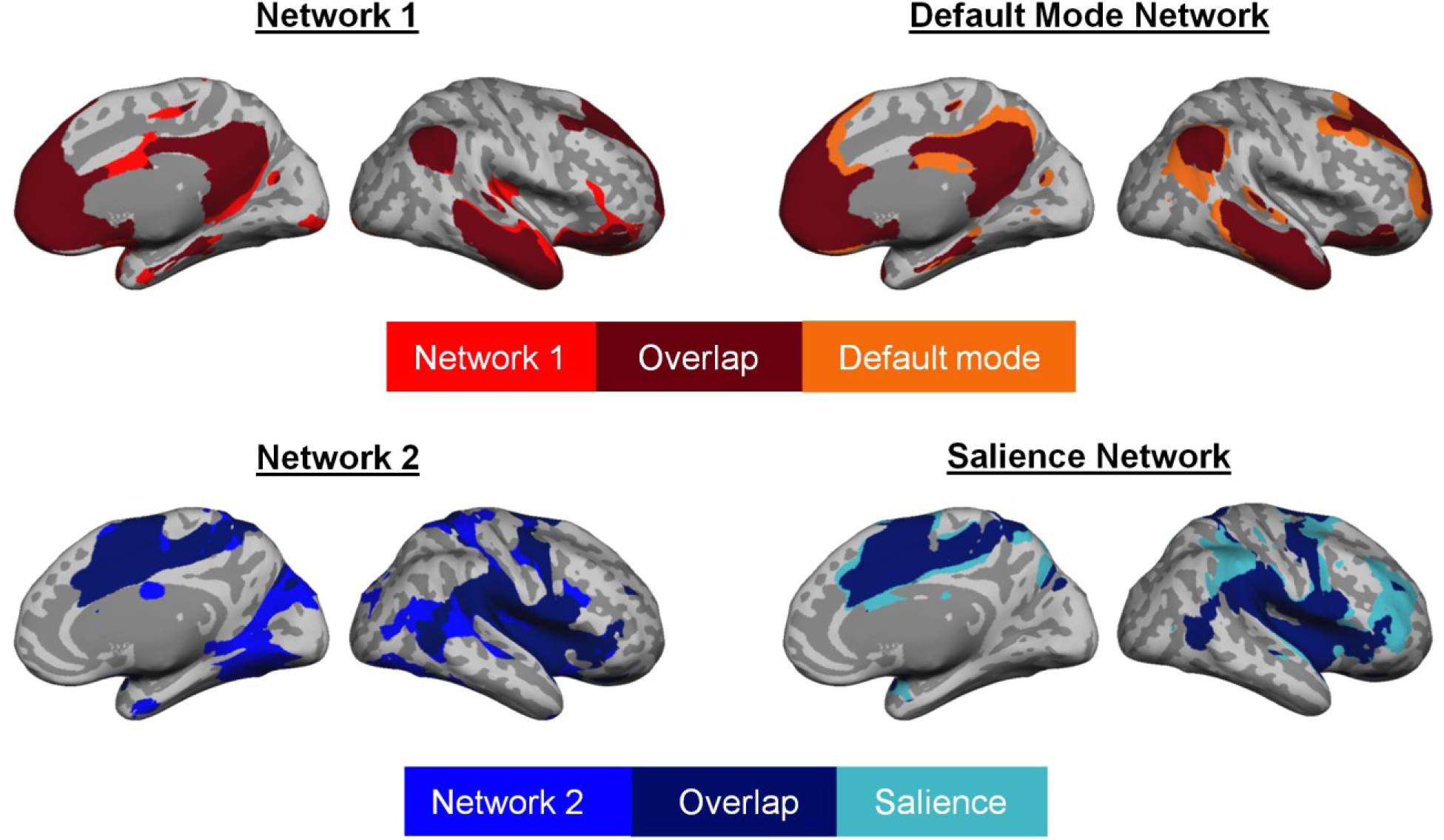
Replication of correspondence between interoceptive system networks (binarized at *p* < 10^-5^ uncorrected) and established default mode and salience networks shown in Fig. S3 using the replication sample (*N* = 270). The masks of the established networks were computed based on Yeo et al.^54^ using a traditional volume-based approac^215,241^ on a sample of 150 subjects (see Methods for description of subjects).

**Fig. S5.**
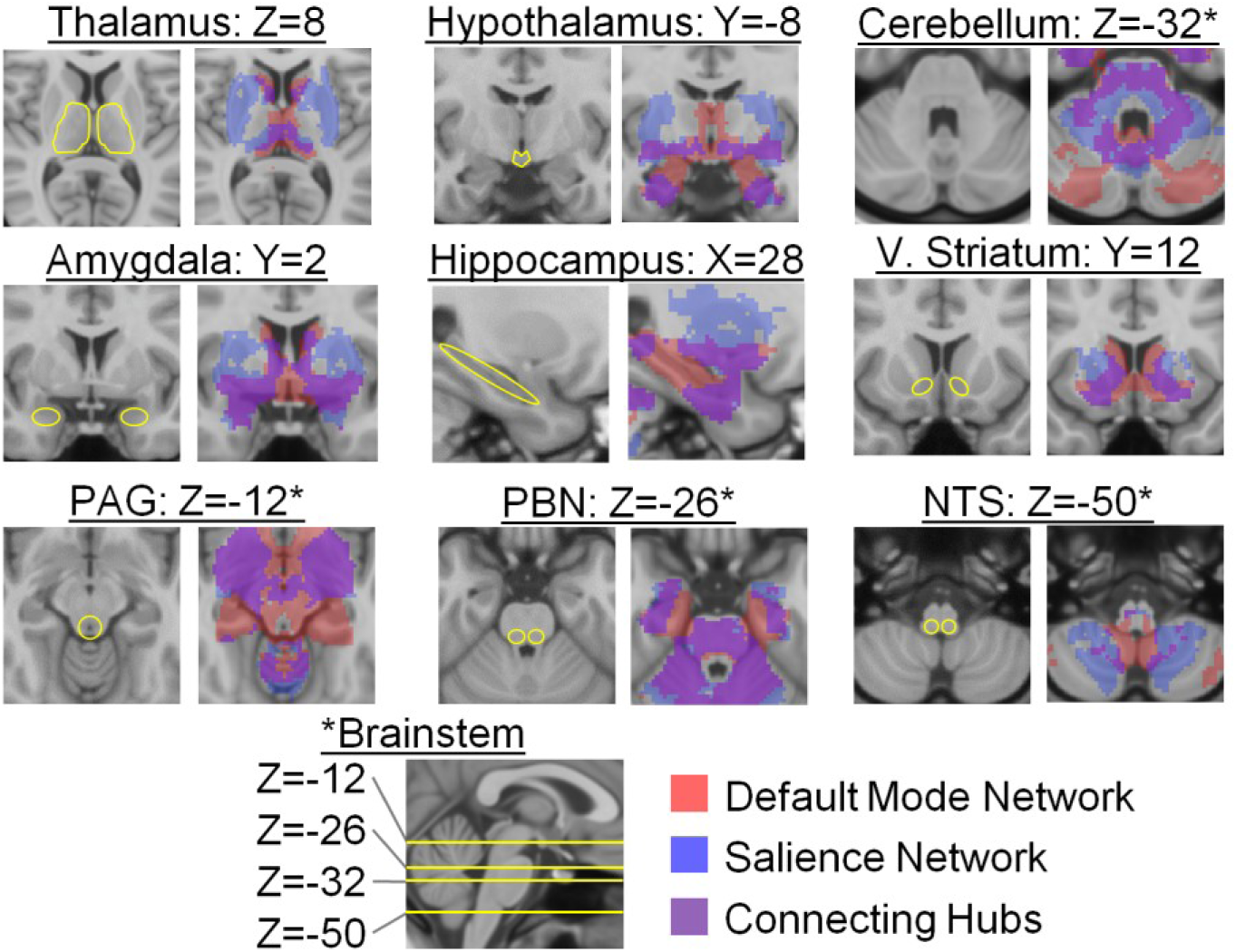
Replication of subcortical connectivity of the two intrinsic networks within the interoceptive system in Fig. 4 using the replication sample (*N* = 270; *p* < 10^-5^ uncorrected, one-sample two-tailed t-test). PAG = periaqueductal gray; PBN = parabrachial nucleus; V. Striatum = ventral striatum; NTS = nucleus of the solitary tract.

**Fig. S6.**
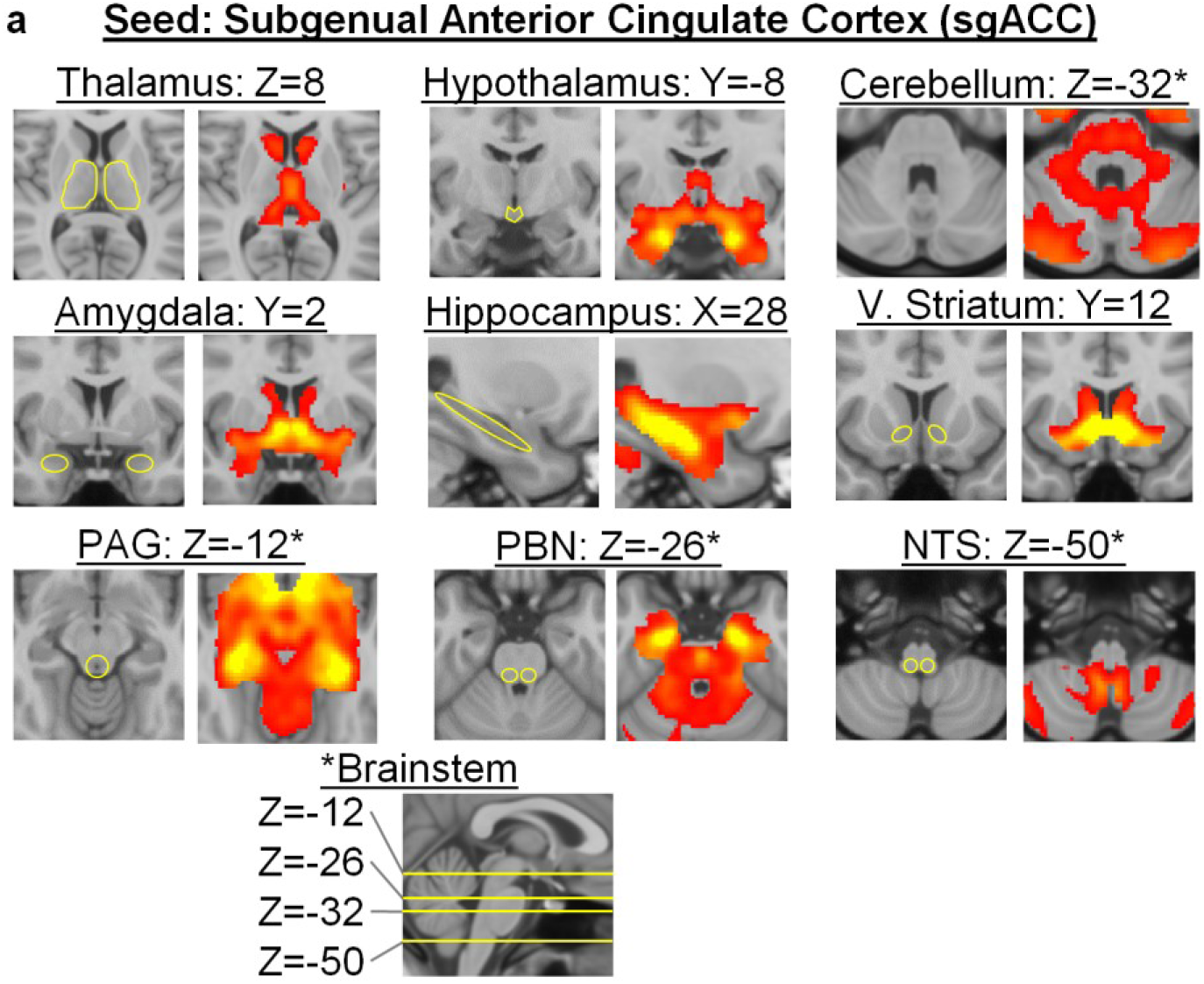
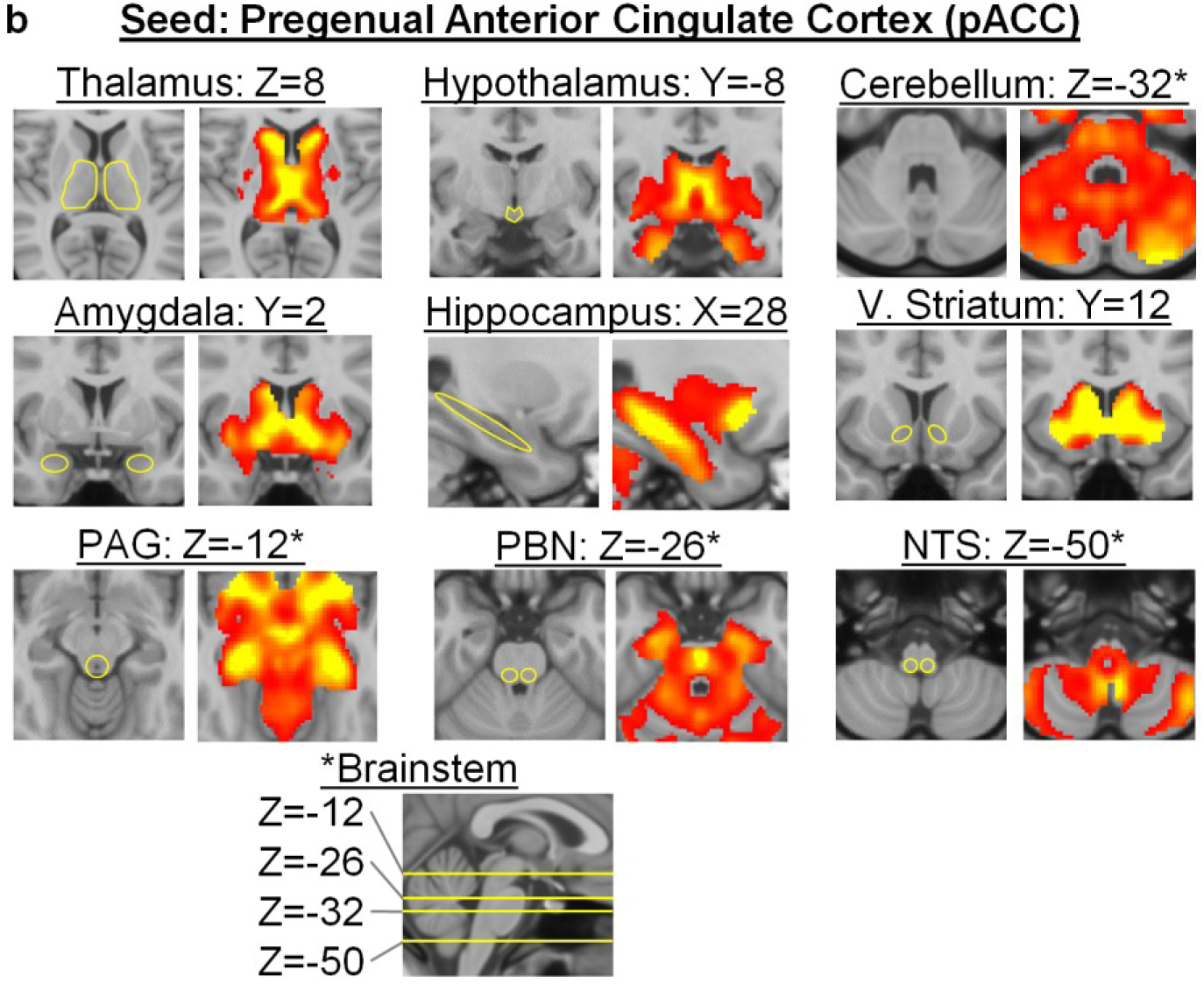
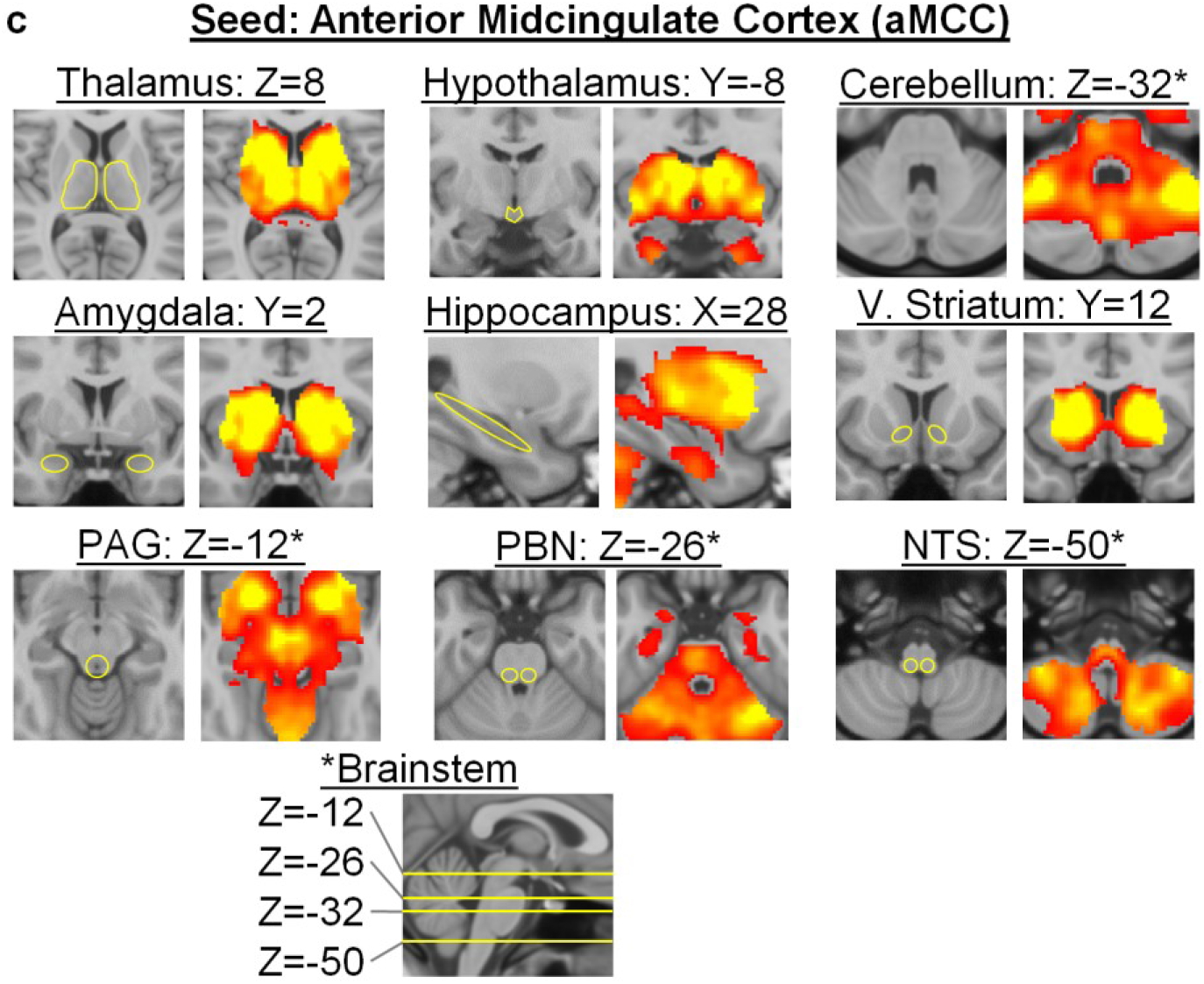
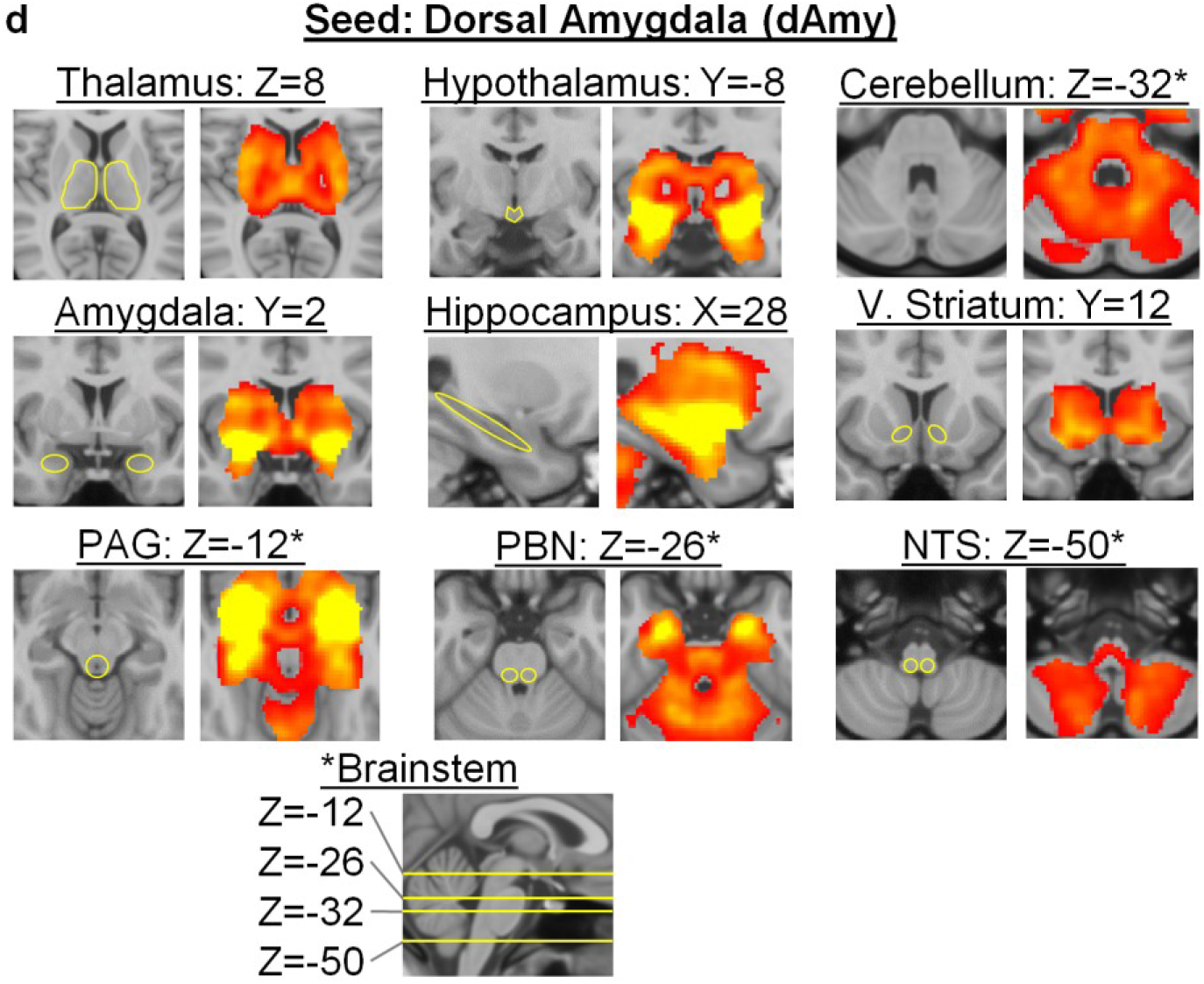
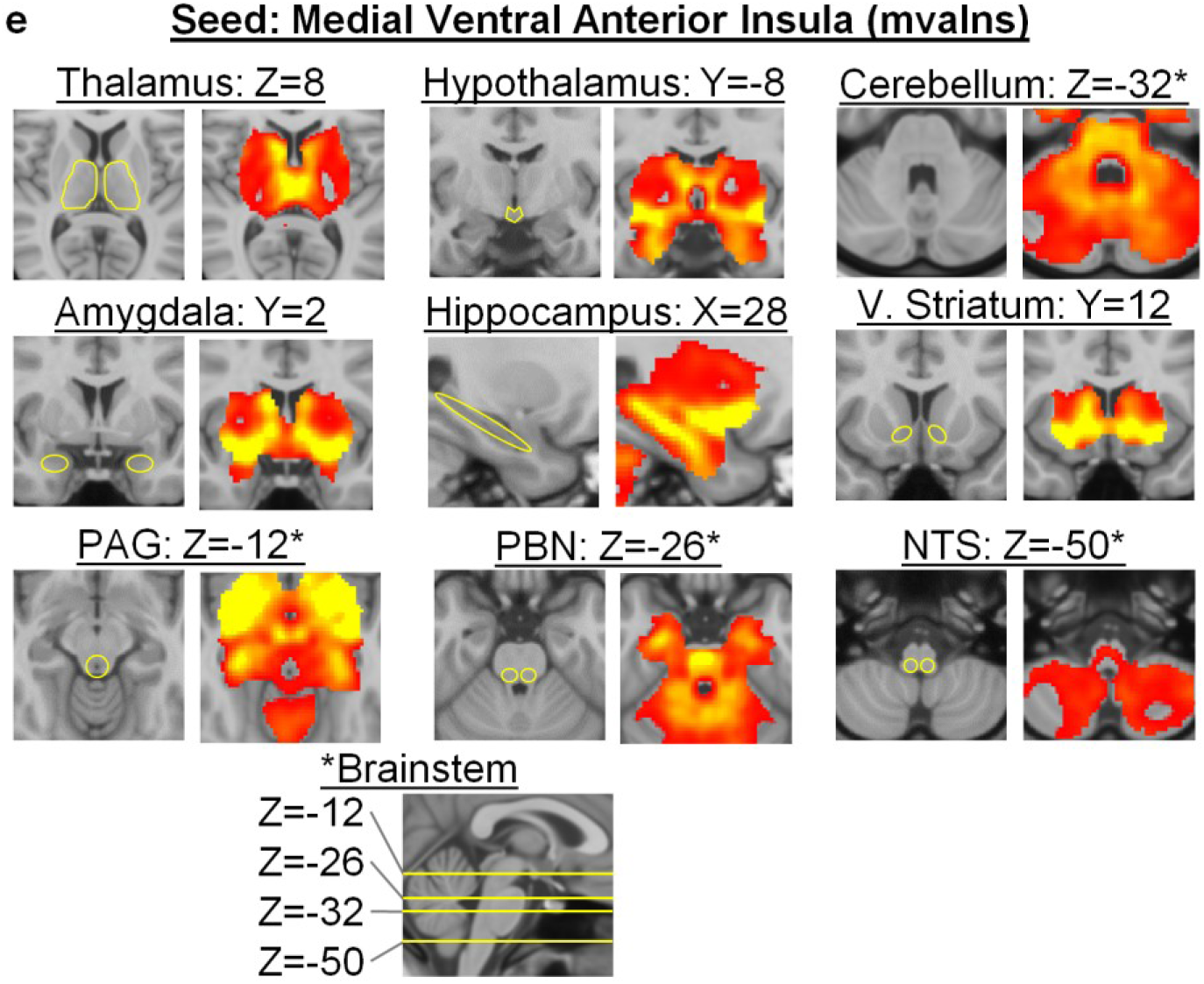
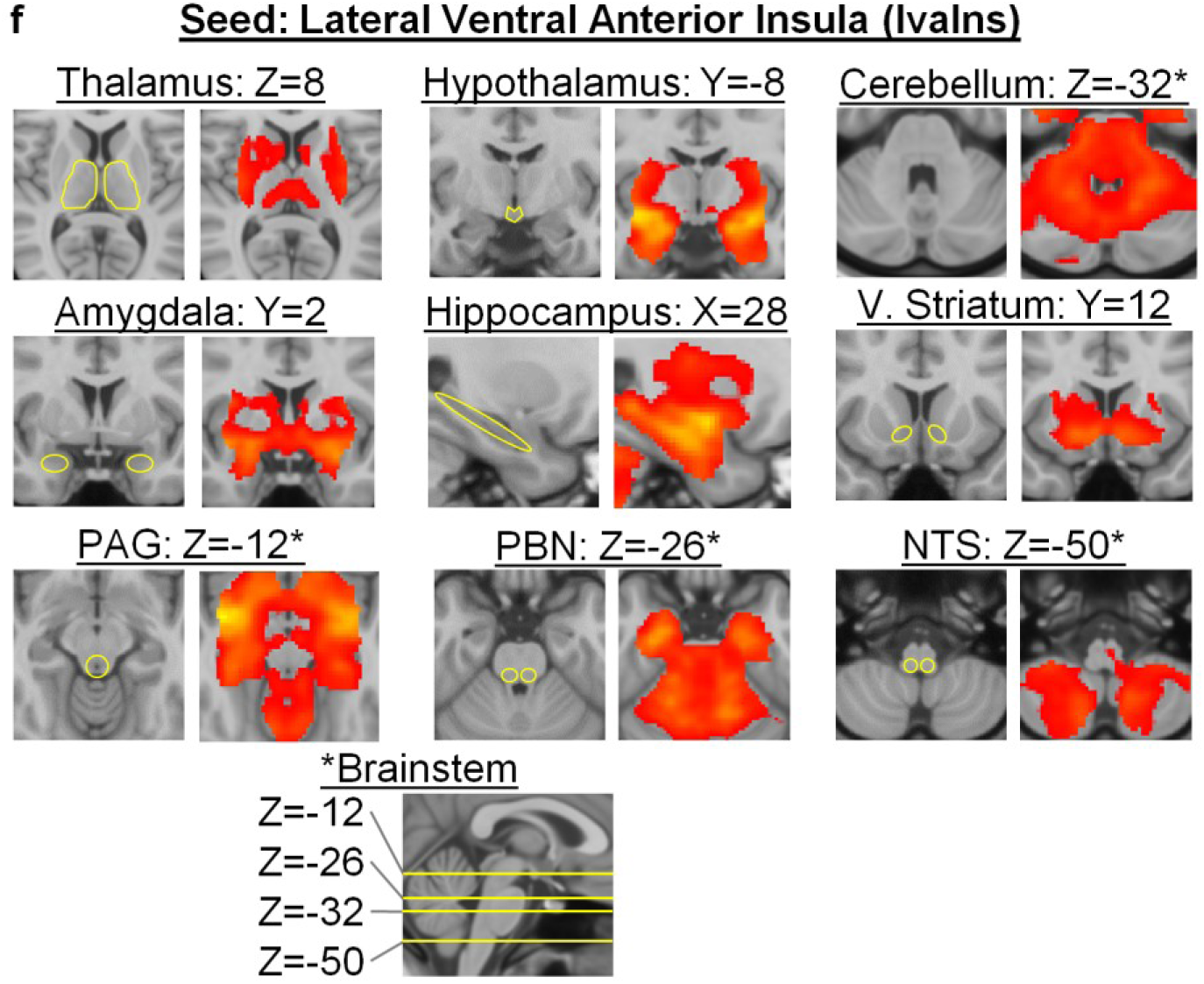
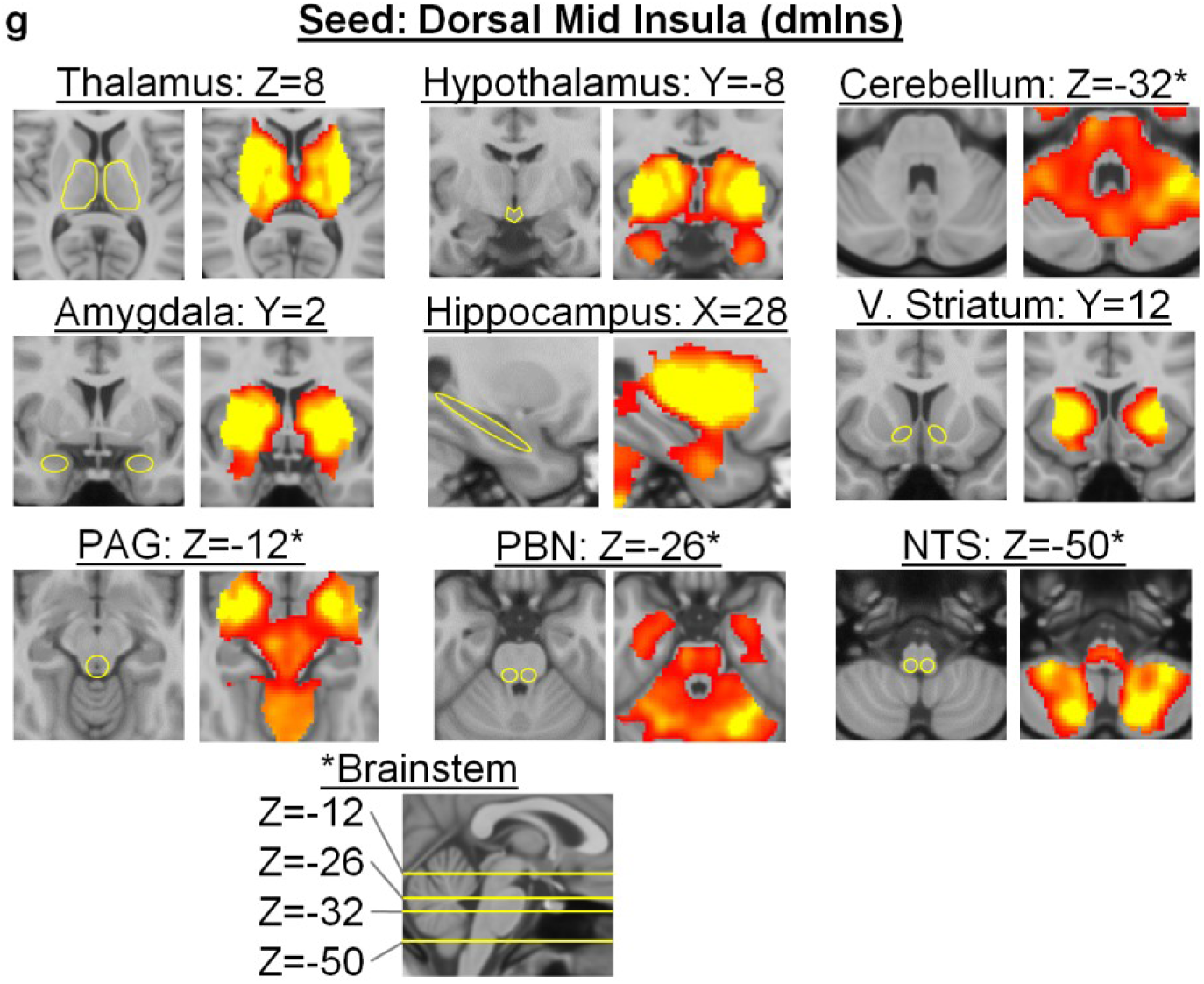
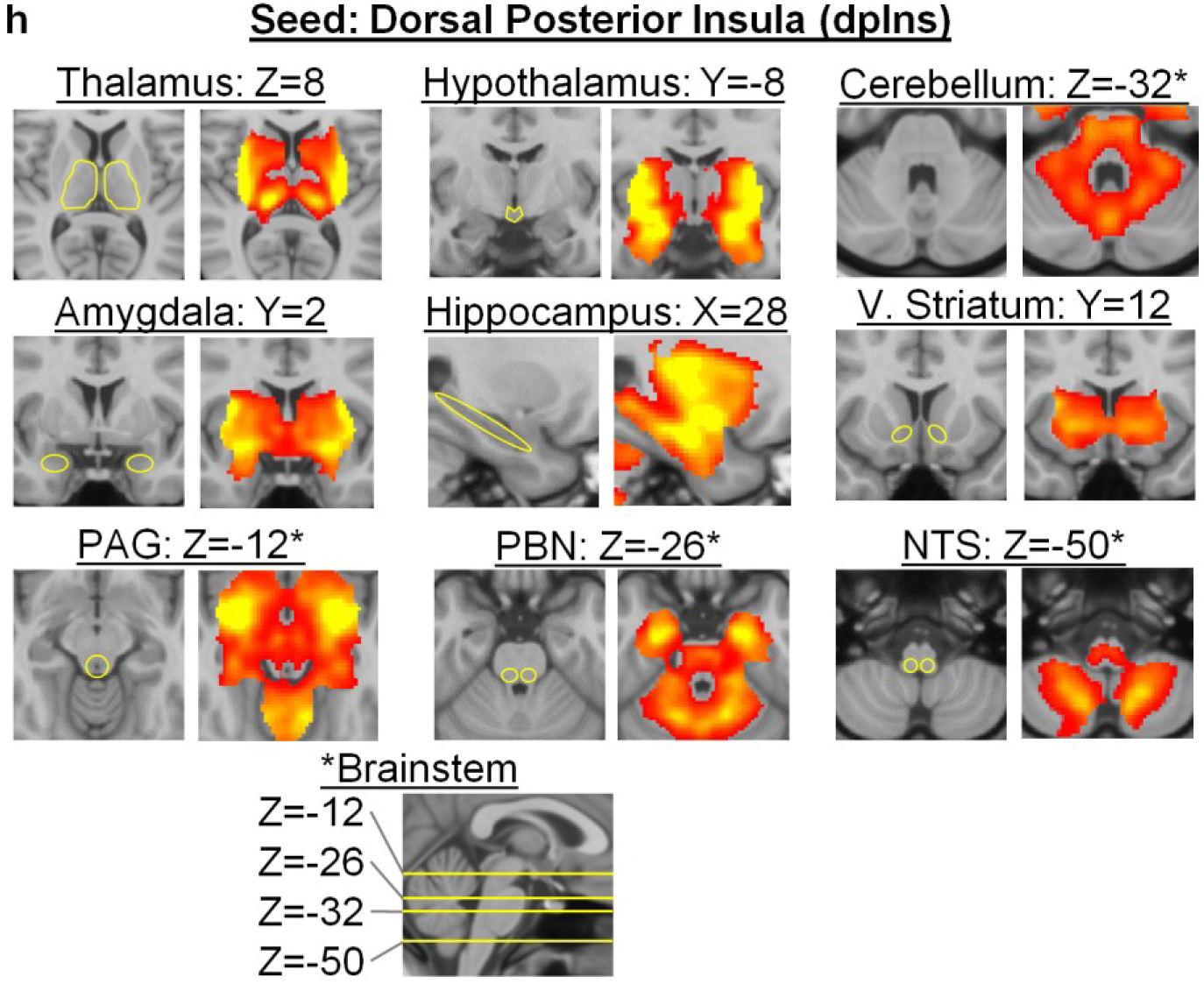
Subcortical connectivity of each of the eight cortical and amygdalar seeds used to delineate the interoceptive system using the discovery sample (*N* = 280). The color scale ranges from *p* = 0.05 in red to *p* = 10^-50^ in yellow, uncorrected and on a log scale. (a) Seed at subgenual anterior cingulate cortex (sgACC). (b) Seed at pregenual anterior cingulate cortex (pACC). (c) Seed at anterior midcingulate cortex (aMCC). (d) Seed at dorsal amygdala (dAmy). (e) Seed at medial ventral anterior insula (mvaIns). (f) Seed at lateral ventral anterior insula (lvaIns). (g) Seed at dorsal mid insula (dmIns). (h) Seed at dorsal posterior insula (dpIns). PAG = periaqueductal gray; PBN = parabrachial nucleus; V. Striatum = ventral striatum; NTS = nucleus of the solitary tract.

**Fig. S7.**
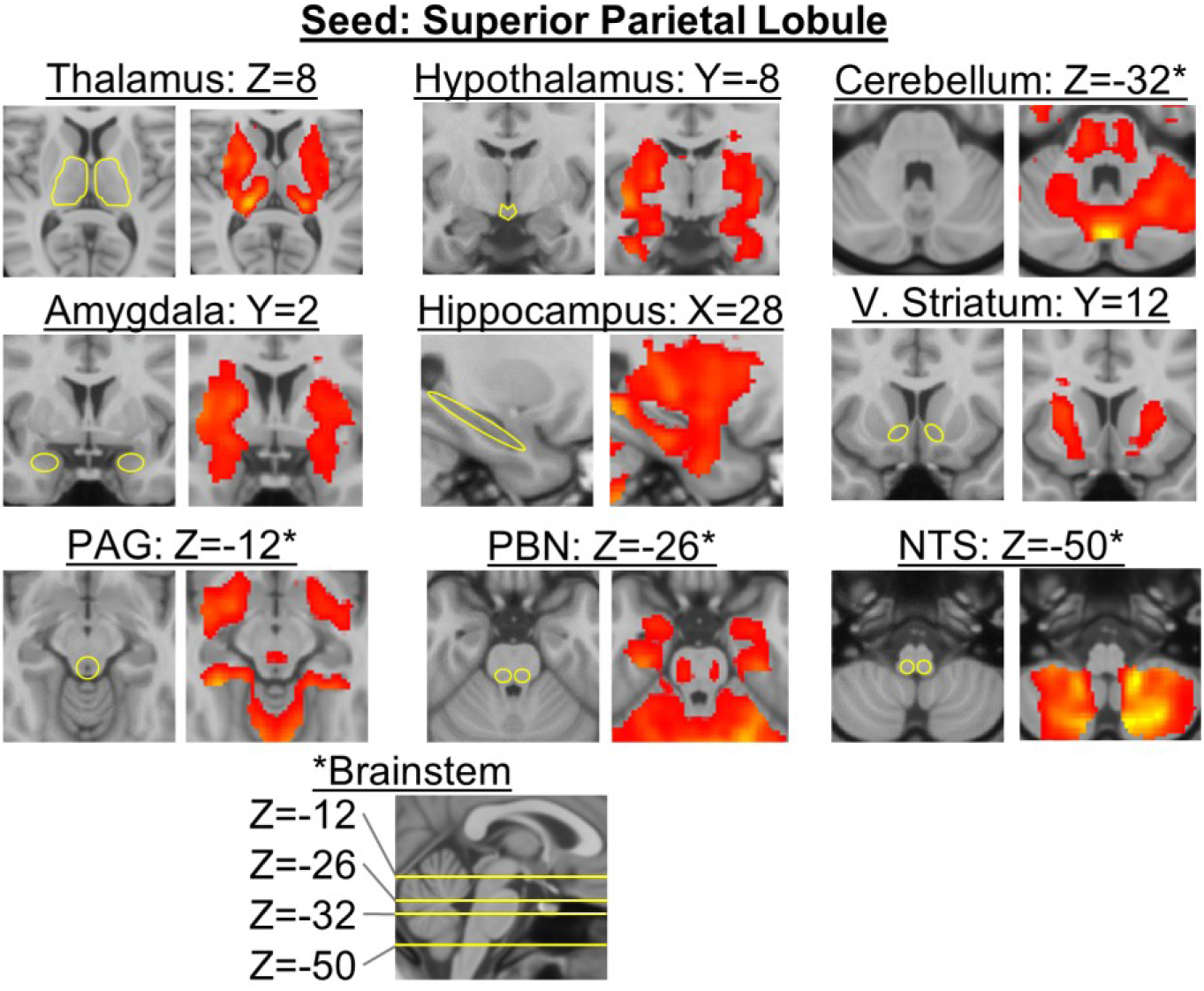
Subcortical connectivity of the superior parietal lobule does not involve non-cortical targets important for allostasis (hypothalamus, PAG, PBN, NTS), suggesting a degree of specificity in the functional connections between cortical regions in the allostatic/interoceptive system and subcortical regions that support allostasis. The seed coordinates are MNI X, Y, Z = 46, 12, 28. These results are from the discovery sample (*N* = 280). Identical conclusions were obtained using results for the replication sample (*N* = 270). The color scale ranges from *p* = 0.05 in red to *p* = 10^-50^ in yellow, uncorrected and on a log scale. PAG = periaqueductal gray; PBN = parabrachial nucleus; V. Striatum = ventral striatum; NTS = nucleus of the solitary tract.

**Fig. S8.**
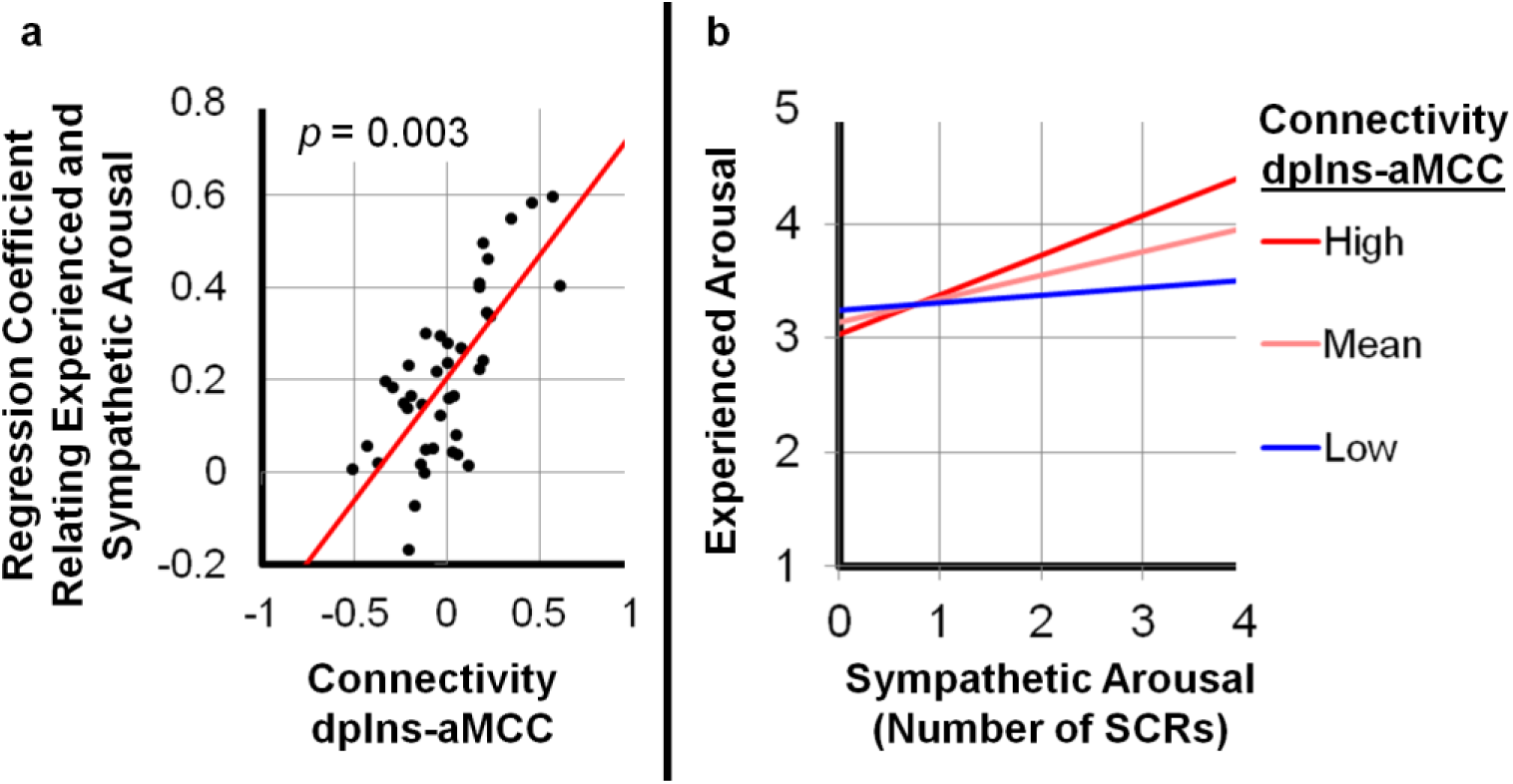
The strength of intrinsic connectivity within the allostatic/interoceptive system predicts the degree of interoception during allostatic fluctuations. (a) Each data point corresponds to one participant (*N* = 41). Interoceptive ability is represented on the y-axis as the unstandardized linear regression coefficient *B* for the relationship between sympathetic nervous system arousal (i.e., the number of skin conductance responses) and the intensity of experienced arousal (rated on a 1 to 5 scale) while participants viewed evocative pictures. The strength of intrinsic connectivity between a primary interoceptive region (dorsal posterior insula, or dpIns) and a key visceromotor control region (the anterior portion of the midcingulate cortex, or aMCC) is represented on the x-axis as the Fisher’s z transformation of the Pearson’s correlation coefficient of the two BOLD time courses (the red line shows the regression line with coefficient *B* = 0.56; *p* = 0.003; multi-level modeling regression analysis). (b) The relationship between experienced arousal and the number of skin conductance responses (as an index of sympathetic nervous system arousal) is moderated by intrinsic connectivity strength. “High” or “Low” corresponds to one SD above or below the mean connectivity across participants, respectively (mean 0.1, SD 0.23). dpIns = dorsal posterior insula; aMCC = anterior midcingulate cortex; SCRs = skin conductance responses.

## Endnotes

1 We included the dAmy in our system because its central nucleus is known to have key visceromotor functions (for a review, see ^139^); the dAmy, being a subcortical region, does not have a laminar structure, but there are connections between the amygdala and primary interoceptive cortex (dmIns/dpIns; e.g., ^59,140,141^) that are predicted by the EPIC model (using Barbas’s structural model of information flow within the cortex). Similarly, the anterior cingulate cortex (ACC), a key limbic visceromotor region, is connected with the amygdala in a pattern consistent with the EPIC model hypothesis that the ACC sends visceromotor prediction signals to the central nucleus (the ACC primarily sends output from its deep layers and receives input from the amygdala in its upper layers^142^). Currently, there are insufficient data to test the EPIC model hypothesis that amygdala projections terminate in the upper layers of dmIns/dpIns and receives inputs from its deep layers, as these data are not available in prior tract-tracing studies involving the insula and amygdala (e.g., ^59,140,141^).

## References and Notes

1 Sepulcre, J., Sabuncu, M. R., Yeo, T. B., Liu, H. & Johnson, K. A. Stepwise connectivity of the modal cortex reveals the multimodal organization of the human brain. J. Neurosci. 32, 10649–10661 (2012).

2 Rao, R. P. & Ballard, D. H. Predictive coding in the visual cortex: A functional interpretation of some extra-classical receptive-field effects. Nat. Neurosci. 2, 79–87 (1999).

3 Chennu, S. et al. Expectation and attention in hierarchical auditory prediction. J. Neurosci. 33, 11194–11205 (2013).

4 Shipp, S. The importance of being agranular: A comparative account of visual and motor cortex. Philos. Trans. R. Soc. Lond. B Biol. Sci. 360, 797–814 (2005).

5 Zelano, C., Mohanty, A. & Gottfried, J. A. Olfactory predictive codes and stimulus templates in piriform cortex. Neuron 72, 178–187 (2011).

6 Kusumoto-Yoshida, I., Liu, H., Chen, B. T., Fontanini, A. & Bonci, A. Central role for the insular cortex in mediating conditioned responses to anticipatory cues. Proc. Natl. Acad. Sci. U. S. A. 112, 1190–1195 (2015).

7 Adams, R. A., Shipp, S. & Friston, K. J. Predictions not commands: Active inference in the motor system. Brain structure & function 218, 611–643 (2013).

8 Clark, A. Whatever next? Predictive brains, situated agents, and the future of cognitive science. Behav. Brain Sci. 36, 181–204 (2013).

9 Friston, K. The free-energy principle: A unified brain theory? Nat. Rev. Neurosci. 11, 127–138 (2010).

10 Pezzulo, G., Rigoli, F. & Friston, K. Active inference, homeostatic regulation and adaptive behavioural control. Prog. Neurobiol. 134, 17–35 (2015).

11 Barrett, L. F. & Simmons, W. K. Interoceptive predictions in the brain. Nat. Rev. Neurosci. 16, 419–429 (2015).

12 Chanes, L. & Barrett, L. F. Redefining the role of limbic areas in cortical processing. Trends in cognitive sciences 20, 96–106 (2016).

13 Seth, A. K. Interoceptive inference, emotion, and the embodied self. Trends in cognitive sciences 17, 565–573 (2013).

14 Seth, A. K., Suzuki, K. & Critchley, H. D. An interoceptive predictive coding model of conscious presence. Front. Psychol. 2, 1–16 (2012).

15 Gu, X. & FitzGerald, T. H. Interoceptive inference: Homeostasis and decision-making. Trends in cognitive sciences 18, 269–270 (2014).

16 Craig, A. D. How do you feel? Interoception: The sense of the physiological condition of the body. Nat. Rev. Neurosci. 3, 655–666 (2002).

17 Craig, B. How do you feel?: An interoceptive moment with your neurobiological self. (Princeton University Press, 2014).

18 Sherrington, C. in Textbook of physiology (ed EA Schäfer) 920–1001 (Pentland, 1900).

19 Sterling, P. & Laughlin, S. Principles of neural design. (MIT Press, 2015).

20 Sterling, P. Allostasis: A model of predictive regulation. Physiol. Behav. 106, 5–15 (2012).

21 McEwen, B. S. & Stellar, E. Stress and the individual. Mechanisms leading to disease. Arch. Intern. Med. 153, 2093–2101 (1993).

22 Barrett, L. F. How emotions are made: The secret life of the brain. (Houghton-Mifflin-Harcourt, 2017).

23 Barrett, L. F. The theory of constructed emotion: An active inference account of interoception and categorization. Soc. Cogn. Affect. Neurosci. (in press).

24 Barrett, L. F., Quigley, K. S. & Hamilton, P. An active inference theory of allostasis and interoception in depression. Phil. Trans. R. Soc. B. (in press).

25 Damasio, A. & Carvalho, G. B. The nature of feelings: Evolutionary and neurobiological origins. Nat. Rev. Neurosci. 14, 143–152 (2013).

26 Critchley, H. D. & Harrison, N. A. Visceral influences on brain and behavior. Neuron 77, 624–638 (2013).

27 Barrett, L. F. & Satpute, A. B. Large-scale brain networks in affective and social neuroscience: Towards an integrative functional architecture of the brain. Curr. Opin. Neurobiol. 23, 361–372 (2013).

28 Anderson, M. L. After phrenology neural reuse and the interactive brain. (MIT Press, 2014).

29 Yeo, B. T. et al. Functional specialization and flexibility in human association cortex. Cereb. Cortex 25, 3654–3672 (2015).

30 Barbas, H. & Rempel-Clower, N. Cortical structure predicts the pattern of corticocortical connections. Cereb. Cortex 7, 635–646 (1997).

31 Barbas, H. General cortical and special prefrontal connections: Principles from structure to function. Annu. Rev. Neurosci. 38, 269–289 (2015).

32 Rempel-Clower, N. L. & Barbas, H. The laminar pattern of connections between prefrontal and anterior temporal cortices in the rhesus monkey is related to cortical structure and function. Cereb. Cortex 10, 851–865 (2000).

33 Medalla, M. & Barbas, H. Specialized prefrontal “auditory fields”: Organization of primate prefrontal-temporal pathways. Front. Neurosci. 8, 77 (2014).

34 Medalla, M. & Barbas, H. Diversity of laminar connections linking periarcuate and lateral intraparietal areas depends on cortical structure. Eur. J. Neurosci. 23, 161–179 (2006).

35 Hilgetag, C. C. & Grant, S. Cytoarchitectural differences are a key determinant of laminar projection origins in the visual cortex. Neuroimage 51, 1006–1017 (2010).

36 Goulas, A., Uylings, H. B. & Stiers, P. Mapping the hierarchical layout of the structural network of the macaque prefrontal cortex. Cereb. Cortex 24, 1178–1194 (2014).

37 Bastos, A. M. et al. Canonical microcircuits for predictive coding. Neuron 76, 695–711 (2012).

38 Adams, R. A., Stephan, K. E., Brown, H. R., Frith, C. D. & Friston, K. J. The computational anatomy of psychosis. Front Psychiatry 4, 47 (2013).

39 Shipp, S., Adams, R. A. & Friston, K. J. Reflections on agranular architecture: Predictive coding in the motor cortex. Trends Neurosci. 36, 706–716 (2013).

40 Weston, C. S. Another major function of the anterior cingulate cortex: The representation of requirements. Neurosci. Biobehav. Rev. 36, 90–110 (2012).

41 Vogt, B. A. Pain and emotion interactions in subregions of the cingulate gyrus. Nat. Rev. Neurosci. 6, 533–544 (2005).

42 Ongur, D., An, X. & Price, J. L. Prefrontal cortical projections to the hypothalamus in macaque monkeys. J. Comp. Neurol. 401, 480–505 (1998).

43 Vogt, B. A., Vogt, L., Farber, N. B. & Bush, G. Architecture and neurocytology of monkey cingulate gyrus. J. Comp. Neurol. 485, 218–239 (2005).

44 Nieuwenhuys, R. The insular cortex: A review. Prog. Brain Res. 195, 123–163 (2012).

45 Avery, J. A. et al. A common gustatory and interoceptive representation in the human mid-insula. Hum. Brain Mapp. 36, 2996–3006 (2015).

46 Hutchison, R. M. & Everling, S. Monkey in the middle: Why non-human primates are needed to bridge the gap in resting-state investigations. Front. Neuroanat. 6, 29 (2012).

47 Deco, G., Jirsa, V. K. & McIntosh, A. R. Emerging concepts for the dynamical organization of resting-state activity in the brain. Nat. Rev. Neurosci. 12, 43–56 (2011).

48 Cohen, A. L. et al. Defining functional areas in individual human brains using resting functional connectivity mri. Neuroimage 41, 45–57 (2008).

49 Raichle, M. E. The brain's default mode network. Annu. Rev. Neurosci. 38, 433–447 (2015).

50 Touroutoglou, A., Hollenbeck, M., Dickerson, B. C. & Barrett, L. F. Dissociable large-scale networks anchored in the right anterior insula subserve affective experience and attention. Neuroimage 60, 1947–1958 (2012).

51 Seeley, W. W. et al. Dissociable intrinsic connectivity networks for salience processing and executive control. J. Neurosci. 27, 2349–2356 (2007).

52 Dosenbach, N. U., Fair, D. A., Cohen, A. L., Schlaggar, B. L. & Petersen, S. E. A dual-networks architecture of top-down control. Trends in cognitive sciences 12, 99–105 (2008).

53 Corbetta, M. & Shulman, G. L. Control of goal-directed and stimulus-driven attention in the brain. Nat. Rev. Neurosci. 3, 201–215 (2002).

54 Yeo, B. T. et al. The organization of the human cerebral cortex estimated by intrinsic functional connectivity. J. Neurophysiol. 106, 1125–1165 (2011).

55 Nieuwenhuys, R., Voogd, J. & Huijzen, C. v. in The human central nervous system Ch. 8, 253–279 (Springer, 2008).

56 Brooks, J. C., Faull, O. K., Pattinson, K. T. & Jenkinson, M. Physiological noise in brainstem fmri. Front. Hum. Neurosci. 7, 623 (2013).

57 Craig, A. D. How do you feel--now? The anterior insula and human awareness. Nat. Rev. Neurosci. 10, 59–70 (2009).

58 Ongur, D. & Price, J. L. The organization of networks within the orbital and medial prefrontal cortex of rats, monkeys and humans. Cereb. Cortex 10, 206–219 (2000).

59 Hoistad, M. & Barbas, H. Sequence of information processing for emotions through pathways linking temporal and insular cortices with the amygdala. Neuroimage 40, 1016–1033 (2008).

60 Lang, P. J., Bradley, M. M. & Cuthbert, B. N. International affective picture system (iaps): Affective ratings of pictures and instruction manual. Technical Report A-8. University of Florida, Gainesville, FL. (2008).

61 Moriguchi, Y. et al. Differential hemodynamic response in affective circuitry with aging: An fmri study of novelty, valence, and arousal. J. Cogn. Neurosci. 23, 1027–1041 (2011).

62 Weierich, M. R., Wright, C. I., Negreira, A., Dickerson, B. C. & Barrett, L. F. Novelty as a dimension in the affective brain. Neuroimage 49, 2871–2878 (2010).

63 Damasio, A. R. Descartes’ error: Emotion, reason, and the human brain. (Putnam, 1994).

64 Wiens, S., Mezzacappa, E. S. & Katkin, E. S. Heartbeat detection and the experience of emotions. Cognition and Emotion 14, 417–427 (2000).

65 Barrett, L. F., Quigley, K. S., Bliss-Moreau, E. & Aronson, K. R. Interoceptive sensitivity and self-reports of emotional experience. J. Pers. Soc. Psychol. 87, 684–697 (2004).

66 Dunn, B. D. et al. Listening to your heart: How interoception shapes emotion experience and intuitive decision making. Psychol. Sci. 21, 1835–1844 (2010).

67 Whitehead, W. E., Drescher, V. M., Heiman, P. & Blackwell, B. Relation of heart rate control to heartbeat perception. Biofeedback and Self-Regulation 2, 371–392 (1977).

68 Schandry, R. Heart beat perception and emotional experience. Psychophysiology 18, 483–488 (1981).

69 Kleckner, I. R., Wormwood, J. B., Simmons, W. K., Barrett, L. F. & Quigley, K. S. Methodological recommendations for a heartbeat detection-based measure of interoceptive sensitivity. Psychophysiology 52, 1432–1440 (2015).

70 Khalsa, S. S., Rudrauf, D., Feinstein, J. S. & Tranel, D. The pathways of interoceptive awareness. Nat. Neurosci. 12, 1494–1496 (2009).

71 Lindquist, K. A. & Barrett, L. F. A functional architecture of the human brain: Emerging insights from the science of emotion. Trends in cognitive sciences 16, 533–540 (2012).

72 van den Heuvel, M. P. & Sporns, O. Rich-club organization of the human connectome. J. Neurosci. 31, 15775–15786 (2011).

73 McIntosh, A. R. Contexts and catalysts: A resolution of the localization and integration of function in the brain. Neuroinformatics 2, 175–182 (2004).

74 Beissner, F., Schumann, A., Brunn, F., Eisentrager, D. & Bar, K. J. Advances in functional magnetic resonance imaging of the human brainstem. Neuroimage 86, 91–98 (2014).

75 Bar, K. J. et al. Functional connectivity and network analysis of midbrain and brainstem nuclei. Neuroimage 134, 53–63 (2016).

76 Edlow, B. L., McNab, J. A., Witzel, T. & Kinney, H. C. The structural connectome of the human central homeostatic network. Brain Connect 6, 187–200 (2016).

77 Dum, R. P., Levinthal, D. J. & Strick, P. L. Motor, cognitive, and affective areas of the cerebral cortex influence the adrenal medulla. Proc. Natl. Acad. Sci. U. S. A. 113, 9922–9927 (2016).

78 Blessing, W. W. & Benarroch, E. E. in The human nervous system (eds Mai J.K. & Paxinos G.) 1058–1073 (Academic Press, 2012).

79 Simmons, W. K. et al. Keeping the body in mind: Insula functional organization and functional connectivity integrate interoceptive, exteroceptive, and emotional awareness. Hum. Brain Mapp. 000 (2012).

80 Margulies, D. S. et al. Mapping the functional connectivity of anterior cingulate cortex. Neuroimage 37, 579–588 (2007).

81 Bickart, K. C., Dickerson, B. C. & Barrett, L. F. The amygdala as a hub in brain networks that support social life. Neuropsychologia 63, 235–248 (2014).

82 Smith, D. V., Clithero, J. A., Boltuck, S. E. & Huettel, S. A. Functional connectivity with ventromedial prefrontal cortex reflects subjective value for social rewards. Soc. Cogn. Affect. Neurosci. 9, 2017–2025 (2014).

83 van den Heuvel, M. P., Kahn, R. S., Goni, J. & Sporns, O. High-cost, high-capacity backbone for global brain communication. Proc. Natl. Acad. Sci. U. S. A. 109, 11372–11377 (2012).

84 van den Heuvel, M. P. & Sporns, O. An anatomical substrate for integration among functional networks in human cortex. J. Neurosci. 33, 14489–14500 (2013).

85 Mesulam, M. M. From sensation to cognition. Brain 121, 1013–1052 (1998).

86 Zold, C. L. & Hussain Shuler, M. G. Theta oscillations in visual cortex emerge with experience to convey expected reward time and experienced reward rate. J. Neurosci. 35, 9603–9614 (2015).

87 Satpute, A. B. et al. Involvement of sensory regions in affective experience: A meta-analysis. Front. Psychol. 6, 1860 (2015).

88 Vuilleumier, P. How brains beware: Neural mechanisms of emotional attention. Trends in cognitive sciences 9, 585–594 (2005).

89 Sun, F. W. et al. Youthful brains in older adults: Preserved neuroanatomy in the default mode and salience networks contributes to youthful memory in superaging. J. Neurosci. 36, 9659–9668 (2016).

90 Barrett, L. F. & Bliss-Moreau, E. Affect as a psychological primitive. Advances in Experimental Social Psychology 41, 167–218 (2009).

91 Barrett, L. F. How emotions are made: The secret life the brain., (Houghton-Mifflin-Harcourt, 2017).

92 Barrett, L. F. & Russell, J. A. Structure of current affect: Controversies and emerging consensus. Current Directions in Psychological Science 8, 10–14 (1999).

93 Kuppens, P., Tuerlinckx, F., Russell, J. A. & Barrett, L. F. The relation between valence and arousal in subjective experience. Psychol Bull 139, 917–940 (2013).

94 Damasio, A. R. The feeling of what happens: Body and emotion in the making of consciousness. (Houghton Mifflin Harcourt, 1999).

95 Dreyfus, G. & Thompson, E. in The cambridge handbook of consciousness (eds Philip David Zelazo, Morris Moscovitch, & Evan Thompson) 89–114 (Cambridge University Press, 2007).

96 Edelman, G. M. & Tononi, G. A universe of consciousness: How matter becomes imagination. (Basic books, 2000).

97 James, W. The principles of psychology. Vol. 1 (Dover, 1890/2007).

98 Searle, J. R. The rediscovery of the mind. (MIT press, 1992).

99 Searle, J. R. Mind: A brief introduction. (Oxford University Press, 2004).

100 Wundt, W. Outlines of psychology. (Wilhelm Engelmann, 1897).

101 Fodor, J. A. The modularity of mind: An essay on faculty psychology. (MIT press, 1983).

102 Li, D., Christ, S. E. & Cowan, N. Domain-general and domain-specific functional networks in working memory. Neuroimage 102 Pt 2, 646–656 (2014).

103 Fuster, J. M. (Cell Press, 2000).

104 Kayser, C., Petkov, C. I., Augath, M. & Logothetis, N. K. Functional imaging reveals visual modulation of specific fields in auditory cortex. J. Neurosci. 27, 1824–1835 (2007).

105 Liang, M., Mouraux, A., Hu, L. & Iannetti, G. D. Primary sensory cortices contain distinguishable spatial patterns of activity for each sense. Nature communications 4, 1979 (2013).

106 Edelman, G. M. & Gally, J. A. Degeneracy and complexity in biological systems. Proceedings of the National Academy of Sciences 98, 13763–13768 (2001).

107 Tong, Y. et al. Systemic low-frequency oscillations in bold signal vary with tissue type. Front. Neurosci. 10, 313 (2016).

108 Satpute, A. B. et al. Identification of discrete functional subregions of the human periaqueductal gray. Proc. Natl. Acad. Sci. U. S. A. 110, 17101–17106 (2013).

109 Murphy, K., Birn, R. M., Handwerker, D. A., Jones, T. B. & Bandettini, P. A. The impact of global signal regression on resting state correlations: Are anti-correlated networks introduced? Neuroimage 44, 893–905 (2009).

110 Raz, G. et al. Functional connectivity dynamics during film viewing reveal common networks for different emotional experiences. Cogn. Affect. Behav. Neurosci. (2016).

111 Morrison, S. F. Differential control of sympathetic outflow. Am. J. Physiol. Regul. Integr. Comp. Physiol. 281, R683–698 (2001).

112 Buckner, R. L. The serendipitous discovery of the brain’s default network. Neuroimage 62, 1137–1145 (2012).

113 wMesulam, M. The evolving landscape of human cortical connectivity: Facts and inferences. Neuroimage 62, 2182–2189 (2012).

114 Hassabis, D. & Maguire, E. A. The construction system of the brain. Philos. Trans. R. Soc. Lond. B Biol. Sci. 364, 1263–1271 (2009).

115 Power, J. D. et al. Functional network organization of the human brain. Neuron 72, 665–678 (2011).

116 Menon, V. & Uddin, L. Q. Saliency, switching, attention and control: A network model of insula function. Brain structure & function 214, 655–667 (2010).

117 Li, S. S. & McNally, G. P. The conditions that promote fear learning: Prediction error and pavlovian fear conditioning. Neurobiol. Learn. Mem. 108, 14–21 (2014).

118 Barrett, L. F. How emotions are made: The new science of the mind and brain. (Houghton-Mifflin-Harcourt, in press).

119 Guidi, M., Huber, L., Lampe, L., Gauthier, C. J. & Moller, H. E. Lamina-dependent calibrated bold response in human primary motor cortex. Neuroimage 141, 250–261 (2016).

120 Crossley, N. A. et al. The hubs of the human connectome are generally implicated in the anatomy of brain disorders. Brain 137, 2382–2395 (2014).

121 Goodkind, M. et al. Identification of a common neurobiological substrate for mental illness. JAMA psychiatry 72, 305–315 (2015).

122 Menon, V. Large-scale brain networks and psychopathology: A unifying triple network model. Trends in cognitive sciences 15, 483–506 (2011).

123 Harshaw, C. Interoceptive dysfunction: Toward an integrated framework for understanding somatic and affective disturbance in depression. Psychol. Bull. 141, 311–363 (2015).

124 Paulus, M. P. & Stein, M. B. Interoception in anxiety and depression. Brain Structure Function 214, 451–463 (2010).

125 Naqvi, N. H. & Bechara, A. The insula and drug addiction: An interoceptive view of pleasure, urges, and decision-making. Brain Structure Function 214, 435–450 (2010).

126 Farmer, M. A., Baliki, M. N. & Apkarian, A. V. A dynamic network perspective of chronic pain. Neurosci. Lett. 520, 197–203 (2012).

127 Mayer, E. A. Gut feelings: The emerging biology of gut-brain communication. Nat. Rev. Neurosci. 12, 453–466 (2011).

128 Radley, J., Morilak, D., Viau, V. & Campeau, S. Chronic stress and brain plasticity: Mechanisms underlying adaptive and maladaptive changes and implications for stress-related cns disorders. Neurosci. Biobehav. Rev. 58, 79–91 (2015).

129 Gianaros, P. J. & Wager, T. D. Brain-body pathways linking psychological stress and physical health. Curr. Dir. Psychol. Sci. 24, 313–321 (2015).

130 Gefen, T. et al. Morphometric and histologic substrates of cingulate integrity in elders with exceptional memory capacity. J. Neurosci. 35, 1781–1791 (2015).

131 Rogalski, E. J. et al. Youthful memory capacity in old brains: Anatomic and genetic clues from the northwestern superaging project. J. Cogn. Neurosci. 25, 29–36 (2013).

132 Angevaren, M., Aufdemkampe, G., Verhaar, H. J., Aleman, A. & Vanhees, L. Physical activity and enhanced fitness to improve cognitive function in older people without known cognitive impairment. Cochrane Database Syst Rev, CD005381 (2008).

133 Farb, N. et al. Interoception, contemplative practice, and health. Front. Psychol. 6, 763 (2015).

134 Tang, Y. Y., Holzel, B. K. & Posner, M. I. The neuroscience of mindfulness meditation. Nat. Rev. Neurosci. 16, 213–225 (2015).

135 Thombs, B. D. et al. Prevalence of depression in survivors of acute myocardial infarction. J. Gen. Intern. Med. 21, 30–38 (2006).

136 Moreno-Smith, M., Lutgendorf, S. K. & Sood, A. K. Impact of stress on cancer metastasis. Future Oncol. 6, 1863–1881 (2010).

137 Kolodny, A. et al. The prescription opioid and heroin crisis: A public health approach to an epidemic of addiction. Annu. Rev. Public Health 36, 559–574 (2015).

138 Turecki, G. & Brent, D. A. Suicide and suicidal behaviour. Lancet 387, 1227–1239 (2016).

139 Bohus, B. et al. The neurobiology of the central nucleus of the amygdala in relation to neuroendocrine and autonomic outflow. Prog. Brain Res. 107, 447–460 (1996).

140 Mufson, E. J., Mesulam, M. M. & Pandya, D. N. Insular interconnections with the amygdala in rhesus monkey. Neuroscience 6, 1231–1248 (1981).

141 Mufson, E. J. & Mesulam, M. M. Insula of the old world monkey. Ii: Afferent cortical input and comments on the claustrum. J. Comp. Neurol. 212, 23–37 (1982).

142 Ghashghaei, H. T., Hilgetag, C. C. & Barbas, H. Sequence of information processing for emotions based on the anatomic dialogue between prefrontal cortex and amygdala. Neuroimage 34, 905–923 (2007).

143 Morecraft, R. J. et al. Cytoarchitecture and cortical connections of the anterior cingulate and adjacent somatomotor fields in the rhesus monkey. Brain Res. Bull. 87, 457–497 (2012).

144 Mesulam, M. M. & Mufson, E. J. Insula of the old world monkey. Iii: Efferent cortical output and comments on function. J. Comp. Neurol. 212, 38–52 (1982).

145 Cavdar, S. et al. The afferent connections of the posterior hypothalamic nucleus in the rat using horseradish peroxidase. J. Anat. 198, 463–472 (2001).

146 An, X., Bandler, R., Ongur, D. & Price, J. L. Prefrontal cortical projections to longitudinal columns in the midbrain periaqueductal gray in macaque monkeys. J. Comp. Neurol. 401, 455–479 (1998).

147 Saper, C. B. Reciprocal parabrachial-cortical connections in the rat. Brain Res. 242, 33–40 (1982).

148 Saper, C. B. Convergence of autonomic and limbic connections in the insular cortex of the rat. J. Comp. Neurol. 210, 163–173 (1982).

149 Fudge, J. L., Breitbart, M. A., Danish, M. & Pannoni, V. Insular and gustatory inputs to the caudal ventral striatum in primates. J. Comp. Neurol. 490, 101–118 (2005).

150 Carmichael, S. T. & Price, J. L. Connectional networks within the orbital and medial prefrontal cortex of macaque monkeys. J. Comp. Neurol. 371, 179–207 (1996).

151 Aggleton, J. P., Burton, M. J. & Passingham, R. E. Cortical and subcortical afferents to the amygdala of the rhesus monkey (macaca mulatta). Brain Res. 190, 347–368 (1980).

152 Stefanacci, L. & Amaral, D. G. Some observations on cortical inputs to the macaque monkey amygdala: An anterograde tracing study. J. Comp. Neurol. 451, 301–323 (2002).

153 Chikama, M., McFarland, N. R., Amaral, D. G. & Haber, S. N. Insular cortical projections to functional regions of the striatum correlate with cortical cytoarchitectonic organization in the primate. J. Neurosci. 17, 9686–9705 (1997).

154 Chiba, T., Kayahara, T. & Nakano, K. Efferent projections of infralimbic and prelimbic areas of the medial prefrontal cortex in the japanese monkey, macaca fuscata. Brain Res. 888, 83–101 (2001).

155 Vogt, B. A. & Pandya, D. N. Cingulate cortex of the rhesus monkey: Ii. Cortical afferents. J. Comp. Neurol. 262, 271–289 (1987).

156 Rempel-Clower, N. L. & Barbas, H. Topographic organization of connections between the hypothalamus and prefrontal cortex in the rhesus monkey. J. Comp. Neurol. 398, 393–419 (1998).

157 Freedman, L. J., Insel, T. R. & Smith, Y. Subcortical projections of area 25 (subgenual cortex) of the macaque monkey. J. Comp. Neurol. 421, 172–188 (2000).

158 Terreberry, R. R. & Neafsey, E. J. Rat medial frontal cortex: A visceral motor region with a direct projection to the solitary nucleus. Brain Res. 278, 245–249 (1983).

159 van der Kooy, D., McGinty, J. F., Koda, L. Y., Gerfen, C. R. & Bloom, F. E. Visceral cortex: A direct connection from prefrontal cortex to the solitary nucleus in rat. Neurosci. Lett. 33, 123–127 (1982).

160 Room, P., Russchen, F. T., Groenewegen, H. J. & Lohman, A. H. Efferent connections of the prelimbic (area 32) and the infralimbic (area 25) cortices: An anterograde tracing study in the cat. J. Comp. Neurol. 242, 40–55 (1985).

161 Pandya, D. N., Van Hoesen, G. W. & Mesulam, M. M. Efferent connections of the cingulate gyrus in the rhesus monkey. Exp. Brain Res. 42, 319–330 (1981).

162 Vogt, B. A. & Palomero-Gallagher, N. in The human central nervous system (eds J.K. Mai & G. Paxinos) Ch. 25, 943–987 (Elsevier, 2012).

163 Haber, S. N., Kim, K. S., Mailly, P. & Calzavara, R. Reward-related cortical inputs define a large striatal region in primates that interface with associative cortical connections, providing a substrate for incentive-based learning. J. Neurosci. 26, 8368–8376 (2006).

164 Price, J. L. & Amaral, D. G. An autoradiographic study of the projections of the central nucleus of the monkey amygdala. J. Neurosci. 1, 1242–1259 (1981).

165 Fudge, J. L., Kunishio, K., Walsh, P., Richard, C. & Haber, S. N. Amygdaloid projections to ventromedial striatal subterritories in the primate. Neuroscience 110, 257–275 (2002).

166 Morecraft, R. J. & Van Hoesen, G. W. Convergence of limbic input to the cingulate motor cortex in the rhesus monkey. Brain Res. Bull. 45, 209–232 (1998).

167 Saleem, K. S., Kondo, H. & Price, J. L. Complementary circuits connecting the orbital and medial prefrontal networks with the temporal, insular, and opercular cortex in the macaque monkey. J. Comp. Neurol. 506, 659–693 (2008).

168 Barbas, H., Ghashghaei, H., Dombrowski, S. M. & Rempel-Clower, N. L. Medial prefrontal cortices are unified by common connections with superior temporal cortices and distinguished by input from memory-related areas in the rhesus monkey. J. Comp. Neurol. 410, 343–367 (1999).

169 Ongur, D., Ferry, A. T. & Price, J. L. Architectonic subdivision of the human orbital and medial prefrontal cortex. J. Comp. Neurol. 460, 425–449 (2003).

170 Ghaziri, J. et al. The corticocortical structural connectivity of the human insula. Cereb. Cortex (2015).

171 Wiech, K., Jbabdi, S., Lin, C. S., Andersson, J. & Tracey, I. Differential structural and resting state connectivity between insular subdivisions and other pain-related brain regions. Pain 155, 2047–2055 (2014).

172 Gianaros, P. J. & Sheu, L. K. A review of neuroimaging studies of stressor-evoked blood pressure reactivity: Emerging evidence for a brain-body pathway to coronary heart disease risk. Neuroimage 47, 922–936 (2009).

173 Kurth, F., Zilles, K., Fox, P. T., Laird, A. R. & Eickhoff, S. B. A link between the systems: Functional differentiation and integration within the human insula revealed by meta-analysis. Brain structure & function 214, 519–534 (2010).

174 Gianaros, P. J. et al. An inflammatory pathway links atherosclerotic cardiovascular disease risk to neural activity evoked by the cognitive regulation of emotion. Biol. Psychiatry 75, 738–745 (2014).

175 Wager, T. D. et al. Brain mediators of cardiovascular responses to social threat: Part i: Reciprocal dorsal and ventral sub-regions of the medial prefrontal cortex and heart-rate reactivity. Neuroimage 47, 821–835 (2009).

176 Harper, R. M. et al. Fmri responses to cold pressor challenges in control and obstructive sleep apnea subjects. J Appl Physiol (1985) 94, 1583–1595 (2003).

177 Gianaros, P. J. et al. Individual differences in stressor-evoked blood pressure reactivity vary with activation, volume, and functional connectivity of the amygdala. J. Neurosci. 28, 990–999 (2008).

178 Zilles, K. & Amunts, K. in The human central nervous system (eds J.K. Mai & G. Paxinos) Ch. 23, 836–895 (Elsevier, 2012).

179 Petrides, M. & Pandya, D. N. in The human central nervous system (eds J.K. Mai & G. Paxinos) Ch. 26, 988–1011 (Elsevier, 2012).

180 Barbas, H. Specialized elements of orbitofrontal cortex in primates. Ann. N. Y. Acad. Sci. 1121, 10–32 (2007).

181 Mesulam, M. M. & Mufson, E. J. Insula of the old world monkey. I. Architectonics in the insulo-orbito-temporal component of the paralimbic brain. J. Comp. Neurol. 212, 1–22 (1982).

182 Yarkoni, T. Big correlations in little studies: Inflated fmri correlations reflect low statistical power-commentary on vul et al. (2009). Perspect. Psychol. Sci. 4, 294–298 (2009).

183 Cronbach, L. J., Rajaratnam, N. & Gleser, G. C. Theory of generalizability: A liberalization of reliability theory. British Journal of Statistical Psychology 16, 137–163 (1963).

184 Wilson-Mendenhall, C. D., Barrett, L. F., Simmons, W. K. & Barsalou, L. W. Grounding emotion in situated conceptualization. Neuropsychologia 49, 1105–1127 (2011).

185 Wilson-Mendenhall, C. D., Barrett, L. F. & Barsalou, L. W. Variety in emotional life: Within-category typicality of emotional experiences is associated with neural activity in large-scale brain networks. Soc. Cogn. Affect. Neurosci. 10, 62–71 (2015).

186 Kober, H. et al. Functional grouping and cortical-subcortical interactions in emotion: A meta-analysis of neuroimaging studies. Neuroimage 42, 998–1031 (2008).

187 Skerry, A. E. & Saxe, R. Neural representations of emotion are organized around abstract event features. Curr. Biol. 25, 1945–1954 (2015).

188 Clithero, J. A. & Rangel, A. Informatic parcellation of the network involved in the computation of subjective value. Soc. Cogn. Affect. Neurosci. 9, 1289–1302 (2014).

189 Bickart, K. C., Hollenbeck, M. C., Barrett, L. F. & Dickerson, B. C. Intrinsic amygdala-cortical functional connectivity predicts social network size in humans. J. Neurosci. 32, 14729–14741 (2012).

190 Baliki, M. N., Mansour, A. R., Baria, A. T. & Apkarian, A. V. Functional reorganization of the default mode network across chronic pain conditions. PLoS One 9, e106133 (2014).

191 Schurz, M., Radua, J., Aichhorn, M., Richlan, F. & Perner, J. Fractionating theory of mind: A meta-analysis of functional brain imaging studies. Neurosci. Biobehav. Rev. 42, 9–34 (2014).

192 Morelli, S. A. & Lieberman, M. D. The role of automaticity and attention in neural processes underlying empathy for happiness, sadness, and anxiety. Front. Hum. Neurosci. 7, 160 (2013).

193 Chiong, W. et al. The salience network causally influences default mode network activity during moral reasoning. Brain 136, 1929–1941 (2013).

194 Liu, X., Hairston, J., Schrier, M. & Fan, J. Common and distinct networks underlying reward valence and processing stages: A meta-analysis of functional neuroimaging studies. Neurosci. Biobehav. Rev. 35, 1219–1236 (2011).

195 Engelmann, J. M. et al. Neural substrates of smoking cue reactivity: A meta-analysis of fmri studies. Neuroimage 60, 252–262 (2012).

196 Schacter, D. L. & Addis, D. R. The cognitive neuroscience of constructive memory: Remembering the past and imagining the future. Philos. Trans. R. Soc. Lond. B Biol. Sci. 362, 773–786 (2007).

197 Bar, M., Aminoff, E., Mason, M. & Fenske, M. The units of thought. Hippocampus 17, 420–428 (2007).

198 Fernandino, L. et al. Concept representation reflects multimodal abstraction: A framework for embodied semantics. Cereb. Cortex 26, 2018–2034 (2016).

199 Touroutoglou, A., Lindquist, K. A., Dickerson, B. C. & Barrett, L. F. Intrinsic connectivity in the human brain does not reveal networks for 'basic' emotions. Soc. Cogn. Affect. Neurosci. 10, 1257–1265 (2015).

200 Dhanjal, N. S. & Wise, R. J. Frontoparietal cognitive control of verbal memory recall in alzheimer's disease. Ann. Neurol. 76, 241–251 (2014).

201 Dosenbach, N. U. et al. Distinct brain networks for adaptive and stable task control in humans. Proc. Natl. Acad. Sci. U. S. A. 104, 11073–11078 (2007).

202 Ansell, E. B., Rando, K., Tuit, K., Guarnaccia, J. & Sinha, R. Cumulative adversity and smaller gray matter volume in medial prefrontal, anterior cingulate, and insula regions. Biol. Psychiatry 72, 57–64 (2012).

203 Caseras, X. et al. Anatomical and functional overlap within the insula and anterior cingulate cortex during interoception and phobic symptom provocation. Hum. Brain Mapp. 34, 1220–1229 (2013).

204 Wolf, D. H. et al. Striatal intrinsic reinforcement signals during recognition memory: Relationship to response bias and dysregulation in schizophrenia. Front. Behav. Neurosci. 5, 81 (2011).

205 Grady, C. L., Luk, G., Craik, F. I. & Bialystok, E. Brain network activity in monolingual and bilingual older adults. Neuropsychologia 66, 170–181 (2015).

206 Derbyshire, S. W., Whalley, M. G., Stenger, V. A. & Oakley, D. A. Cerebral activation during hypnotically induced and imagined pain. Neuroimage 23, 392–401 (2004).

207 Feldstein Ewing, S. W., Filbey, F. M., Sabbineni, A., Chandler, L. D. & Hutchison, K. E. How psychosocial alcohol interventions work: A preliminary look at what fmri can tell us. Alcohol. Clin. Exp. Res. 35, 643–651 (2011).

208 Singer, T. et al. Empathy for pain involves the affective but not sensory components of pain. Science 303, 1157–1162 (2004).

209 Kirk, U., Downar, J. & Montague, P. R. Interoception drives increased rational decision-making in meditators playing the ultimatum game. Front. Neurosci. 5, 49 (2011).

210 FitzGerald, T. H., Schwartenbeck, P. & Dolan, R. J. Reward-related activity in ventral striatum is action contingent and modulated by behavioral relevance. J. Neurosci. 34, 1271–1279 (2014).

211 Fernandino, L., Humphries, C. J., Conant, L., Seidenberg, M. S. & Binder, J. R. Heteromodal cortical areas encode sensory-motor features of word meaning. J. Neuroscience (in press).

212 Hermans, E. J. et al. Stress-related noradrenergic activity prompts large-scale neural network reconfiguration. Science 334, 1151–1153 (2011).

213 Clark-Polner, E., Wager, T. D., Satpute, A. B. & Barrett, L. F. in The handbook of emotion (eds L.F. Barrett, M. Lewis, & J.M. Haviland-Jones) (Guilford, in press).

214 Buckner, R. L., Krienen, F. M., Castellanos, A., Diaz, J. C. & Yeo, B. T. The organization of the human cerebellum estimated by intrinsic functional connectivity. J. Neurophysiol. 106, 2322–2345 (2011).

215 Touroutoglou, A., Andreano, J. M., Barrett, L. F. & Dickerson, B. C. Brain network connectivity-behavioral relationships exhibit trait-like properties: Evidence from hippocampal connectivity and memory. Hippocampus (2015).

216 Kramer, J. H. et al. Nih examiner: Conceptualization and development of an executive function battery. J. Int. Neuropsychol. Soc. 20, 11–19 (2014).

217 Bradley, M. M. & Lang, P. J. Measuring emotion: The self-assessment manikin and the semantic differential. J. Behav. Ther. Exp. Psychiatry 25, 49–59 (1994).

218 Jo, H. J. et al. Effective preprocessing procedures virtually eliminate distance-dependent motion artifacts in resting state fmri. J Appl Math 2013 (2013).

219 Power, J. D., Barnes, K. A., Snyder, A. Z., Schlaggar, B. L. & Petersen, S. E. Spurious but systematic correlations in functional connectivity mri networks arise from subject motion. Neuroimage 59, 2142–2154 (2012).

220 Tucholka, A., Fritsch, V., Poline, J. B. & Thirion, B. An empirical comparison of surface-based and volume-based group studies in neuroimaging. Neuroimage 63, 1443–1453 (2012).

221 Boucsein, W. et al. Publication recommendations for electrodermal measurements. Psychophysiology 49, 1017–1034 (2012).

222 Schell, A. M., Dawson, M. E. & Filion, D. L. Psychophysiological correlates of electrodermal lability. Psychophysiology 25, 619–632 (1988).

223 Xia, C., Touroutoglou, A., Quigley, K. S., Barrett, L. F. & Dickerson, B. C. Task-evoked skin conductance responses modulates the influence of salience network connectivity in predicting subjective experience of arousal. J. Cogn. Neurosci. (in press).

224 Fredrikson, M. et al. Functional neuroanatomical correlates of electrodermal activity: A positron emission tomographic study. Psychophysiology 35, 179–185 (1998).

225 Nieuwenhuys, R., Voogd, J. & Huijzen, C. v. in The human central nervous system Ch. 10, 289–336 (Springer, 2008).

226 Pisotta, I. & Molinari, M. Cerebellar contribution to feedforward control of locomotion. Front. Hum. Neurosci. 8, 475 (2014).

227 Wolpert, D. M. & Kawato, M. Multiple paired forward and inverse models for motor control. Neural Netw 11, 1317–1329 (1998).

228 Nieuwenhuys, R., Voogd, J. & Van Huijzen, C. in The human central nervous system Ch. 13, 401–426 (Springer, 2008).

229 Nieuwenhuys, R., Voogd, J. & Van Huijzen, C. in The human central nervous system Ch. 12, 361–400 (Springer, 2008).

230 Haber, S. N. & Behrens, T. E. The neural network underlying incentive-based learning: Implications for interpreting circuit disruptions in psychiatric disorders. Neuron 83, 1019–1039 (2014).

231 Swanson, L. W. Cerebral hemisphere regulation of motivated behavior. Brain Res. 886, 113–164 (2000).

232 Carrive, P. in The human nervous system (eds Mai J.K. & Paxinos G.) 367–400 (Academic Press, 2012).

233 Paxinos, G., Feng-Xu, H., Sengul, G. & Watson, C. in The human nervous system (eds J. K. Mai & G. Paxinos) Ch. 8, 260–327 (Springer, 2012).

234 Dawson, M. E., Schell, A. M. & Filion, D. L. in Handbook of psychophysiology (eds J. T. Cacioppo, L. G. Tassinary, & G. G. Berntson) 159–181 (Cambridge University Press, 2007).

235 Peugh, J. L. A practical guide to multilevel modeling. J. Sch. Psychol. 48, 85–112 (2010).

236 Bradley, M. M. & Lang, P. J. in Handbook of emotion elicitation and assessment. Series in affective science Vol. viii (eds J. A. Coan & J. J. B. Allen) (Oxford University Press, 2007).

237 Risold, P. Y., Thompson, R. H. & Swanson, L. W. The structural organization of connections between hypothalamus and cerebral cortex. Brain Res. Brain Res. Rev. 24, 197–254 (1997).

238 Parvizi, J., Van Hoesen, G. W., Buckwalter, J. & Damasio, A. Neural connections of the posteromedial cortex in the macaque. Proc. Natl. Acad. Sci. U. S. A. 103, 1563–1568 (2006).

239 Dujardin, E. & Jurgens, U. Afferents of vocalization-controlling periaqueductal regions in the squirrel monkey. Brain Res. 1034, 114–131 (2005).

240 Saper, C. B. in The human nervous system (eds J. K. Mai & G. Paxinos) Ch. 16, 548–583 (Elsevier, 2012).

241 Bickart, K. C. et al. Atrophy in distinct corticolimbic networks in frontotemporal dementia relates to social impairments measured using the social impairment rating scale. J. Neurol. Neurosurg. Psychiatry 85, 438–448 (2014).

